# An ancient monoaminergic signaling system coordinates contractility in a nerveless sponge

**DOI:** 10.64898/2026.02.14.704838

**Authors:** Rong Xuan Zang, Nawaphat Malaiwong, Ling Wang, Jamie D. Maziarz, Kejue Jia, Bernhard Drotleff, Frank Stein, Mah Noor, C. Jackson Roberts, Mandy Rettel, Jennifer J. Schwarz, Aissam Ikmi, Shigeki Watanabe, Robert Prevedel, Michael P. O’Donnell, Jacob M. Musser

## Abstract

Chemical neurotransmission was a key animal innovation, enabling multicellular coordination of physiology and locomotion. Sponges are early-diverging animals that lack neurons and muscles, yet still coordinate contraction and relaxation of their filter-feeding water canals. Here, we show that *Spongilla lacustris* synthesizes the monoamines tryptamine, phenethylamine, and tyramine to elicit distinct canal behaviors. We identify previously uncharacterized decarboxylases and vesicular transporters coexpressed in secretory neuroid and metabolic cells. Using phosphoproteomics and label-free 3D imaging, we show that tryptamine activates GPCR signaling and Rho GTPases, remodeling adhesion and actomyosin networks in contractile canal epithelia to drive localized constrictions and whole-body deflations. Together, these findings define an ancestral monoaminergic system linking secretory and contractile cell types that predate neurons and was later elaborated for neuromodulation of synaptic transmission.

---

Nervous systems coordinate behavior and physiology through chemical transmitters, yet how transmitter systems first evolved and were produced to control specific cellular activities remains unclear (*1–5*). In bilaterian animals, transmitter signaling typically occurs in the nervous system, where small molecules and neuropeptides are released at synapses or act extrasynaptically through volume transmission to regulate contractility, homeostasis, and other cellular behaviors (*6*). This may have been preceded by simpler circuits in which sensory secretory cells broadcast neuropeptides, amines, or other diffusible molecules to nearby effector cells that converted these cues into behavioral responses (*2*, *5*). Determining whether such circuits existed before neurons and synapses is central to reconstructing early animal communication and nervous system evolution.

Sponges diverged early in animal evolution and lack neurons and muscles, but display coordinated contractile behaviors suggestive of chemical transmission (Fig. 1 A-C) (*7–9*). In the freshwater demosponge *Spongilla lacustris*, these behaviors modulate a branched water canal system for filter feeding (*9*, *10*). Water enters through ostia pores in an outer epithelial tent into incurrent canals, passes through choanocyte chambers for food uptake, then exits through excurrent canals and the osculum (Fig. 1C). Contractile pinacocytes line the canal system and regulate canal shape and tension within a mesohyl tissue populated by stem cells and several motile cell types. *S. lacustris* continuously alters its canal geometry, including localized peristaltic twitches that tune canal volume and stereotypic whole-body deflations that transiently close and flush the entire canal system. During deflations, ostia close, incurrent canals collapse, and water is expelled through expanded excurrent canals. Recovery occurs when the sponge reinflates its incurrent system returning canals to a resting configuration (Fig. 1C) (*9*).

**Fig. 1.**
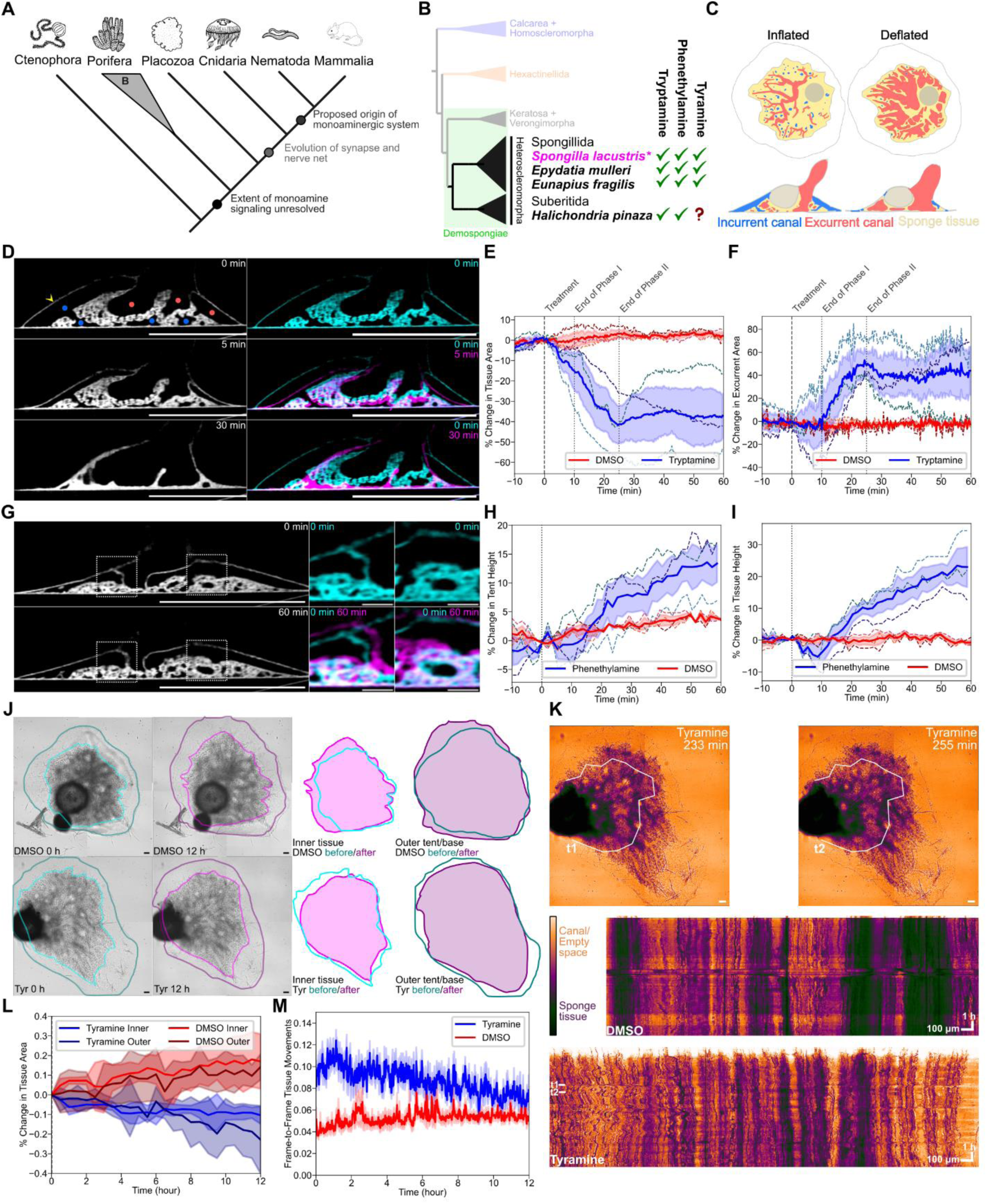
Distinct and conserved behavioral responses to monoamines in demosponges. **(A)** Animal phylogeny depicting major transitions in nervous system evolution. **(B)** Phylogenetic relationships among major sponge lineages and summary of behavioral responses to monoamines. **(C)** Schematic of sponge canal system in inflated and deflated states. **(D)** Representative OCM cross-sections of *S. lacustris* before and after tryptamine treatment (37.5 µM). Cyan and magenta indicate pretreatment (t = 0) and post-treatment time points (t = 5 min and t = 30 min). Markers: yellow arrowhead, outer epithelial tent; blue dots, incurrent canals; red dots, excurrent canals. Scale bar, 1 mm. **(E)** Percent change in tissue area normalized to pretreatment following tryptamine or DMSO treatment. Colored dashed lines indicate biological replicates (n = 3). Shading indicates standard error of the mean (SEM). The vertical dashed line indicates addition of 37.5 μM tryptamine and dotted line marks the transitions between behavioral phases. **(F)** Percent change in excurrent canal area normalized to pretreatment following tryptamine or DMSO treatment. Colored dashed lines indicate biological replicates (n = 3). Shading indicates SEM. Vertical dashed and dotted lines are as in (E). **(G)** Representative OCM cross-sections of *S. lacustris* before and after phenethylamine treatment (37.5 µM). Scale bar (main), 1 mm. Scale bar (inset), 100 µm. **(H)** Percent change in epithelial tent height normalized to pretreatment following application of phenethylamine or DMSO. Colored dashed lines indicate biological replicates (n = 3). Shading indicates SEM. **(I)** Percent change in tissue height normalized to pretreatment following phenethylamine or DMSO treatment. Colored dashed lines indicate biological replicates (n = 3). Shading indicates SEM. **(J)** Representative brightfield images of *S. lacustris* treated with DMSO or tyramine (50 µM) at 0 and 12 h, illustrating gradual centripetal tissue retraction. Scale bar, 100 µm. **(K)** Canal activity visualized by line-scan kymographs from representative DMSO- and tyramine-treated animals. Top panels show the kymograph line position at two time points (t1, t2). Bottom panels show kymographs acquired over 12 hours. Scale bar, 100 µm. **(L)** Percent change in inner tissue and outer epithelial attachment area relative to pretreatment over 12 h following tyramine (n = 7) or DMSO treatment (n = 7). **(M)** Canal activity and tissue movements quantified as pixel dissimilarity (Pearson’s distance, see Materials and Methods) over time following tyramine (n = 7) or DMSO treatment (n = 6). Dissimilarity reflects frame-to-frame intensity changes along canal and tissue structures.

Several decades of work have begun to identify signaling and effector mechanisms underlying sponge deflations (*9*, *11–15*). In particular, nitric oxide (NO) acts as a rapidly diffusing gas transmitter that drives relaxation of contractile pinacocytes during deflation, similar to its role in vertebrate vasculature. Sponges also encode extensive machinery for regulated secretion, and homologs of synaptic and secretory genes are expressed in specific cell types, including neuroid cells associated with choanocyte chambers, as well as digestive, contractile, and metabolic cell types (*10*). Consistent with a role for other transmitters, different molecules have been reported to induce or alter sponge deflations. For instance, in marine sponges the monoamines serotonin and epinephrine can alter canal behavior (*11*), and dopamine and several simple monoamines produced by decarboxylation of amino acids have been shown to affect larval phototaxis (*16*). However, canonical enzymes and vesicular transporters associated with monoamine synthesis and storage in bilaterians appear to be absent in sponges (*17*). As a result, it remains unclear whether early animals were capable of endogenous synthesis, storage, and regulated release of monoamines or other transmitters, and how such signaling systems might be organized spatially to control specific behaviors.

Here we uncover a monoaminergic circuit in *S. lacustris* based on regulated secretion of monoamines. We find that tryptamine, phenethylamine, and tyramine, all derived via decarboxylation of amino acid precursors, elicit distinct, stereotyped canal behaviors at micromolar concentrations, and that these responses are conserved across demosponges. Using untargeted metabolomics, we demonstrate that *S. lacustris* synthesizes these molecules and identify two novel monoamine decarboxylases and vesicular transporters coexpressed in sponge neuroid cells and metabolocytes, consistent with endogenous synthesis and storage despite the absence of canonical monoamine pathway components. Phosphoproteomics and cellular imaging reveal that tryptamine couples G protein-coupled receptor (GPCR) signaling to activation of conserved Rho family GTPases, remodeling actomyosin contractility and adhesion complexes in the tent and incurrent canals, and activating NO signaling to propagate deflations. Together, these results support an ancient origin of monoaminergic signaling in early animals that was later elaborated within nervous systems to activate and modulate synaptic neuronal networks.

## Tryptamine, phenethylamine, and tyramine elicit distinct behavioral responses in sponges

We first screened 18 neurotransmitters and neuromodulators, including amino acids, acetylcholine, and mono- and polyamines across a broad concentration range (37.5 µM - 5 mM) in juveniles of the freshwater demosponge *S. lacustris*. Using overhead differential interference contrast (DIC) imaging, we found that canonical mammalian transmitters failed to trigger reproducible changes in tissue volume, canal morphology, or epithelial tent position at sub-millimolar concentrations, and induced only occasional canal twitches or minor movements at very high concentrations (≥5 mM; fig. S1). In contrast, treatment with three monoamines, tryptamine, tyramine, and phenethylamine, elicited distinct and stereotyped canal behaviors at micromolar concentrations (fig. S2).

To characterize these responses in more detail, we performed label-free optical coherence microscopy (OCM) time-lapse imaging that captures the refractive heterogeneities between water and sponge allowing for 3D visualization of the entire canal system during monoamine treatment (*9*). Treatment with tryptamine (37.5 µM) induced an immediate constriction of sponge tissue, closure of incurrent canals, and partial lowering of the epithelial tent (Fig. 1, D and E, and movie S1), with higher concentrations producing more rapid and pronounced responses. This initial phase was followed by a delayed expansion of the excurrent canal system and complete tent collapse beginning approximately 10 minutes after treatment (Fig. 1, D and F, and movie S1). This biphasic cycle, comprised of an initial constriction leading to whole-body deflation resembled endogenous behavior, in which localized twitches occasionally initiate deflation of the entire canal system (movie S2). In contrast to tryptamine, treatment with phenethylamine (37.5 µM) induced opposing morphological changes, including expansion of sponge tissues and elevation of the epithelial tent (Fig. 1, G to I, and movie S3). This resulted in increased overall tissue volume and sustained tent height (Fig. 1, H and I, and movie S3), which was distinct from the constricted and deflation-associated responses induced by tryptamine. For both monoamines, washout and replacement with normal media reverted sponge behavioral responses (fig. S3).

Unlike tryptamine and phenethylamine, treatment with tyramine (50 µM) did not induce large-scale changes in tissue or canal morphology. Instead, tyramine treatment increased the frequency of endogenous canal twitches and tonality changes over short timescales (Fig. 1, K and M), as well as the frequency of endogenous whole-body deflations over longer timescales (movie S4). In addition, we observed a gradual centripetal retraction of the sponge body over approximately 12 h, resulting in an overall reduction in tissue area (Fig. 1, J and L, and movie S4). These observations suggest that tyramine acts primarily as a modulatory signal, altering activity levels and sensitivity of endogenous contractile behaviors rather than inducing immediate large-scale morphological remodeling.

To assess whether these responses are conserved across demosponges, we performed comparable experiments in juveniles of two additional freshwater species, *Ephydatia muelleri* and *Eunapius fragilis*, as well as in reaggregated juveniles of the distantly related marine demosponge *Halichondria pinaza* (Fig. 1B). In contrast to dimethyl sulfoxide (DMSO) controls (fig. S4), all three species exhibited distinct and stereotyped responses to each monoamine (fig. S5 to S7, and movies S5 to S7). Tryptamine treatment elicited broadly similar responses across species, including constriction of inner tissues and partial lowering of the epithelial tent (fig. S5 and movie S5). Responses to phenethylamine and tyramine were likewise broadly conserved, including tent elevation in response to phenethylamine and altered activity levels and deflation frequency in response to tyramine. However, we also observed species-specific differences. In both *E. mulleri* and *E. fragilis*, phenethylamine treatment additionally resulted in noticeable expansion of excurrent canals (fig. S6, A to E, and movie S6). Tyramine responses also diverged, with *E. mulleri* displaying reduced canal activity despite similar long-term area changes, and *E. fragilis* exhibiting an immediate deflation response but only minor changes in canal activity (fig. S7 and movie S7). Together, these results indicate that demosponges exhibit a conserved capacity to detect and respond to decarboxylated monoamines, with both shared and lineage-specific behavioral responses.

## *De novo* synthesis of tryptamine, phenethylamine, and tyramine in *S. lacustris*

To directly test whether sponge cells can synthesize monoamines *de novo* (Fig. 2A), we fed heavy isotope-labeled amino acids to *S. lacustris* juveniles hatched from surface-sterilized gemmules with a depleted microbiome and performed high resolution liquid chromatography-tandem mass spectrometry (LC-MS/MS) following a 24-hour incubation sufficient for metabolic incorporation (Fig. 2B and figs. S8 and S9A). Isotopically enriched metabolites were identified at level 1 confidence (*18*) using the EMBL-MCF 2.0 reference library (*19*) with unlabeled controls used to confirm incorporation (data S1). Across all feeding conditions, we detected extensive incorporation of heavy carbon into endogenous metabolites, indicating efficient uptake and catabolic processing of amino acid precursors by gemmule-derived juveniles (fig S9 A-C). Enrichment analysis of labeled metabolites revealed over-representation of pathways associated with amino acid, purine, and nicotinamide metabolism (Fig. 2C). Comparison of isotopically labeled metabolites against standards for different monoamines only identified high levels of tryptamine, phenethylamine, and tyramine (Fig. 2D), which are each derived via a single decarboxylation reaction from their amino acid precursors (Fig. 2A). In contrast, we did not detect monoamines that require additional enzymatic modifications including hydroxylation or methylation for biosynthesis (Fig. 2A) despite sufficient sensitivity to detect other labeled metabolites (fig. S9, A to C, and data S1). For example, feeding with ^13^C_11_-tryptophan produced labeled kynurenine and kynurenic acid (fig. S9C). Similarly, ^13^C_9_-phenylalanine contributed to labeled tyrosine (fig. S9A), indicating active interconversion between aromatic amino acids.

**Fig. 2.**
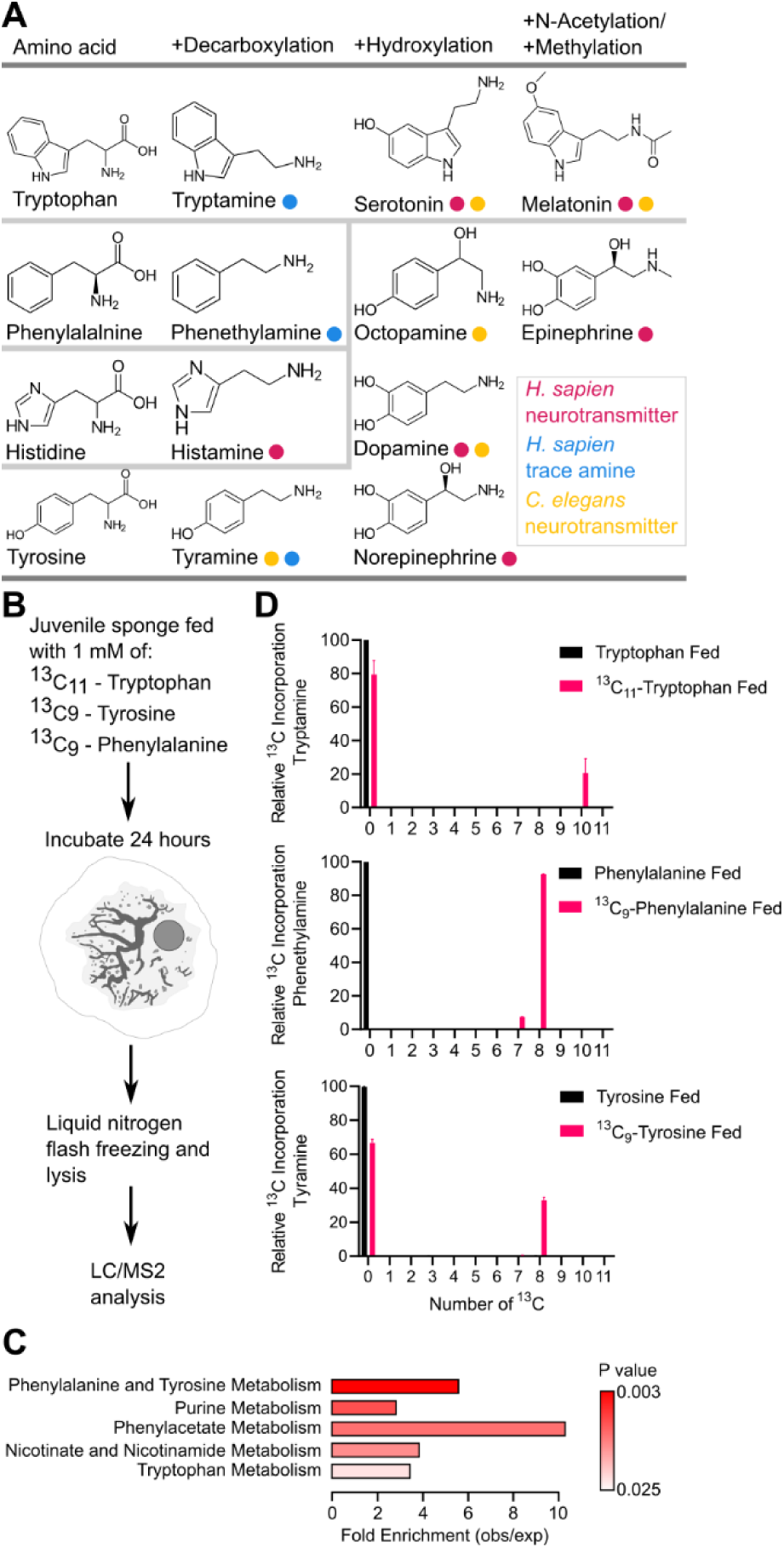
*S. lacustris* synthesizes monoamines from amino acid precursors. **(A)** Schematic classification of monoamines based on biochemical reactions required for their synthesis from amino acid precursors. **(B)** Experimental schematic of ^13^C-isotope feeding and untargeted metabolomic tracing in microbiome-depleted *S. lacustris* juveniles. **(C)** Over-representation analysis of ^13^C-labeled metabolites, showing enrichment of pathways associated with amino acid, purine, and nicotinamide metabolism. **(D)** Bar plots showing relative incorporation of ^13^C into different monoamines from control (black) or ^13^C-amino acid fed sponges (magenta). Only the monoamines tryptamine, phenethylamine, and tyramine were detected.

In sum, these results demonstrate that *S. lacustris* can synthesize the monoamines tryptamine, phenethylamine, and tyramine *de novo* from amino acid precursors. In contrast to bilaterians, which diversify monoamine signaling through additional hydroxylation and methylation reactions, *S. lacustris* appears to employ a restricted monoamine repertoire limited to decarboxylated products, providing biochemical evidence that these molecules may be synthesized for regulating behavior.

## Functional screening of putative *S. lacustris* monoamine decarboxylases and transporters

Although sponges lack canonical aromatic L-amino acid decarboxylase (AADC) and vesicular monoamine transporter (VMAT), these proteins belong to larger families of pyridoxal-5’-phosphate-dependent enzyme (PLPαF) decarboxylases and major facilitator superfamily (MFS) transporters, respectively, which include many functionally-uncharacterized members (*20*, *21*). Based on this, we searched for additional sponge genes within these families and identified two candidate PLPαF decarboxylases and nine MFS vesicular transporters expressed in the *S. lacustris* single-cell dataset (fig. S10).

To test their function, we expressed candidates in monoaminergic neurons in *C. elegans* and assayed rescue of monoamine-dependent behaviors in *Aadc* (*bas-1*), tyrosine decarboxylase (*tdc-1*) and *Vmat* (*cat-1*) mutants (*22–26*). In *C. elegans*, disruption of tyramine signaling leads to reduced backward body bends in response to anterior touch (Fig. 3A, fig. S11A) (*24*). As positive controls, *tdc-1* and *cat-1* expression in their respective mutants led to a near complete recovery of reversal behavior (Fig. 3A). Remarkably, heterologous expression of *S. lacustris* PLPαF decarboxylases *Pdxdc1* and *Pdxdc2* as well as MFS transporters *Slc17a1-8* and *Svop* also partially rescued tyramine-dependent backward body bends in *tdc-1* and *cat-1* mutants (Fig. 3A). Notably, *S. lacustris Slc18B1 a/b/c*, the closest relatives of VMAT, failed to rescue tyramine-dependent behaviors (fig. S11). We next tested dopamine- and serotonin-dependent behaviors by rescuing basal slowing rate when encountering a bacterial food source and using serotonin immunofluorescence (*22*, *23*). As with tyramine, reintroduction of C. elegans *bas-1* and *cat-1* rescued the dopamine-dependent slowing response and serotonin staining (Fig. 3B-C, fig. S11B). *S. lacustris Pdxdc1*, *Slc17A1-8*, and *Svop* also partially restored basal slowing response and serotonin levels (Fig. 3B-C, fig. S11B), suggesting a broad substrate recognition for production or transport of different aromatic monoamines. In contrast, *S. lacustris Pdxdc2* failed to rescue dopamine-dependent slowing and displayed modest serotonin staining, suggesting it is specialized for tyramine synthesis similar to *C. elegans tdc-1* (Fig. 3B-C, fig. S11B). As further validation, we found that AlphaFold3 (AF3) predicted structures of *S. lacustris* decarboxylases and vesicular transporters had structural folds comparable to human homologs supporting their monoaminergic functions (Fig. 3D and E). Together, these results define a noncanonical monoaminergic toolkit in *S. lacustris*, with *Pdxdc1* as a candidate broad aromatic decarboxylase, *Pdxdc2* as a tyramine-specific decarboxylase, and *Slc17a1-8* and *Svop* as promiscuous vesicular transporters.

**Fig. 3.**
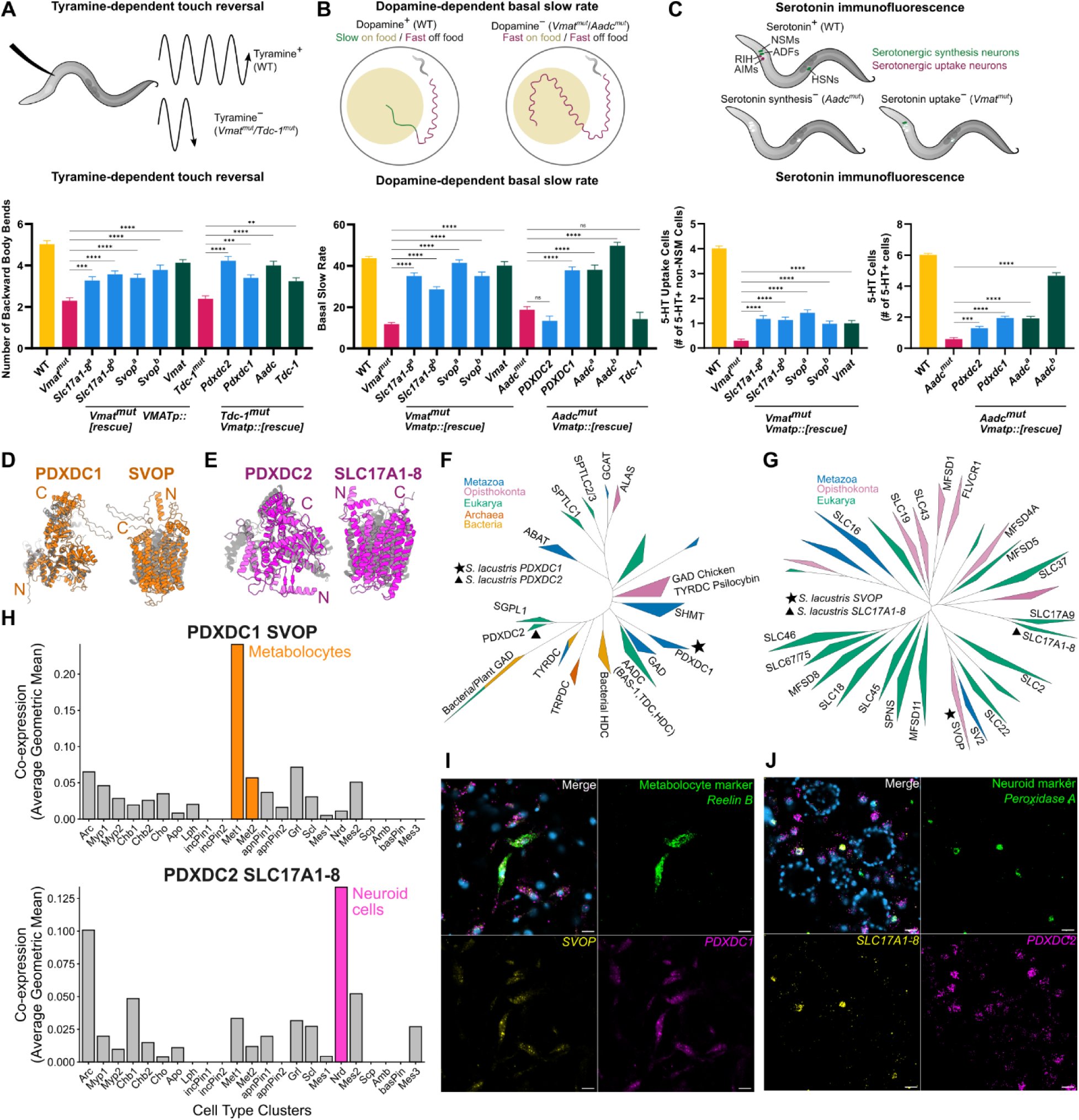
Monoamine production machinery in *S. lacustris* metabolocytes and neuroid cells. **(A)** Top: Experimental schematic used to evaluate tyramine synthesis and storage as part of the *C. elegans* tyramine-dependent touch reversal response. Bottom: Bar graph quantification of backward body bends in response to anterior touch of wild-type (WT) animals (N2, n = 22), *Vmat* mutants (*cat-1(ok411),* n = 25), and *Tdc-1* mutants (*tdc-1(ok914),* n = 25). Expression of *S. lacustris Slc17a1-8* (strain a n = 27; strain b n = 27), *Svop* (strain a, n = 25; strain b, n = 23), *Pdxdc2* (n = 26), *Pdxdc1* (n = 25) and *C.* elegans *Vmat* (*cat-1,* n = 28), *Aadc* (*bas-1*, n = 24), *Tdc-1* (*tdc-1*, n = 25) significantly increased the number of backward body bends compared to *Vmat* or *Tdc-1* mutants. Nonmatching one-way ANOVA followed by Šídák test with a 95% confidence interval (α = 0.05); ns = not significant P ≥ 0.05; **P < 0.01; *** P < 0.001; **** P < 0.0001; bars show mean with SEM. **(B)** Top: Experimental schematic used to evaluate dopamine synthesis and storage as part of the *C. elegans* dopamine-dependent basal slowing response. Bottom: Bar graph quantification of basal slow rate, where higher bar indicates slower movement on food, of well-fed WT animals (yellow, N2, n = 23), *Vmat* mutants (magenta, *cat-1(ok411)*, n = 22), and *Aadc* mutants (magenta, *bas-1(tm351)*, n = 22) on bacteria. Expression of *Spongilla* (blue) *Slc17a1-8* (strain a n = 25; strain b n = 27), *Svop* (strain a n = 22; strain b n = 30), *Pdxdc1* (n = 24) and *C.* elegans (green) *Vmat* (*cat-1,* n = 23), *Aadc* (*bas-1*, strain a, n = 16, strain b, n = 23), significantly increased the basal slow rate compared to *Vmat* and *Aadc* mutants. Expression of *S. lacustris* (blue) *Pdxdc2* (n = 23) or *C. elegans* (green) *Tdc-1* (*tdc-1,* n = 18) did not increase the basal slow rate compared to *Vmat* and *Aadc* mutants. Nonmatching one-way ANOVA followed by Šídák test with a 95% confidence interval (α = 0.05); ns = not significant P ≥ 0.05; **** P < 0.0001; bars show mean with SEM. **(C)** Top: Experimental schematic used to evaluate serotonin synthesis and storage by staining serotonergic neurons and serotonin uptake neurons in the head of *C. elegans*. Bottom: Quantification of *Vmat*-dependent serotonin positive head neurons in adult/late L4 WT animals (yellow, N2, n = 90) and *Vmat* mutants (*cat-1(*ok411), n = 90) or total serotonin positive head neurons in adult/late L4 WT animals and *Aadc* mutants (*bas-1(tm351)*, n = 90) after immunofluorescence with an anti-serotonin antibody. Expression of *S. lacustris* (blue) *Slc17a1-8* (strain a, n = 90; strain b, n = 90), *Svop* (strain a, n = 90; strain b, n = 90), Pdxdc2 (n = 90), Pdxdc1 (n = 90) and *C.* elegans (green) *Vmat* (*cat-1,* n = 88), *Aadc* (*bas-1,* strain a, n = 90, strain b, n = 90), significantly increased the number of serotonin positive head neurons compared to *Vmat* and *Aadc* mutants. Nonmatching one-way ANOVA followed by Šídák test with a 95% confidence interval (α = 0.05); ns = not significant P ≥ 0.05; *** P < 0.001; **** P < 0.0001; bars show mean with standard errors of the mean. **(D)** AF3 structures of *S. lacustris* Pdxdc1 (orange, residues 1-644) and Svop (orange, residues 21-568) aligned to AF3 structure of human PDXDC1 (grey, residues 11-600) and experimental structure of human VMAT1 (grey, 8TGJ). **(E)** AF3 structures of *S. lacustris* Pdxdc2 (magenta, residues 61-572) and Slc17A1-8 (magenta, residues 26-498) aligned to experimental structure of human AADC (grey, 3RBL) and experimental structure of human VMAT1 (grey, 8TGJ). **(F)** Maximum-likelihood phylogenetic tree of PLPαF decarboxylases. **(G)** Maximum-likelihood phylogenetic tree of MFS vesicular transporters. **(H)** Bar plot showing co-expression of *S. lacustris Pdxdc1* with *Svop* and *Pdxdc2* with *Slc17a1-8* across sponge cell types. **(I)** HCR staining of *S. lacustris Pdxdc1*, *Svop,* and metabolocyte cell marker *Reelin B*. Scale bar, 10 μm. **(J)** HCR staining of *S. lacustris Pdxdc2*, *Slc17a1-8*, and neuroid cell marker *Peroxidase a*. Scale bar, 10 μm.

## *S. lacustris* monoamine decarboxylases and transporters belong to alternative PLPαF and MFS subfamilies

To reconstruct the evolutionary history of *S. lacustris* decarboxylases and vesicular monoamine transporters, we inferred phylogenies for PLPαF and MFS superfamilies from a curated set of eukaryote proteomes (Fig. 3F and G, data S2). *Pdxdc1* is a one-to-one ortholog of human *PDXDC1*, which represents a largely uncharacterized clade that diverged from *Aadc* and *Gad* prior to the last common ancestor of animals and fungi (Opisthokonta). *Pdxdc2* belongs to a separate orthology group that also predates opisthokonts and was subsequently lost in multiple animal lineages, including vertebrates, arthropods, and nematodes (Fig. 3F, fig. S12). Within MFS transporters, *S. lacustris Svop* groups with mammalian synaptic vesicle glycoprotein 2-related proteins (*SVOP* and *SVOPL)*, that represent conserved synaptic vesicle proteins with no identified transport substrates (*27*) (Fig. 3G, fig. S13). However, our tree shows this orthology group as a sister clade to the *Slc22*a subfamily, which transports a diverse array of organic cations including monoamines, raising the possibility that monoamines were ancestral substrates for the combined clade. Lastly, *S. lacustris Slc17A1*-8 is in a clade sister to the vesicular glutamate, nucleotide, and sialin transporters, indicating it diverged prior to the diversification of these specialized organic anion transporters in bilaterians (Fig. 3G, fig. S13). Together, these phylogenies place the sponge decarboxylases and vesicular transporters in ancient, sparsely characterized PLPαF and MFS subfamilies, consistent with a noncanonical monoaminergic toolkit that predates *Aadc* and *Vmat*.

## Monoamine pathway genes are coexpressed in metabolocytes and neuroid cells

We next asked whether sponge decarboxylases and transporters are coexpressed in specific sponge cell types, a requirement for regulated monoaminergic signaling. In the *S. lacustris* single-cell atlas (*10*), *Pdxdc1* and *Svop* were coexpressed in metabolocytes, a metabolic storage cell type localized near the tent and incurrent canals, whereas tyramine-specific *Pdxdc2* and *Slc17a1-8* were enriched in neuroid cells that closely associate with sponge digestive chambers (Fig. 3H, fig S14). We validated these patterns using Hybridization Chain Reaction (HCR), which confirmed colocalization of *Pdxdc1* and *Svop* with a metabolocyte marker and of *Pdxdc2* and *Slc17a1-8* with a neuroid cell marker (Fig. 3 I and J). *Slc17a1-8* was also detected in some metabolocytes, and *Pdxdc1*, *Svop*, and *Pdxdc2* each individually displayed scattered expression in other cells, including archaeocytes and adjacent progenitors. Together, these data suggest metabolocytes and neuroid cells are candidate monoaminergic secretory cell types and are positioned to enact spatially segregated signaling along the sponge canal system.

## Tryptamine couples Rho/Rac-linked contractility, adhesion remodeling, and NO signaling

Our behavioral observations showed that tryptamine induces rapid closure of incurrent canals followed by whole-body deflations. To determine the molecular basis of this response, we performed quantitative phosphoproteomics at 3, 15, and 30 minutes after treatment with 37.5 µM tryptamine or dimethyl sulfoxide (DMSO) (Fig. 4A). We identified 2385 differentially phosphorylated peptides mapping to 1266 unique proteins (Fig. 4B, fig. S15, data S3). Phosphorylation changes were strongly time-dependent, with most regulated phosphosites detected at 15 and 30 minutes (1243 and 1086, respectively), and the largest shift occurring between 3 and 15 minutes (1867; Fig. 4B, coinciding with the major changes in canal morphology. Enrichment analysis of regulated proteins found overrepresentation of GTPase activator activity, cadherin binding, and cytoskeletal functions (Fig. 4C).

**Fig. 4.**
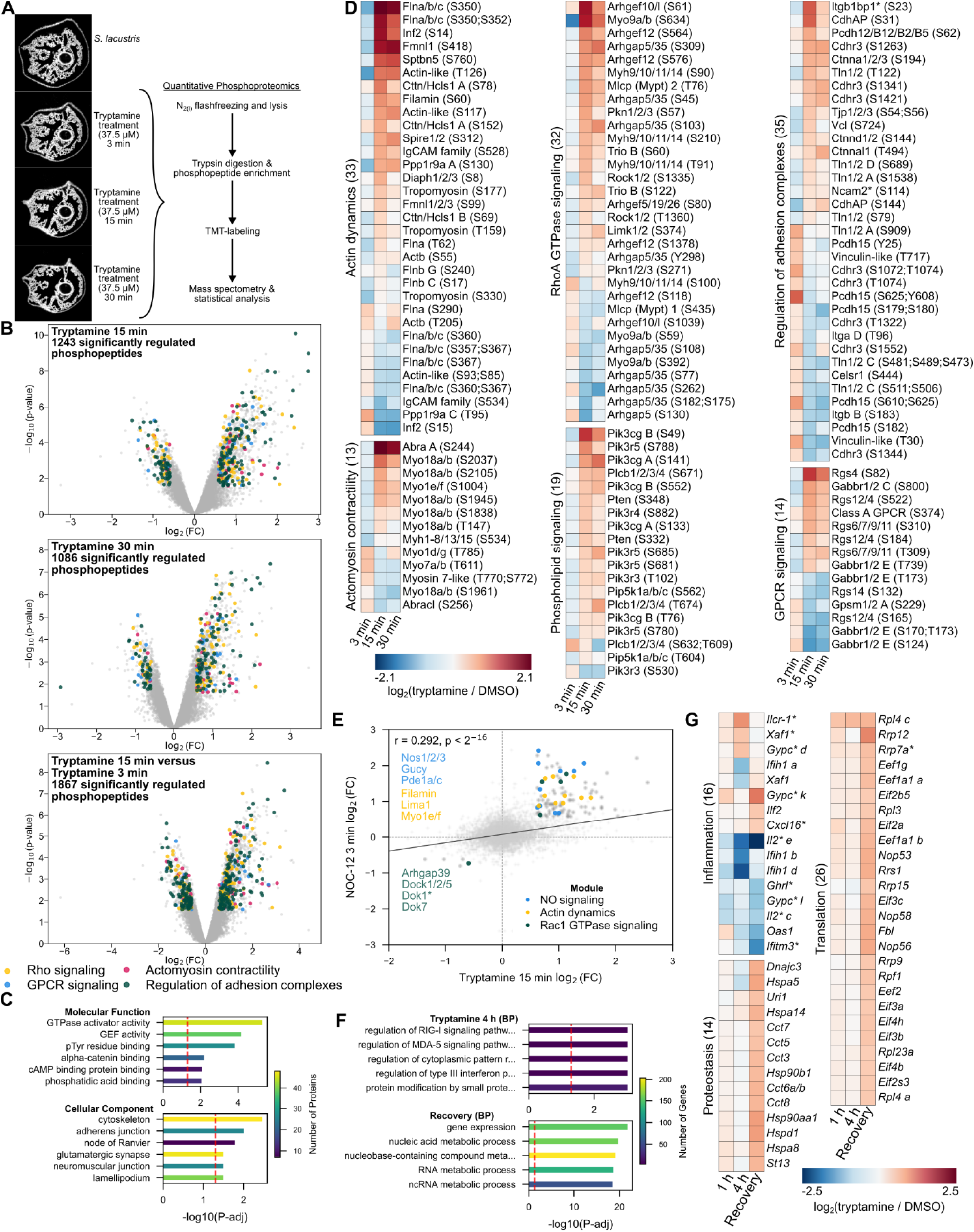
Phosphoproteomics of tryptamine-treated *S. lacustris*. **(A)** Experimental setup of sponge phosphoproteomics. *S. lacustris* juveniles were treated with 37.5 µM tryptamine for 3, 15, and 30 minutes before sample processing for phosphoproteomics. **(B)** Volcano plots illustrating quantitative differences in phosphopeptides between tryptamine- or DMSO-treated *S. lacustris*. The x-axis shows log2 fold changes (FC) of tryptamine/DMSO at 15 min (upper panel), tryptamine/DMSO at 30 min (middle panel), and tryptamine 15 min/3 min (after normalization to DMSO, lower panel). The y-axis represents -log₁₀ p-values. Significantly regulated phosphopeptides that belong to proteins related to molecular modules of interest are colored. GPCR, G-protein coupled receptor. Rho, Rat sarcoma homolog family. **(C)** Top enriched molecular function and cellular compartment GO terms of significantly differentially phosphorylated proteins after tryptamine treatment. **(D)** Heatmaps of statistically significant phosphosites in cellular modules identified in tryptamine-treated S. lacustris. Gene names indicate orthology to *H. sapiens* genes. For genes marked with an asterisk, homology to *H. sapiens* was inferred using the protein language model-based PRotein Ortholog Search Tool (PROST). **(E)** Phosphosite abundance calculated relative to timepoint-matched DMSO-treated controls. Correlation of phosphopeptide fold changes between tryptamine-treated and NO-treated *S. lacustris*. Colored proteins represent significant phosphorylation changes in both treatments related to molecular modules of interest. **(F)** Top enriched biological process (BP) categories for significantly differentially expressed transcripts from tryptamine (37.5 µM)-treated *S. lacustris*. **(G)** Heatmaps of differentially expressed genes of three molecular modules of interest based on manual annotation in tryptamine (37.5 µM)-treated *S. lacustris*. The colormap indicates fold change in transcript counts between time-matched tryptamine- and DMSO-treated samples.

Across the time series, tryptamine induced phosphosite changes in contractile machinery associated with vertebrate stress fibers and smooth muscle (Fig. 4D, fig. S16). This included phosphosite changes in actin (Actb) and non-muscle myosin heavy chains (Myh9/10/11/14), as well as Rho-associated protein kinase (Rock1/2) and myosin light chain phosphatase (Ppp1r12/16), key regulators of actomyosin tension (Fig. 4D, fig. S16) (*28*, *29*). We also observed phosphorylation changes in actin modifiers, including filamins, formins, and cofilins, as well as in unconventional myosins Myo1e/f and Myo18a/b that link actin to the cell membrane, suggesting remodeling of force-bearing actin networks during the response (*30–32*). This contractile signature was accompanied by differential phosphorylation of junctional and mechanotransduction proteins, including α- and δ-catenin, talin (Tln1/2), integrin β, vinculin, and numerous non-classical cadherins (*33*, *34*). Notably, tryptamine also regulated phosphorylation of conserved exocytosis proteins including Unc13, syntaxins (Stx/Stxbp), synaptosomal-associated protein 27 (SNAP27), synaptotagmin 7 (Syt7), and post-synaptic scaffolding Dlg and Shank proteins (fig. S16) (*35*). The activation of these mechanotransduction pathways by tryptamine is further reflected with specific phosphorylation of actin-binding rho-activating (ABRA) protein that promotes nuclear localization of myocardin-related transcription factor (MRTF) (Fig. 4D, fig. S16), previously demonstrated to differentially regulate expression of contractility genes in *E. mulleri* (*15*). Together, these data point to coordinated regulation of actomyosin and adhesion machinery as molecular drivers of morphological changes observed in the canal system during tryptamine treatment.

In addition to these cellular contractile and junction proteins, the pattern of regulated phosphosites strongly implicated two antagonistic pathways coupled to GPCR signaling (Fig. 4D). First, we detected differential phosphorylation of multiple GPCRs and regulators of G-protein signaling (*Rgs*) (fig. S16) (*36*), consistent with activation of heterotrimeric G-protein signaling during the tryptamine response. Second, the coordinated phosphorylation of rho guanine nucleotide exchange factors (Trio, Arhgef10/12) and Rock/filamin contractile signature described above is characteristic of a Gα12-linked pathway in which a G protein activates RhoA-centered contractility and adhesion junction maturation via RGS-domain Arhgef12 (*37*, *38*). In parallel, tryptamine induced phosphorylation changes in phosphoinositide 3-kinase-γ (PI3Kγ) and phospholipase C β (PLCβ), canonical effectors frequently recruited downstream of Gβγ to drive phosphoinositide/diacylglycerol second messenger signaling and recruitment of cytoskeletal adapters to promote junctional remodeling and actin-based membrane protrusions (*39*). Together, these signatures support a model in which tryptamine simultaneously engages both a contractility-based Gα12-RhoGEF-Rock pathway and a membrane/second-messenger Gβγ-PI3Kγ/PLCβ arm (fig. S16).

A delayed expansion of excurrent canals resembles the deflation response induced by NO. To explore this, we compared tryptamine-regulated phosphosites to a previously published phosphoproteomic dataset of NO-treated sponges. Consistent with activation of NO/cyclic guanosine monophosphate (cGMP) signaling, tryptamine treatment induced phosphosite changes in NO synthase (NOS), guanylate cyclase (GUCY), and phosphodiesterase (PDE1a/c) (Fig. 4E) (*9*). Overall, the tryptamine and NO responses shared 82 phosphosites at 15 minutes and 66 phosphosites at 30 minutes, with all but one shared site changing in the same direction relative to DMSO control. Among shared targets, phosphorylation of rho GTPase activating protein 39 (Arhgap39) and LIM domain and actin-binding protein 1 (Lima1) is consistent with activation of Rac/Cdc42 remodeling known to promote membrane ruffling and protrusive actin states in other systems (Fig. 4E) (*40*, *41*). Shared regulation of adhesion and membrane-coupling components, including Myo1/18, further supports coordinated remodeling of cell junctional and cytoskeletal machinery (Fig. 4E).

## Tryptamine activates inflammation and growth transcriptional programs during recovery

In bilaterians, monoamines such as serotonin can activate transcriptional programs in contractile tissues that extend beyond the initial response (*42*). To determine whether sponges also exhibit a longer-term transcriptional response to tryptamine, we performed bulk RNA sequencing 1 and 4 hours post-treatment, and after 24 h recovery following washout (data S4). Transcriptional responses were minimal after 1 hour, but included downregulation of *Nfat* and *Gucy*, consistent with attenuation of NO-cGMP signaling (Fig. 4F–G; fig. S17) (*43*, *44*). In contrast, prolonged exposure (4 h) induced broad transcriptional changes (559 genes) enriched for inflammatory and stress pathways, including upregulation of a cytokine-like gene (*Ilcr1*), downregulation of immune sensing machinery (*Mda5*/*Ifih1*), and decreased indoleamine 2,3-dioxygenase 1 (*Ido*1/2) expression, suggesting altered tryptophan metabolism (Fig. 4, F and G, and fig. S17) (*45*, *46*). During recovery (24 h post-washout), we observed extensive transcriptional changes (992 genes) enriched for translation and protein homeostasis programs, together with increased expression of extracellular matrix-associated genes, including collagens (Fig. 4, F and G, and fig. S17). These results suggest that tryptamine signaling remodels sponge tissue over multiple timescales, spanning immediate attenuation of NO-cGMP pathways, inflammatory and stress programs after sustained exposure, and growth and extracellular matrix-associated programs during recovery.

## Tryptamine activates GPCR signaling in incurrent contractile cell types

Tryptamine treatment induced differential phosphorylation of three GPCRs, consistent with activation-dependent GPCR regulation during the behavioral response. To systematically evaluate candidate receptors for all three monoamines, we performed AF3 docking (*47*) of tryptamine, phenethylamine, and tyramine against all *S. lacustris* GPCRs with detectable single-cell expression (*10*) and constructed a GPCR phylogeny to infer orthology and assign candidates to receptor classes (*48*) (Fig. 5A, figs. S18–S20, and data S5). AF3 predicted compatible poses for tryptamine in two of the three phosphorylated GPCRs, belonging to class A (iPTM = 0.93) and class C receptor families (iPTM = 0.59) (Fig. 5B). AF3 additionally identified a candidate phenethylamine-responsive class A GPCR and a candidate tyramine-responsive class E GPCR (Fig. 5A; fig. S18–S21). All candidate monoamine GPCRs are differentially expressed in contractile cell types and choanocytes, consistent with monoamine behavioral responses (Fig. 5A).

**Fig. 5.**
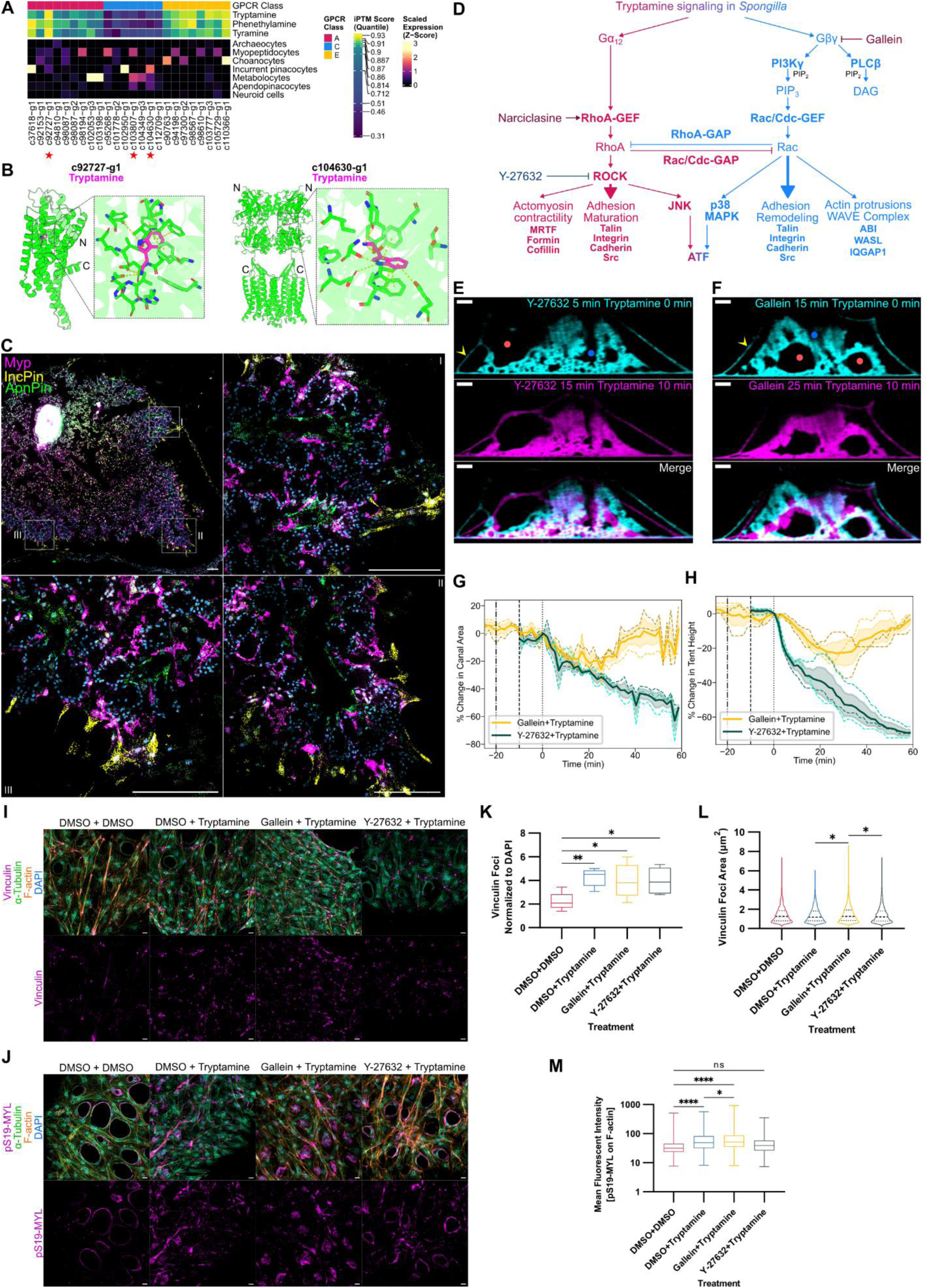
Tryptamine signal transduction in *S. lacustris* myoepithelial cells. **(A)** Scaled GPCR expression across major *S. lacustris* cell types. GPCRs are grouped by likely class (fig. S20) and annotated with AF3 docking iPTM scores for tryptamine, phenethylamine, and tyramine. Red stars mark GPCRs phosphorylated after tryptamine treatment. **(B)** AF3 docking models of tryptamine bound to class A GPCR (c92727-g1, residues 1-365) and class C GPCR (c104630-g1, residues 20-770), with predicted binding pockets highlighted. **(C)** HCR labeling of myopeptidocytes (Myo), incurrent pinacocytes (IncPin), and apendopinacocytes (ApnPin). Scale bar, 100 μm. **(D)** Model for tryptamine signal transduction. **(E,F)** Representative OCM cross-sections after pretreatment with Y-27632 (20 μM) (E) or gallein (5 μM) (G) followed by tryptamine (37.5 μM). Cyan and magenta indicate t = 0 and t = 10 min. Markers: yellow arrowhead, outer epithelial tent; blue dots, incurrent canals; red dots, excurrent canals. Scale bars, 100 μm. **(G,H)** Canal area (G) and tent height (H) normalized to pretreatment during inhibitor pretreatment and subsequent tryptamine exposure (n = 3 biological replicates). Lines indicate replicates, shading indicates SEM, vertical lines indicate additions of gallein, Y-27632, and tryptamine. **(I,J)** Immunofluorescence for vinculin (I) and pMYL(S19) (J) after DMSO with tryptamine, or Y-27632 or gallein with tryptamine. Scale bars, 10 μm. **(K–M)** Quantification of vinculin foci per nucleus (K), vinculin foci area (L), and pMYL(S19) intensity on F-actin fibers (M) (n = 6). Non-matching one-way ANOVA with Šídák (K,M) or Kruskal–Wallis with Dunn’s (L); ns P ≥ 0.05, *P < 0.05, **P < 0.01, ****P < 0.0001.

Tryptamine-responsive GPCRs are expressed in three contractile cell types, including incurrent pinacocytes, apendopinacocytes, and myopeptidocytes, a recently described population of mesohyl cells expressing smooth-muscle contractile genes (*10*). To resolve the spatial distribution of these putative effector populations, we performed whole-body hybridization chain reaction (HCR) targeting cell type-specific markers (Fig. 5C). Incurrent pinacocytes are largely restricted to the outer tent and subdermal lacunae that form the initial portion of the water canal system (Fig. 5C). In contrast, myopeptidocytes form a continuous cellular network lining much of the incurrent canal system, reminiscent of sieve cells and reticuloapopylocytes described in other sponges (*49*, *50*). This network extends from the canals into the mesohyl to enwrap individual choanocyte chambers, consistent with a role regulating water flow into individual chambers (Fig. 5C). Finally, apendopinacocytes formed the principal lining of excurrent canals and osculum through which water exits the sponge (*10*) (Fig. 5C). To further examine how contraction of these canal-lining cells affects tryptamine-induced behavior, we imaged the outer tent and adjacent choanocyte chambers by DIC during tryptamine treatment. Tryptamine led to immediate closure of tent ostia followed by closure of incurrent canals and constriction of choanocyte chambers (Movie S8). Together, spatial segregation of contractile cell types along the canal system positions distinct effector populations to independently regulate canal tonality, consistent with tissue responses to tryptamine treatment.

## RhoA–Rock and Gβγ signaling are required for tryptamine-induced canal remodeling

To functionally test the two intracellular signaling pathways implicated by phosphoproteomics (Fig. 5D), we treated sponges with tryptamine in combination with either the Rock inhibitor Y-27632 (*51*) or the Gβγ inhibitor gallein (*52*) (Fig. 5E and F). Neither inhibitor alone produced substantial behavioral changes, with Y-27632 producing no obvious effect and gallein inducing only minor canal shape changes within the first ∼15 minutes (fig. S22). Co-treatment with Y-27632 and tryptamine abolished the typical tryptamine response and instead produced progressive collapse of the tent and canal system (Fig. 5E, G to H, and movie S9), as well as expansion of choanocyte chambers and membrane ruffling (fig. S23 A-B, E-F). Later in treatment, the tent separated into distinct layers (movie S9), consistent with loss of actomyosin-dependent mechanical integrity. In contrast, co-treatment with gallein and tryptamine resulted in constriction of canals comparable to tryptamine alone but without tent lowering, except at later time points during NO-mediated endogenous deflations (Fig. 5, F to H, and movie S10).

Excurrent canals also appeared more constricted at later time points relative to tryptamine alone (movie S1 and S10). Higher resolution imaging showed that gallein treatment resulted in incomplete ostia closure, supporting a specific role for Gβγ signaling in tent epithelial shape change (fig. S23 A-B, C-D). Together, these perturbations indicate that Rock-dependent contractility and Gβγ-dependent epithelial changes are both required for tryptamine-induced canal and tent behaviors.

To connect the Rock and Gβγ perturbation phenotypes to changes in adhesion and contractility at the cellular level, we stained tent incurrent pinacocytes for vinculin (*33*) and phosphorylated myosin light chain (pMYL) (*15*) in control, tryptamine-treated, and tryptamine and inhibitor co-treated sponges (Fig. 5 I and J). In control sponges, incurrent pinacocytes formed a myoepithelial sheet characterized by long intercellular F-actin cables and discrete vinculin foci (Fig. 5I), consistent with previous studies (*9*, *15*). Following tryptamine treatment, the number of vinculin foci increased (Fig. 5 I and K), indicating recruitment of additional adhesion sites during the response. When Gβγ signaling was inhibited with gallein during tryptamine treatment, vinculin foci became larger (Fig. 5 I and L), suggesting that Rock promotes adhesion maturation. We observed a similar pattern in actomyosin-based contractility. Basal pMYL levels along F-actin were similar in control and Y-27632-treated sponges (Fig. 5 J and M), indicating that Rock inhibition does not substantially alter baseline pMYL signal under these conditions. In contrast, tryptamine strongly increased pMYL intensity on F-actin, and this increase was further elevated in sponges co-treated with gallein (Fig. 5 J and M), consistent with enhanced contractile fiber phosphorylation under Gβγ-dependent pathway inhibition. In addition, we observed tent ostia marked by high cytoplasmic pMYL and circular α-tubulin staining (Fig. 5 I and J). Incurrent pinacocytes lining the subdermal lacunae showed similar patterns (fig. S24), whereas signal in the choanoderm could not be reliably quantified due to tissue thickness.

## Discussion

In animals with nervous systems, monoamines act at synapses or extrasynaptically through volume transmission to modulate organismal physiology and regulate contractile cell types. Here, we show that a signaling system using monoamines derived through a single decarboxylation step from amino acid precursors exists in *Spongilla lacustris* (Fig. 6A), an early branching animal lacking canonical neurons and muscles. We identify three endogenously synthesized monoamines - tryptamine, phenethylamine, and tyramine - that elicit distinct contractile behaviors within the canal system that are conserved in demosponges. Two secretory cell types coexpress biosynthetic enzymes and vesicular transporters for these transmitters, metabolocytes expressing *Pdxdc1* and *Svop*, most consistent with tryptamine and phenethylamine signaling, and neuroid cells expressing *Pdxdc2* and *Slc17A1-8*, consistent with tyramine signaling. These signals act on spatially segregated effector tissues arrayed along the canal system, including incurrent pinacocytes that compose the outer tent and subdermal lacunae, a network of myopeptidocytes comprising the interior incurrent canals, and apendopinacocytes lining excurrent canals and the osculum. Tryptamine drives rapid closure of the incurrent system via activation of RhoA-ROCK and Gβγ-PI3Kγ signaling and then triggers a NO-associated deflation, phenethylamine induces the opposing state, sustaining inflation and tent elevation, and tyramine modulates endogenous activity and deflation propensity over longer timescales. Together, these results define a distributed chemical transmitter circuit in which decarboxylated amines act as local paracrine regulators of canal tone, while NO, a membrane permeant gas, acts as a fast broadcast signal to coordinate whole-body deflations. Consistent with the origin of this circuit in early animals, recent receptor deorphanization in placozoans identified GPCRs responsive to tryptamine, phenethylamine, and tyramine, and showed they elicit distinct effects on locomotion speed and body shape (*53*), similar to the behaviors we observe in sponges.

**Fig. 6.**
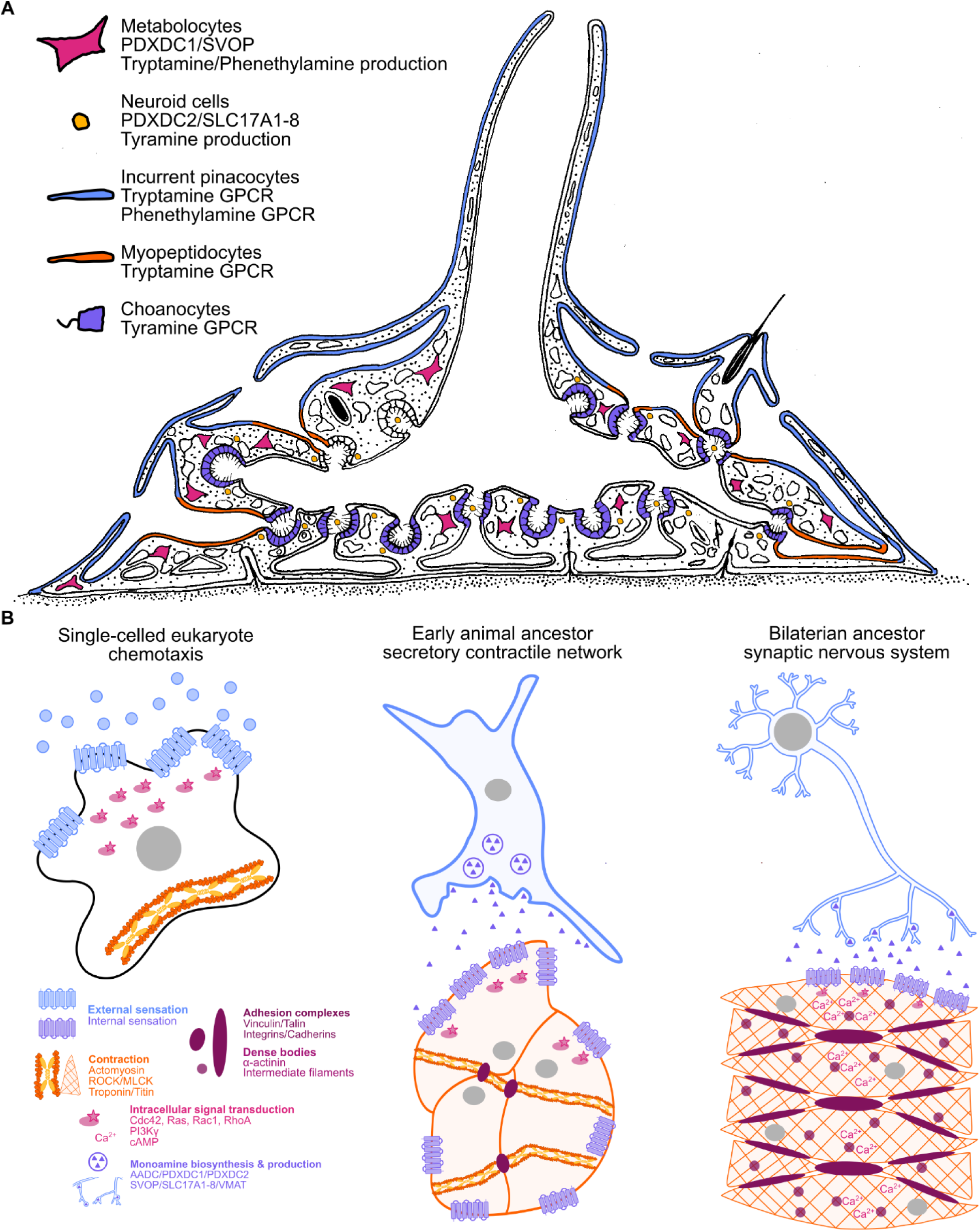
Sponge monoaminergic circuit and its evolutionary assembly. **(A)** Model of monoamine signaling in *S. lacustris*. **(B)** Hypothesized evolutionary assembly of monoaminergic signaling. Unicellular eukaryotes couple hemosensation to actomyosin driven motility via Rho family GTPases. In early animals, division of labor separated sensory secretory cells that release amines from contractile epithelia that interpret these cues through GPCR signaling to Rho GTPases, adhesion complexes, and actomyosin. In bilaterians, amines were elaborated as neurotransmitters integrated into synaptic nervous systems and musculature. Abbreviations: ROCK, rho associated coiled coil containing protein kinase; MLCK, myosin light chain kinase; FAK, focal adhesion kinase; GPCRs, G protein coupled receptors; PI3Kγ, phosphatidylinositol 3 kinase catalytic subunit gamma; cAMP, cyclic adenosine monophosphate.

Our study also shows that the sponge monoaminergic system is built from biosynthetic enzymes, vesicular transporters, and receptors drawn from the same broader gene families as the canonical toolkit, but from distinct ortholog groups. Sponges lack clear *Aadc* and *Vmat* orthologs, the enzyme and vesicular transporter traditionally considered essential for monoamine signaling (***17***). Instead, *S. lacustris* possesses *Pdxdc1* and *Pdxdc2* for decarboxylation and *Svop* and *Slc17A1-8* for vesicular transport, and these proteins functionally rescue monoamine-dependent behaviors in *C. elegans* mutants lacking the canonical pathway. On the receptor side, candidate sponge monoamine GPCRs span at least three receptor classes, class A, class C, and an expanded clade of sponge receptors most closely related to class E receptors in *Dictyostelium* slime molds. This raises the question of whether these genes are sponge-specific inventions or remnants of a larger ancestral toolkit. Indeed, monoamine receptors have evolved repeatedly within bilaterian GPCR lineages (*54–56*), indicating at least some components of this toolkit may be specific to sponges. However, *Pdxdc1*, *Svop*, and members of the *Slc17A1-8* family originated early in eukaryotes, were present in the animal ancestor, and are highly conserved across animals. *Pdxdc1* is a single-copy ortholog in all major animal lineages and is expressed in mouse hippocampus, striatum, and brainstem, where it has been linked to schizophrenia-related phenotypes and implicated in monoamine signaling (*57*). *Svop* and its paralogs *Sv2a/b/c* are key constituents of synaptic vesicles and members of the MFS family, but with no established substrates. *Slc17A1-8*, positioned phylogenetically as a sister group to the bilaterian *SLC17* family that later diversified into vesicular glutamate and organic anion transporters, appears permissive to cationic monoamines in *S. lacustris*, which may reflect an ancestral substrate breadth early in animal evolution (*20*). Lastly, *Pdxdc2* represents an ancient eukaryotic orthology group that was lost in vertebrates and ecdysozoans but retained in many other invertebrate clades, where it may serve as a tyrosine decarboxylase, similar to Tdc-1 in *C. elegans*. Together, these findings suggest that the molecular toolkit for monoamine signaling in animals is broader than currently appreciated and that components of this alternative pathway may be deployed where canonical machinery is absent.

Strikingly, most components of monoaminergic signaling present in sponges predate animals (Fig. 6B). Secretion, GPCR-based chemosensation, and small GTPase control of actomyosin contractility are widespread in unicellular eukaryotes (*58*, *59*). This suggests that a key innovation in early animals occurred through cellular division of labor, in which synthesis and regulated secretion became core machinery of sensory secretory cells monitoring internal or external cues, while effector cells, including contractile epithelia, inherited monoamine reception coupled to regulation of actomyosin and adhesion machinery to generate tissue-level contractions. The transition to synaptic nervous systems was accompanied by elaboration of the monoamine repertoire and the addition of other transmitters (*60*, *61*). Tryptamine, phenethylamine, and tyramine are effective for short-range paracrine signaling in part because they readily diffuse and are cleared rapidly by monoamine oxidases (*62*). The emergence of additional monoamine modifications, particularly hydroxylated derivatives, enabled more efficient storage in vesicles and longer-acting neuromodulatory activity. Remarkably, historical roles of monoamines are still apparent in bilaterians, where monoamines often couple with NO to regulate contractility and fluid flow in contexts that are not synaptic, including gut peristalsis, vascular tonality, and myoepithelial contractions regulated by GPCR coupling to RhoA-ROCK signaling (*63*, *64*).

## Acknowledgments

We thank A. Horton for providing *Eunapius fragilis* gemmules and S. Nichols for providing anti-*Em*Vinculin antibody. We thank Haig Keshishian, Paul Forscher, Michael Higley, and members of the Musser Lab for fruitful discussions and feedback. We also thank the Yale Center for Genome Analysis (YCGA), EMBL Metabolomics Core Facility, EMBL Proteomics Core Facility, and EMBL mechanical and electronic workshops for their help with sample processing, analysis, custom manufacturing and other support.

## Funding

Human Frontiers Science Program Grant RGP024/2023 (JMM)

Natural Sciences and Engineering Research Council of Canada/Conseil de recherches en sciences naturelles et en génie du Canada CGS D - 587666 - 2024 (RXZ)

NIGMS National Institutes of Health grant 1S10OD030363-01A1 (YCGA)

European Research Council Consolidator grant 864027/Brillouin4Life (RP, LW)

HORIZON-INFRA-2022-TECH-01 grant 101094250/IMAGINE (RP, LW)

European Molecular Biology Laboratory Germany (BD, FS, MR, JJS, AI, RP, LW)

NIH New Innovator’s Award 1DP2GM154014-01 (MOD)

## Author contributions

Conceptualization: RXZ, JMM

Methodology: RXZ, NM, JDM, KJ, BD, FS, MN, CJR, MR, JS, LW, RP, JMM

Investigation: RXZ, NM, JDM, KJ, BD, FS, MR, JS, JMM

Software: RXZ, KJ, FS, JMM

Formal analysis: RXZ, KJ, BD, FS, JMM

Visualization: RXZ, JMM

Funding acquisition: RXZ, AK, SW, JMM, RP

Project administration: RXZ, JDM, JMM

Supervision: RXZ, JMM

Writing – original draft: RXZ, JMM

Writing – review & editing: RXZ, NM, JDM, KJ, BD, FS, MN, CJR, MR, JS, LW, RP, AI, SW, MPO, JMM

## Declaration of AI usage

We utilized ChatGPT 5.2, Gemini 3, and Opus 4.5 and 4.6 to generate code for plotting, and ChatGPT 5.2 for proofreading and minor clarity suggestions of the manuscript. AI-generated code was carefully scrutinized and annotated to ensure accuracy.

---

**Fig. S1.**
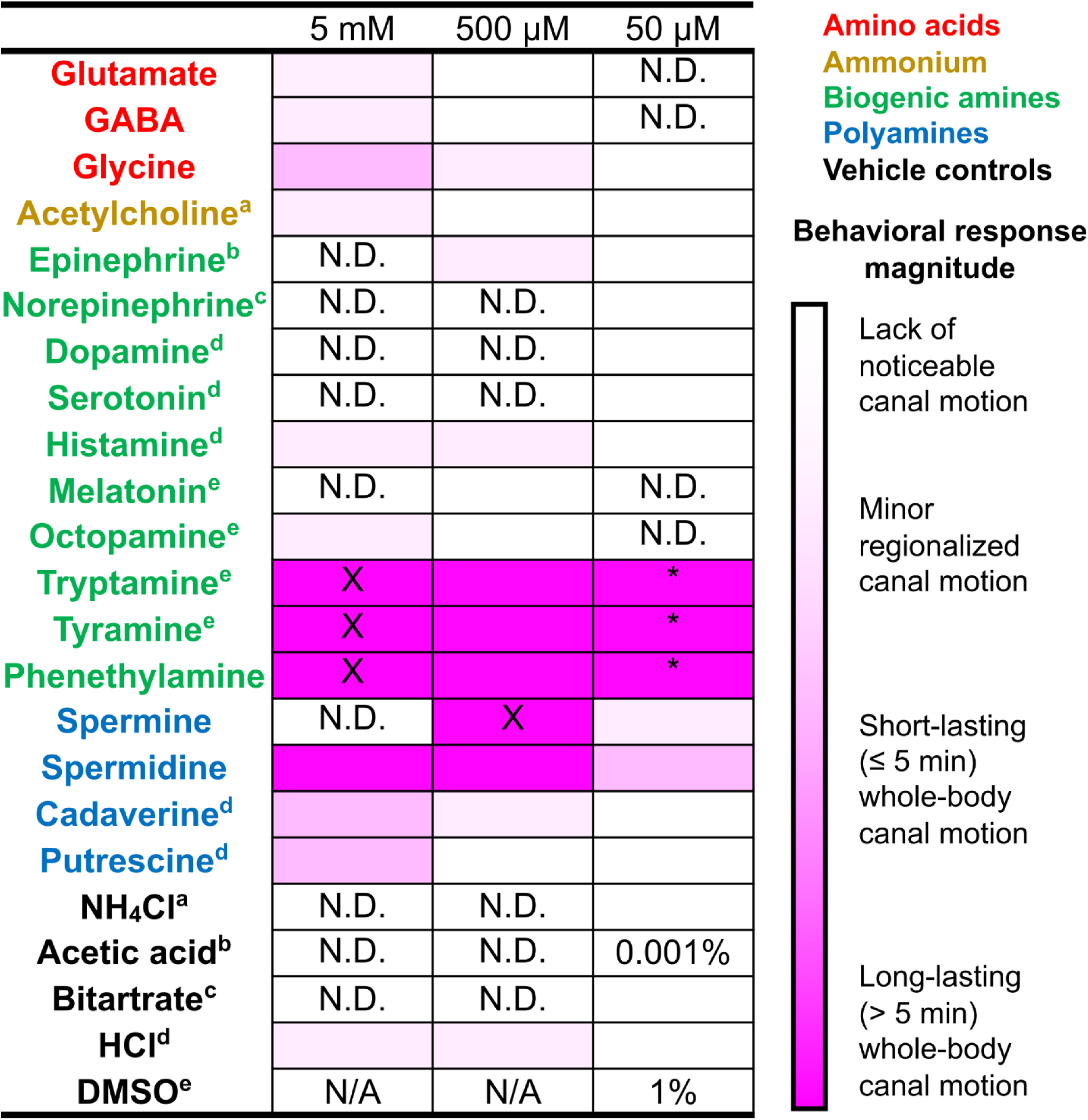
Behavioral screen of neurotransmitters and neuromodulators in *S. lacustris*. Heatmap summarizing behavioral responses of juvenile *S. lacustris* to bath application of amino acids, acetylcholine, monoamines, and polyamines across a range of concentrations (n ≥ 2). Color intensity indicates the magnitude of behavioral responses scored during the first hour following treatment (see Materials and Methods). Canonical neurotransmitters did not induce reproducible behaviors or elicited only subtle behaviors at millimolar concentrations, whereas tryptamine, phenethylamine, and tyramine elicited robust responses at micromolar concentrations. N.D., no detectable response. M-medium was used to dissolve compounds whenever possible. Superscript letters indicate alternative vehicle control used for solubility.

**Fig. S2.**
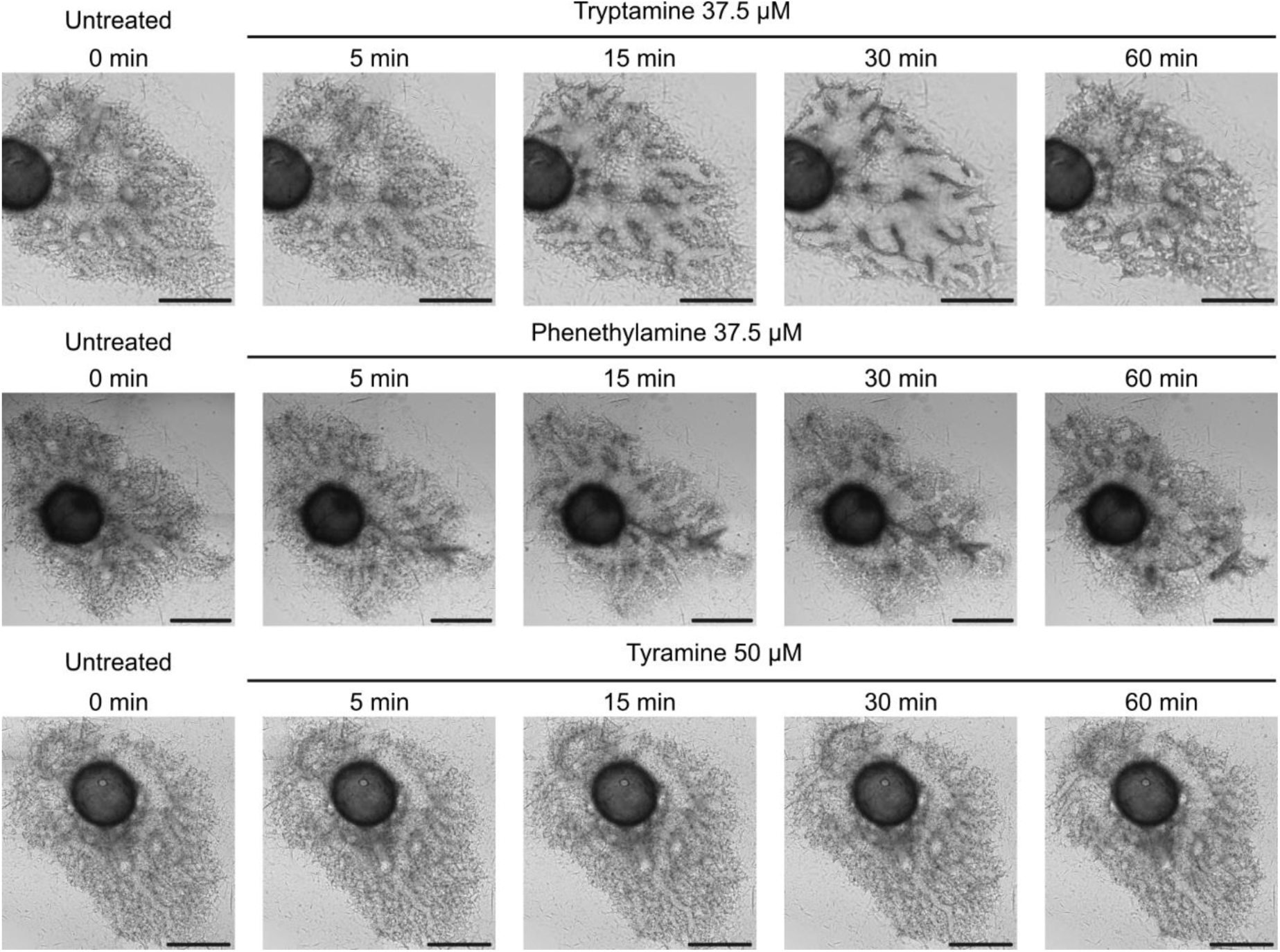
Overhead widefield imaging of decarboxylated monoamine responses in *S. lacustris*. Representative brightfield images of juvenile *S. lacustris* before and after treatment with tryptamine (37.5 µM), phenethylamine (37.5 µM), or tyramine (50 µM), showing distinct morphological changes. Overhead imaging provides qualitative visualization of gross morphological responses observable in two-dimensional projections. Scale bar, 500 μm.

**Fig. S3.**
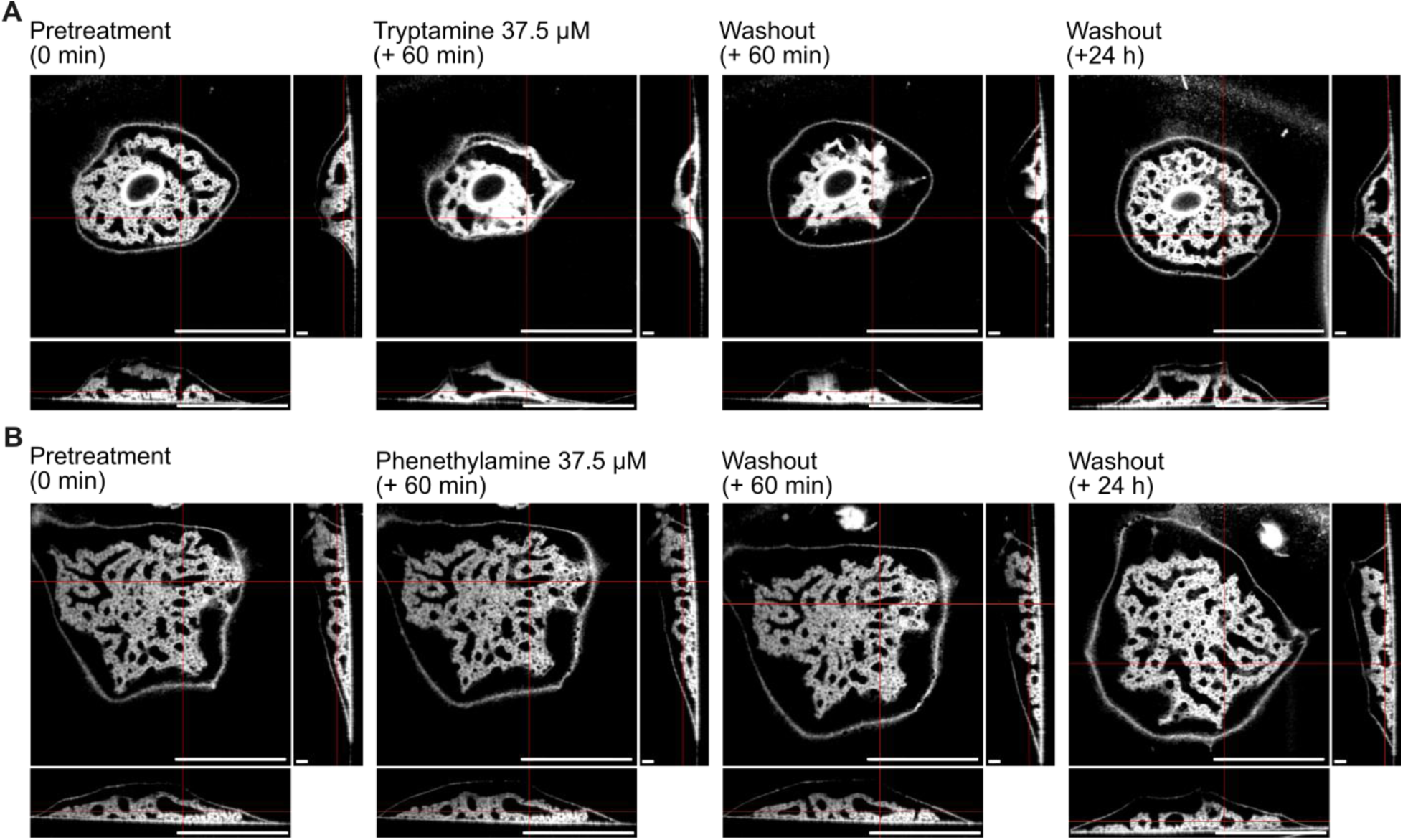
Reversibility of *S. lacustris* trace amine response following washout. **(A)** Orthogonal OCM views of juvenile sponges at four time points: pretreatment, 60 min after tryptamine treatment (37.5 µM), 60 min after washout, and 24 h after washout. Scale bar 1 mm (XY, XZ); 100 μm (ZY). **(B)** Orthogonal OCM views of juvenile sponges at four time points: pretreatment, 60 min after tryptamine treatment (37.5 µM), 60 min after washout, and 24 h after washout. Scale bar 1 mm (XY, XZ); 100 μm (ZY).

**Fig. S4.**
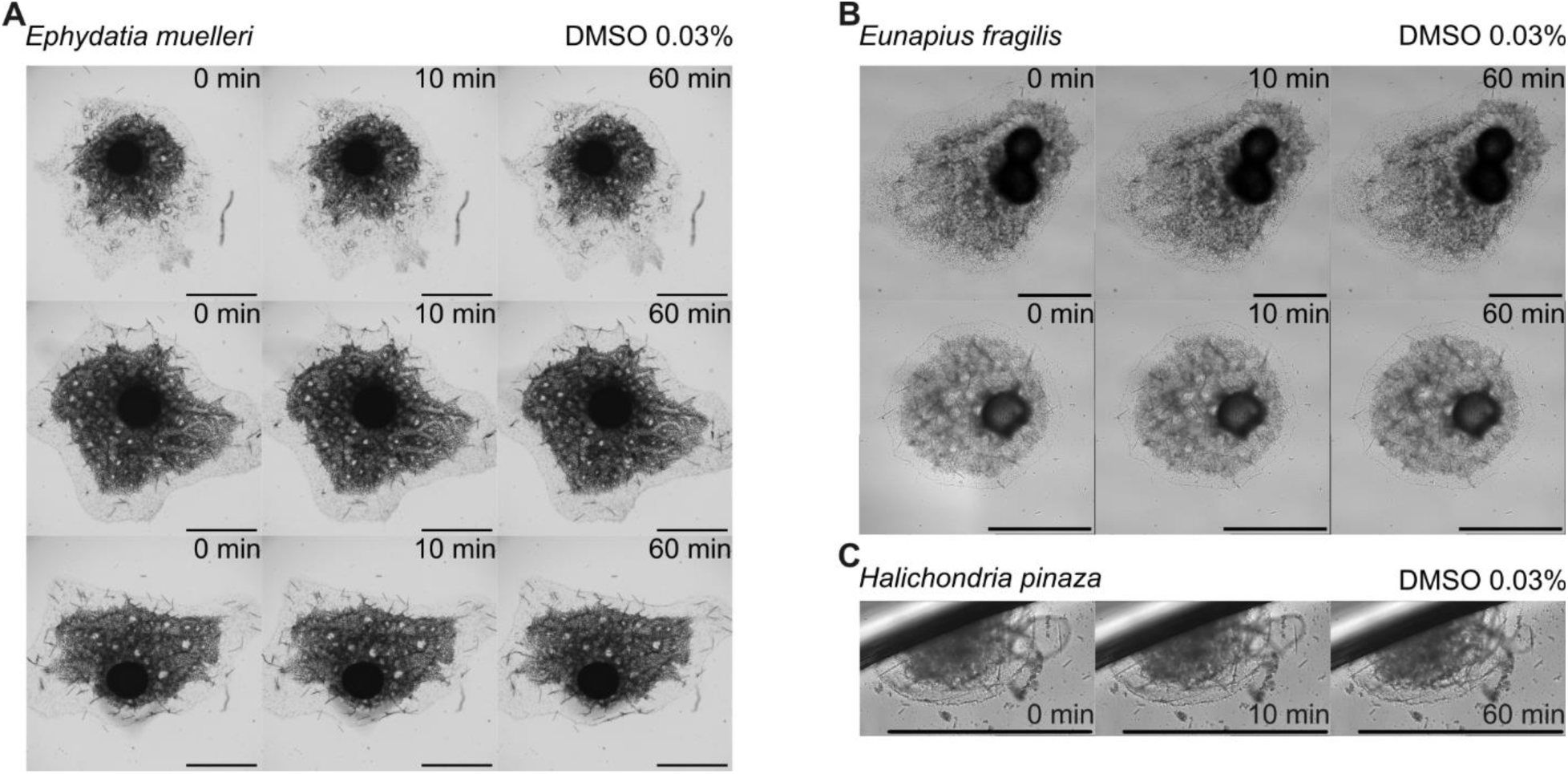
DMSO control treatments in freshwater and marine demosponges. **(A)** Brightfield images of *E. muelleri* treated with DMSO (0.03%) at three time points. Scale bars, 1 mm. **(B)** Brightfield images of *E. fragilis* treated with DMSO (0.03%) at three time points. Scale bars, 1 mm. **(C)** Brightfield images of *H. pinaza* treated with DMSO (0.03%) at three time points. Scale bars, 1 mm.

**Fig. S5.**
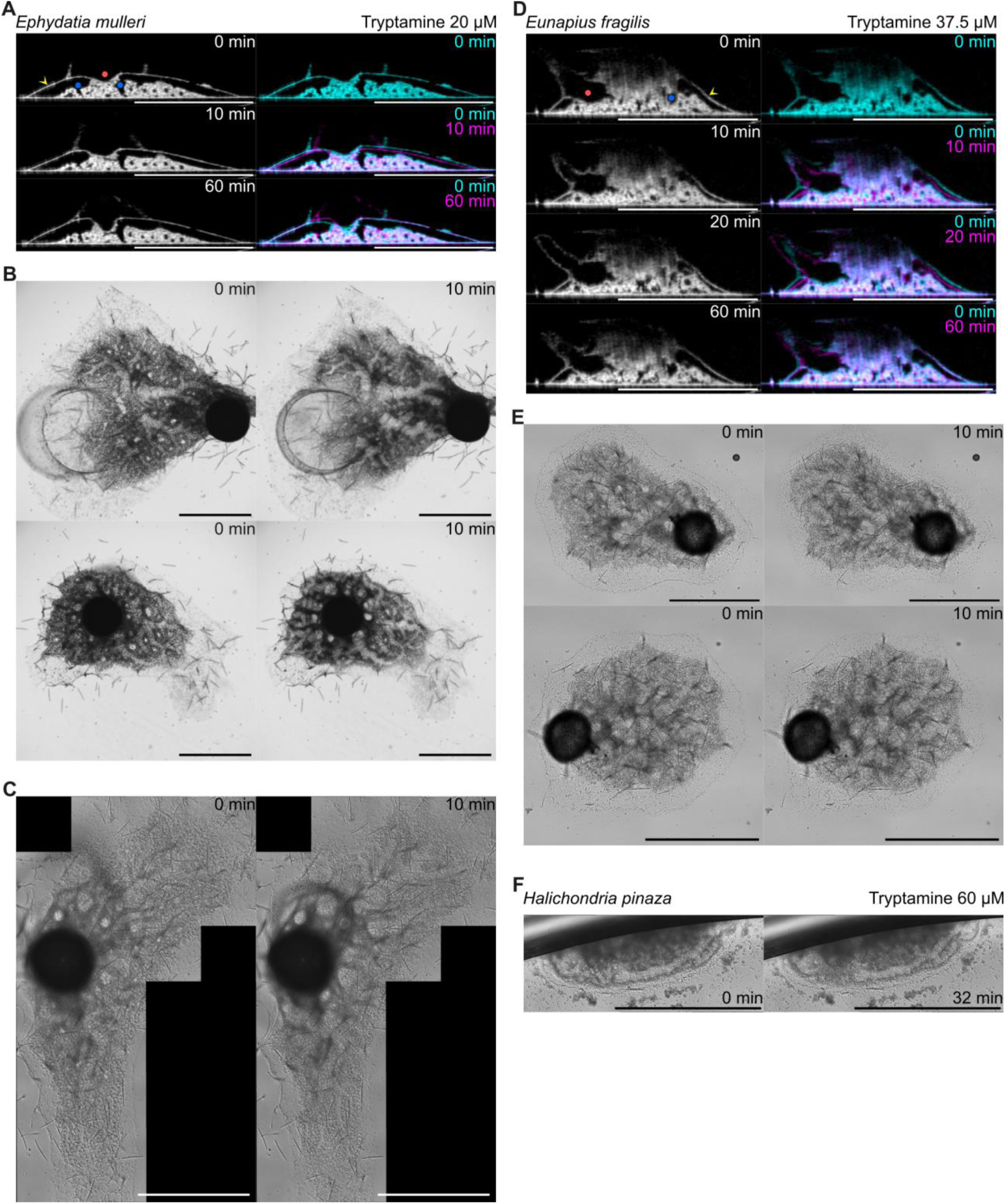
Tryptamine-induced morphological changes in demosponges. **(A)** Representative OCM cross-sections of *E. muelleri* before and after tryptamine treatment. Cyan and magenta indicate pretreatment (t = 0) and post-treatment time points (t = 10 min and t = 60 min). Markers: yellow arrowhead, outer epithelial tent; blue dots, incurrent canals; red dots, excurrent canals. Scale bar, 1 mm. **(B)** Brightfield images of *E. muelleri* treated with tryptamine (20 µM) across two time points. Scale bar, 1 mm. **(C)** DIC images of *E. muelleri* treated with tryptamine (20 µM) across two time points. Scale bar, 1 mm. **(D)** Representative OCM cross-sections of *E. fragilis* before and after tryptamine treatment. Cyan and magenta indicate pretreatment (t = 0) and post-treatment time points (t = 10 min, t = 20 min, and t = 60 min). Markers: yellow, outer epithelial tent; blue dots, incurrent canals; red dots, excurrent canals. Scale bar, 1 mm. **(E)** Brightfield images of *E. fragilis* treated with tryptamine (37.5 µM) across two time points. Scale bars, 1 mm. **(F)** DIC images of *H. pinaza* treated with tryptamine across two timepoints. Scar bar, 1 mm.

**Fig. S6.**
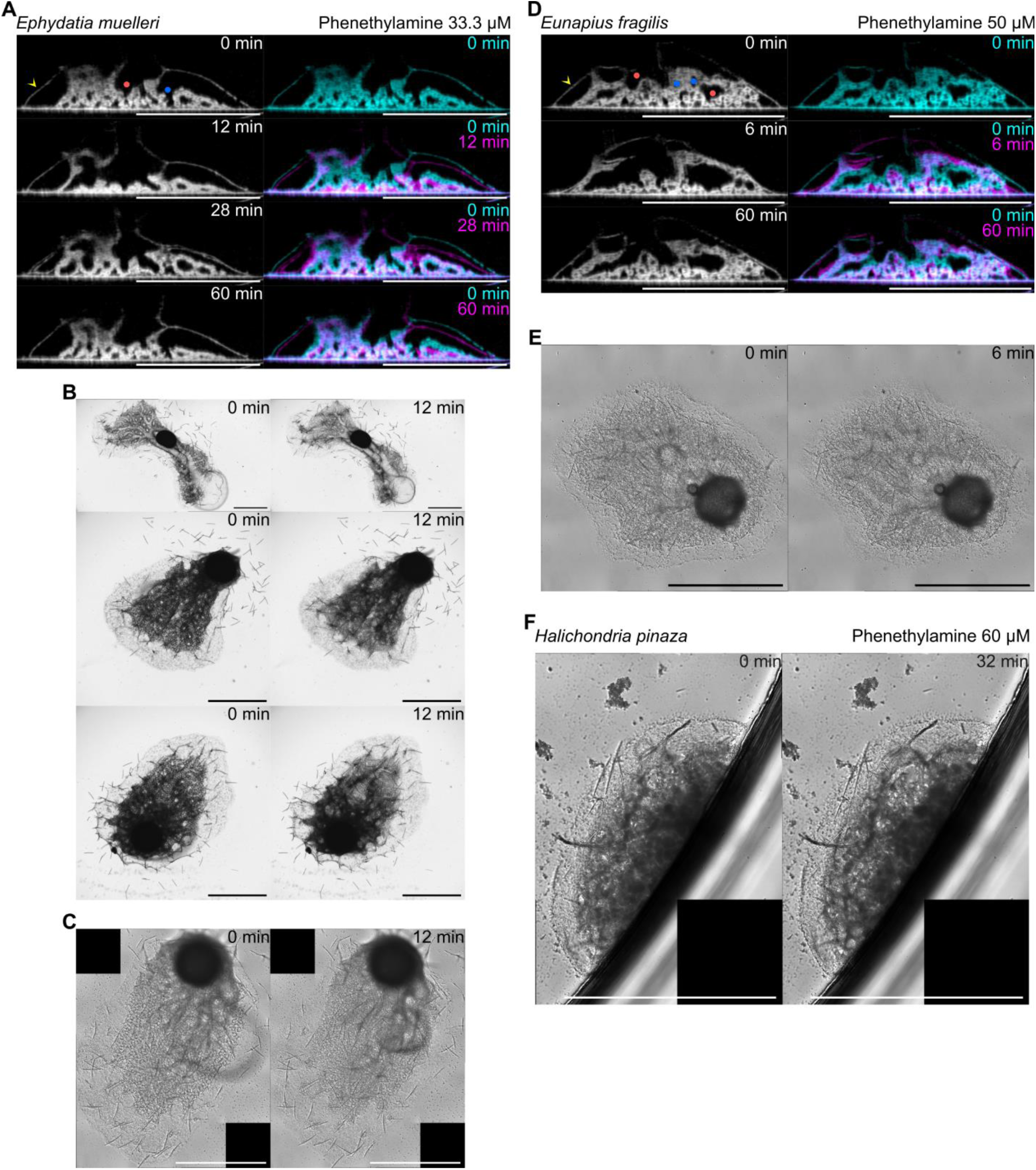
Phenethylamine-induced morphological changes in demosponges. **(A)** Representative OCM cross-sections of *E. muelleri* before and after tryptamine treatment. Cyan and magenta indicate pretreatment (t = 0) and post-treatment time points (t = 12 min, t = 28 min, and t = 60 min). For the grayscale pretreatment image, yellow arrowhead points at the outer epithelial tent, blue dots represent incurrent canals, and red dots represent excurrent canals. Scale bar, 1 mm. **(B)** Brightfield images of *E, muelleri* treated with phenethylamine (20 µM) across two time points. **(C)** DIC images of *E. muelleri* treated with phenethylamine (20 µM) across two time points. **(D)** Representative OCM cross-sections of *E. fragilis* before and after tryptamine treatment. Cyan and magenta indicate pretreatment (t = 0) and post-treatment time points (t = 6 min and t = 60 min). Markers: yellow arrowhead, outer epithelial tent; blue dots, incurrent canals; red dots; excurrent canals. Scale bar, 1 mm. **(E)** Brightfield images of *E. fragilis* treated with phenethylamine (37.5 µM) across two time points. **(F)** DIC images of *H. pinaza* treated with phenethylamine across two time points. Scale bars, 1 mm.

**Fig. S7.**
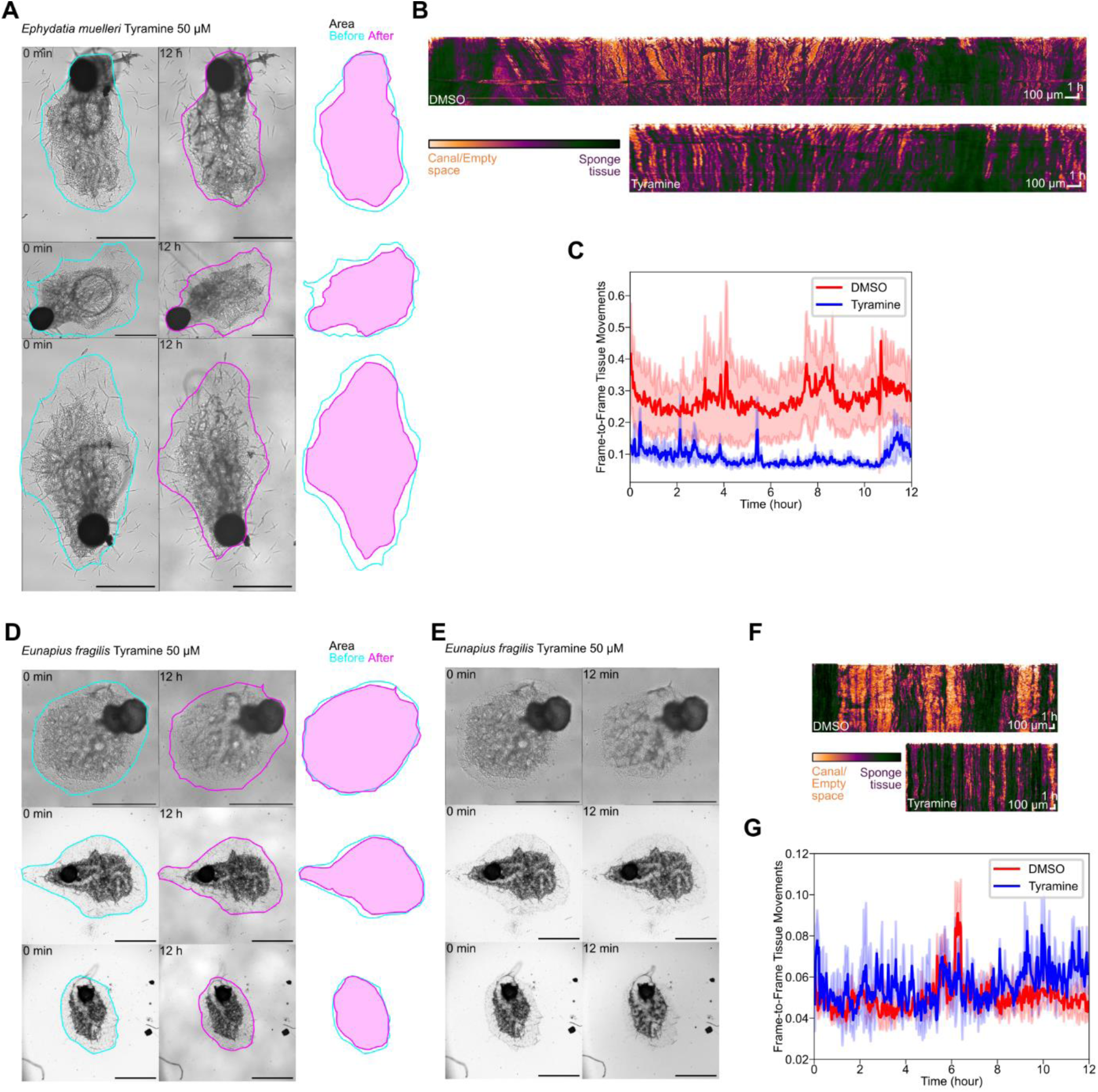
Tyramine-induced morphological and activity changes in demosponges. **(A)** Brightfield images of *E. muelleri* before (cyan) and after (magenta) 12 h tyramine treatment, with traced outlines of outer tissue overlaid on the right. Scale bar, 1 mm. **(B)** Canal activity visualized by line-scan kymographs from DMSO- and tyramine-treated *E. mulleri*, generated as illustrated in Fig. 1K. **(C)** Canal activity and tissue movements of *E. mulleri* quantified as dissimilarity of pixels (Pearson’s distance, see Materials and Methods) over time following tyramine (n = 3) or DMSO treatment (n = 3). Dissimilarity reflects frame-to-frame intensity changes along canal structures. **(D)** Brightfield images of *E. fragilis* before (cyan) and after (magenta) 12 h tyramine treatment, with traced outlines of outer tissue overlaid on the right. Scale bar, 1 mm. **(E)** Brightfield images of *E. fragilis* treated with tyramine (50 µM) across two time points. Scale bar, 1mm. **(F)** Canal activity visualized by line scan kymographs from DMSO- and tyramine-treated *E. fragilis* generated as illustrated in Fig. 1K. **(G)** Canal activity and tissue movements of *E. fragilis* quantified as dissimilarity of pixels (Pearson’s distance, see Materials and Methods) over time following tyramine (n = 3) or DMSO treatment (n = 3). Dissimilarity reflects frame-to-frame intensity changes along canal structures.

**Fig. S8.**
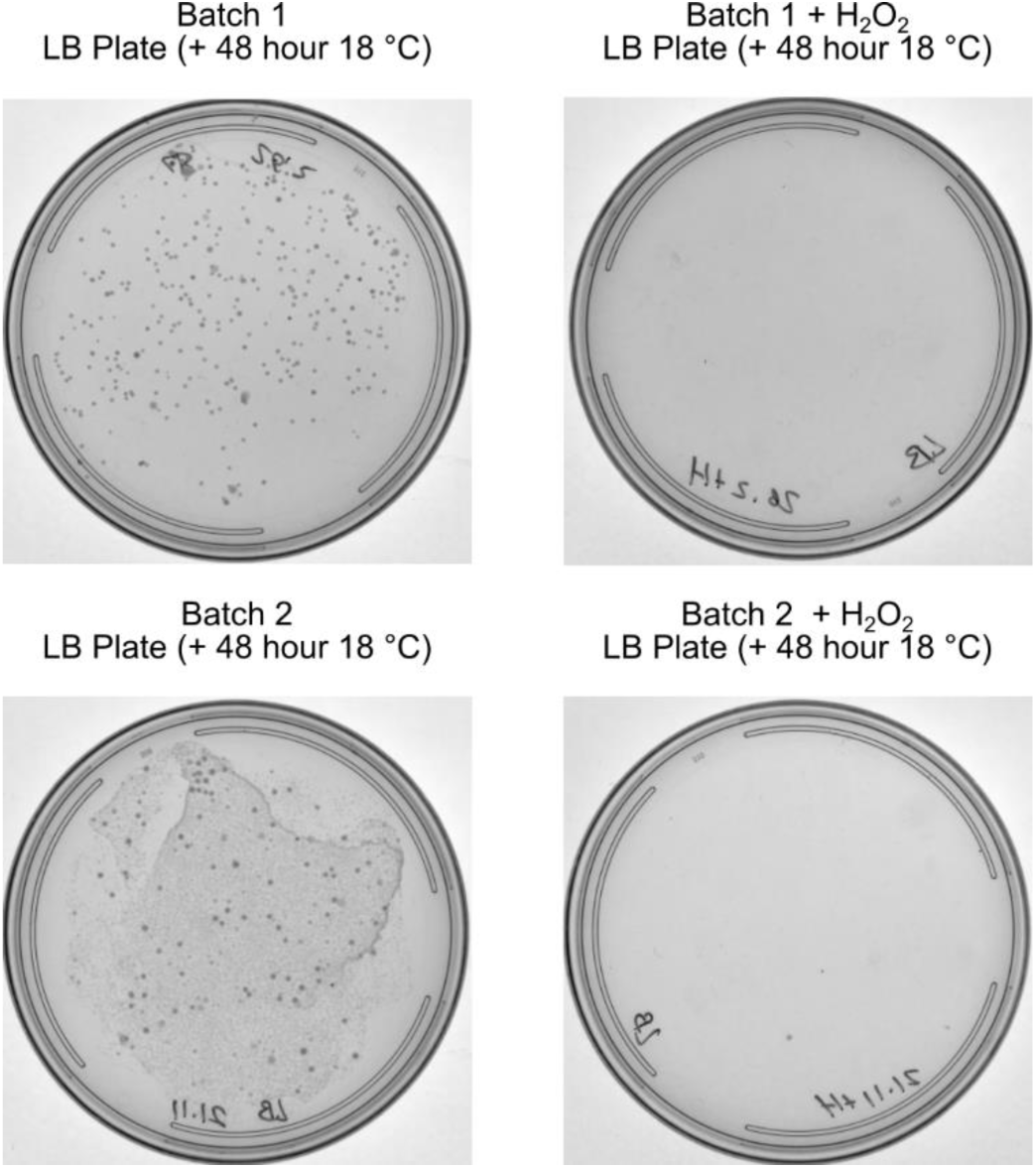
*S. lacustris* has reduced bacterial load after H_2_O_2_ surface sterilization. Pictures of lysogeny broth (LB) agar plates streaked with mashed extract of *S. lacustris* gemmules with and without H_2_O_2_ surface sterilization. Two different batches were used as biological replicates.

**Fig. S9.**
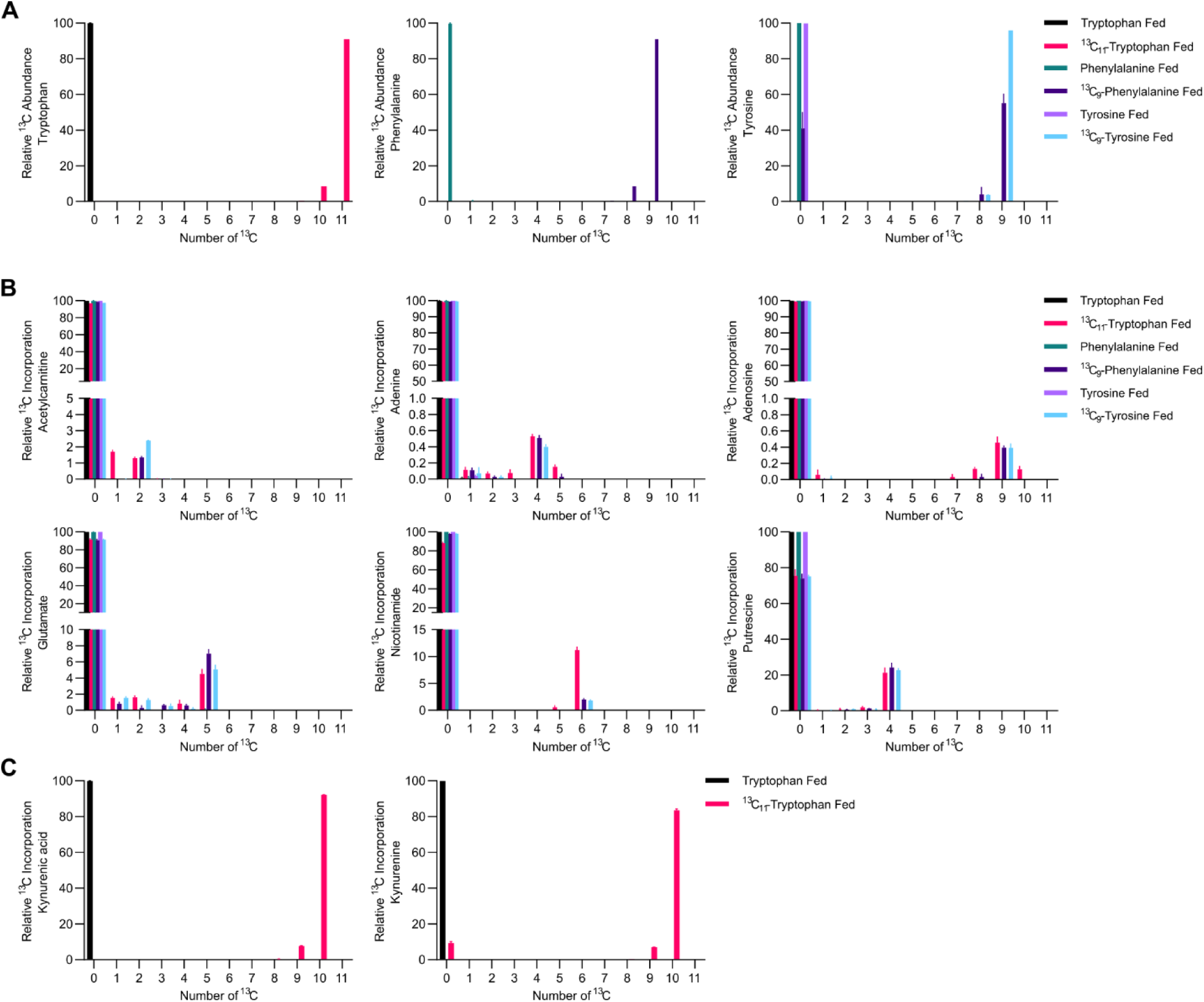
Compounds synthesized *in S. lacustris* from amino acid precursors. **(A)** Bar plots of high-res heavy isotope tracing showing relative incorporation of ^13^C into different amino acids. **(B)** Bar plots of high-res heavy isotope tracing showing relative incorporation of ^13^C into different metabolites from all non-heavy or ^13^C amino acid fed sponges. **(C)** Bar plots of high-res heavy isotope tracing showing relative incorporation of ^13^C into different kynurenic acid and kynurenine from all non-heavy or ^13^C tryptophan fed sponges.

**Fig. S10.**
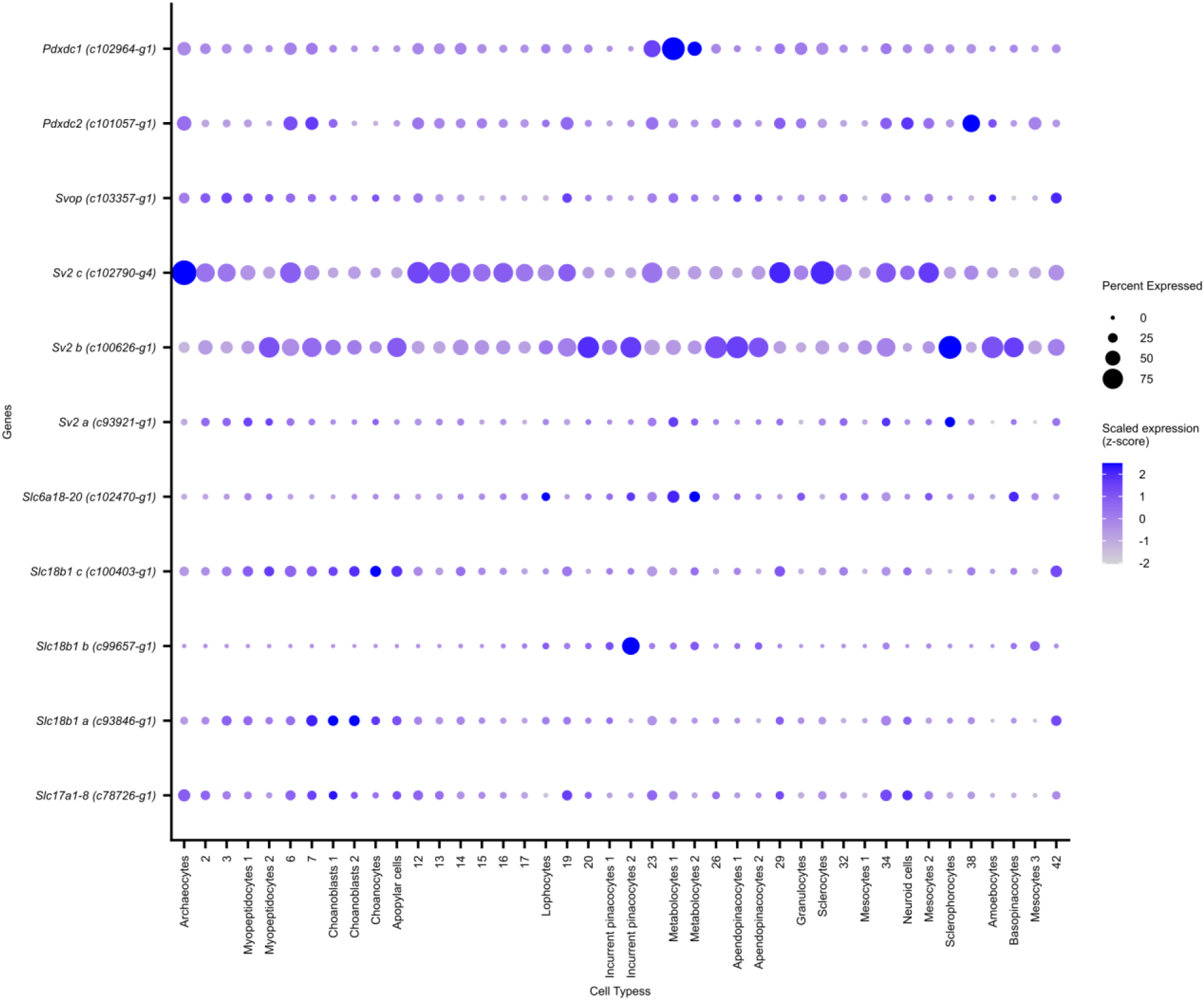
Dot plot of *S. lacustris* monoamine decarboxylase and vesicular transporters.

**Fig. S11.**
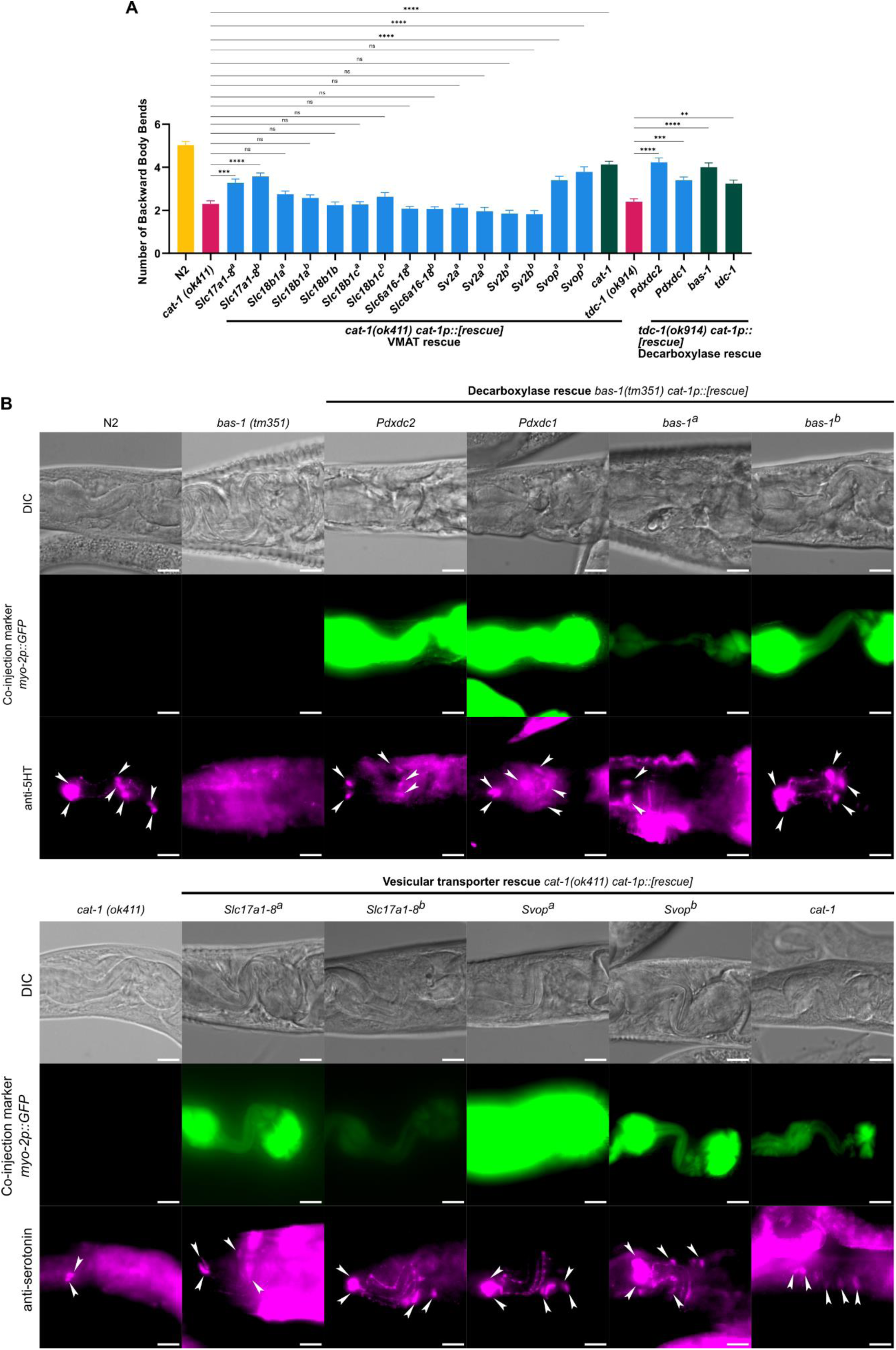
*C. elegans* monoamine rescue. **(A)** Bar graph quantification of backward body bends in response to anterior touch of wild-type (WT) animals (yellow, N2, n = 22), *Vmat* mutants (magenta, *cat-1(ok411),* n = 25), and *Tdc-1* mutants (magenta, *tdc-1(ok914),* n = 25). Expression of *S. lacustris* (blue) *Slc17a1-8* (strain a n = 27; strain b n = 27), *Svop* (strain a n = 25; strain b n = 23), *Pdxdc2* (n = 26), *Pdxdc1* (n = 25) and *C.* elegans (green) *Vmat* (*cat-1,* n = 28), *Aadc* (*bas-1*, n = 24), *Tdc-1* (*tdc-1*, n = 25) significantly increased the number of backward body bends compared to *Vmat* or *Tdc-1* mutants. Expression of *S. lacustris* (blue) *Slc18b1a* (strain a, n = 28, strain b, n = 28), *Slc18b1b* (n = 25), *Slc18b1c* (strain a, n = 24, strain b, n = 23), *Slc6a16-18* (strain a, n = 20, strain b, n = 17), *Sv2a* (strain a, n = 25, strain b, n = 23), and *Sv2b* (strain a, n = 23, strain b, n = 22) did not significantly increase the number of backward body bends compared to *Vmat* or *Tdc-1* mutants. Nonmatching one-way ANOVA followed by Šídák test with a 95% confidence interval (α = 0.05); ns = not significant P ≥ 0.05; **P < 0.01; *** P < 0.001; **** P < 0.0001; bars on scatter plot show mean with standard errors of the mean. **(B)** Representative images of adult WT (N2 strain), mutants, and transgenic *C. elegans* stained with anti-serotonin antibody. Arrowheads represent serotonin-positive neurons. WT worms have seven serotonin positive neurons near the head (serotonin synthesis: 2 NSMs and 2 ADFs, serotonin uptake: RIH and 2 AIMs). For decarboxylase rescue, all serotonin positive neurons were counted. For vesicular transporter rescue, any serotonin positive neurons in addition to 2 NSMs were counted. Scale bar 10 µm.

**Fig. S12.**
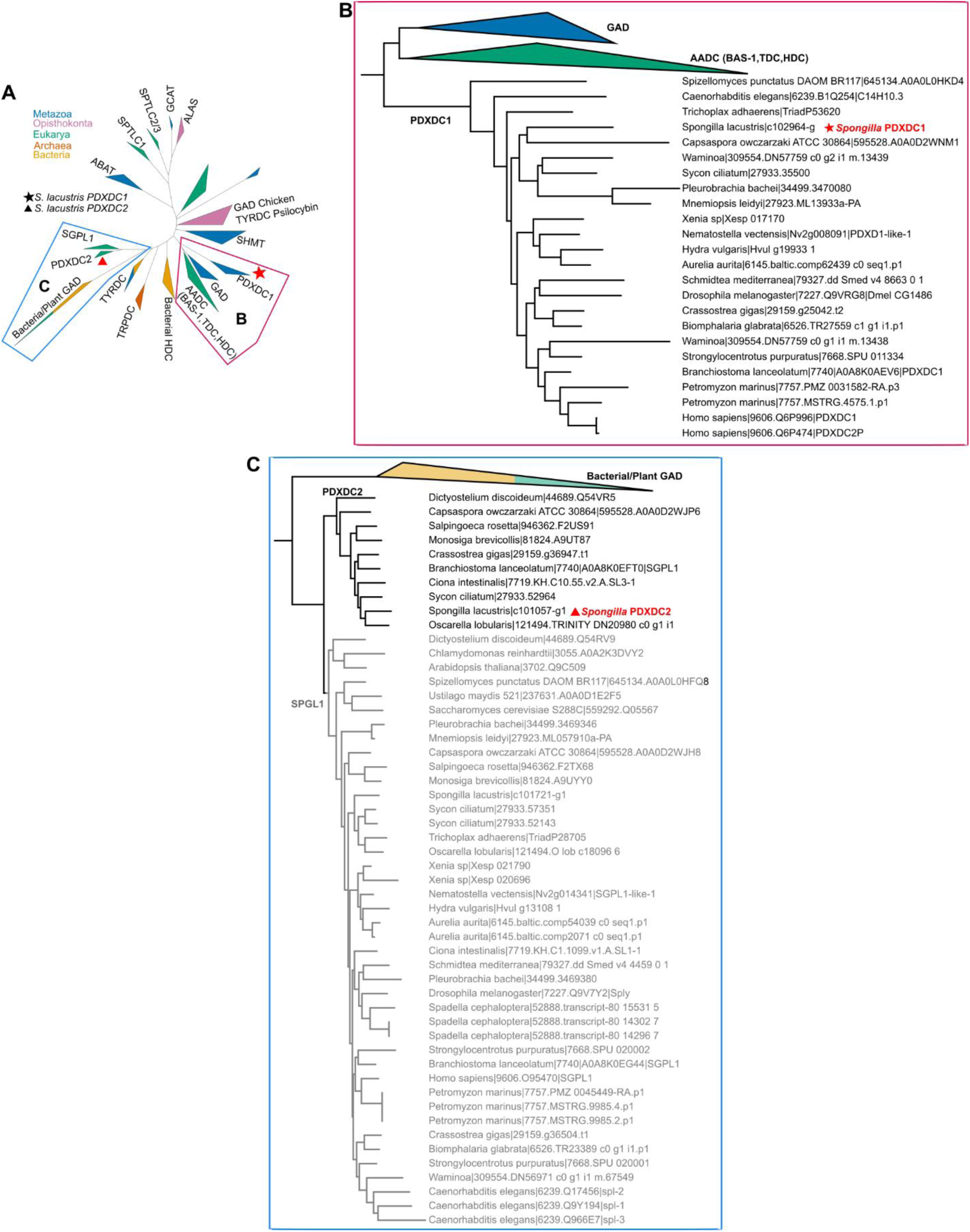
PLPαF Decarboxylase family phylogeny. **(A)** Maximum-likelihood phylogenetic tree of PLPαF decarboxylases found. Blue and magenta boxes indicate expanded trees in panels B and C. **(B)** Expanded region of PLPαF decarboxylase maximum-likelihood phylogenetic tree showing PDXDC1 orthology group in animals. **(C)** Expanded region of PLPαF decarboxylase maximum-likelihood phylogenetic tree showing PDXDC2 orthology group with eukaryotic origin and subsequent loss in some animal lineages.

**Fig. S13.**
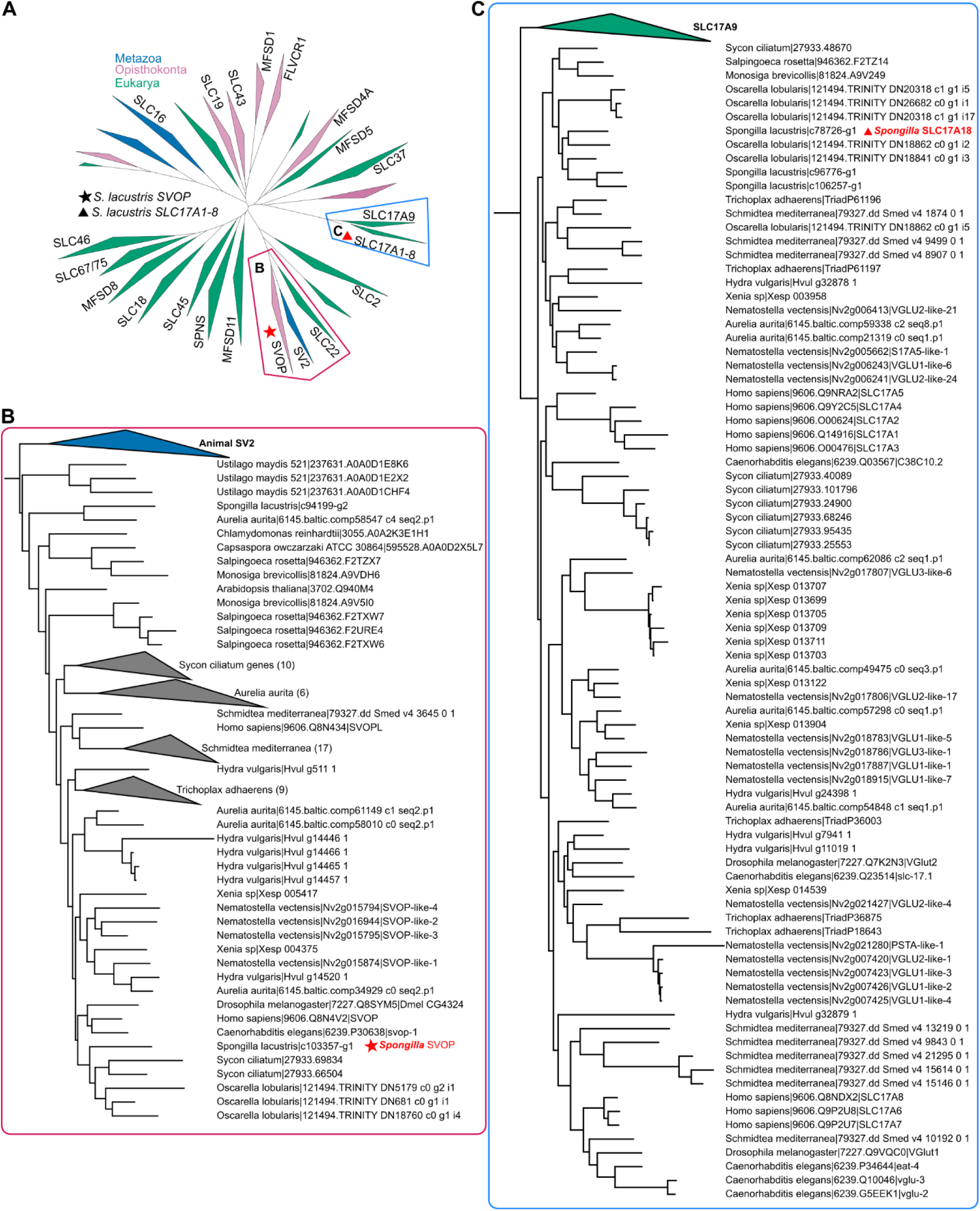
MFS family transporter phylogeny. **(A)** Maximum-likelihood phylogenetic tree of MFS vesicular transporters. Blue and magenta boxes show expanded trees in panels B and C. **(B)** Expanded region showing SVOP orthology group is conserved in animals with additional lineage specific duplication in *Sycon, Aurelia, Schmidtea,* and *Trichoplax*. Number in parenthesis indicates the number of genes within the collapsed clade. **(C)** Expanded region of maximum-likelihood MFS phylogenetic tree showing *S. lacustris* SLC17A1-8 is in a clade sister to human SLC17A1 to 8 genes.

**Fig. S14.**
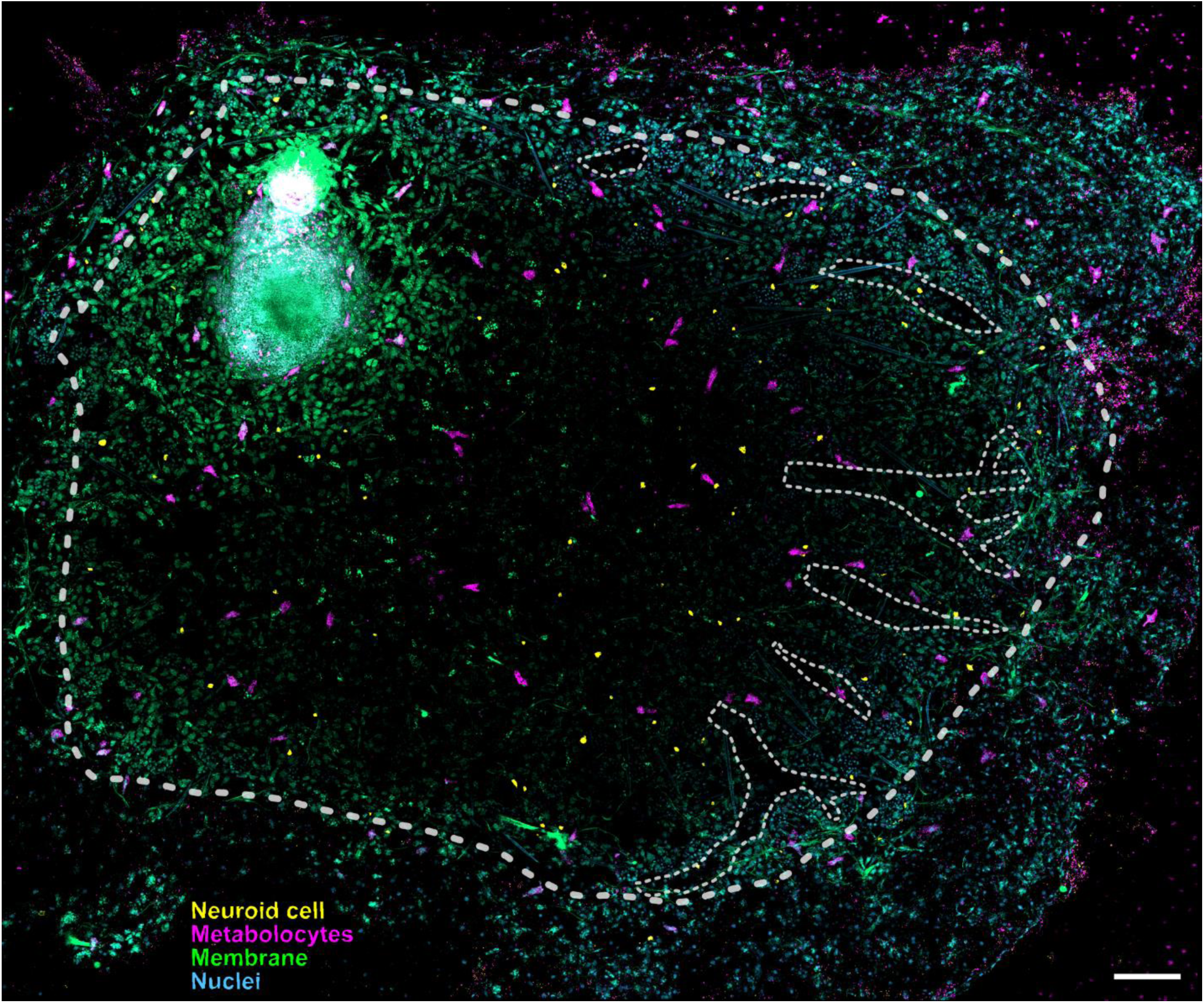
*S. lacustris* monoaminergic secretory cells are differentially distributed across the body. Maximum Z projection of whole-body HCR staining of metabolocytes and neuroid cell type expression markers. Large, dashed line indicates the outer border of the sponge tissue and small dashed lines indicate outline of visible canal systems. Scale bar, 1 mm.

**Fig. S15.**
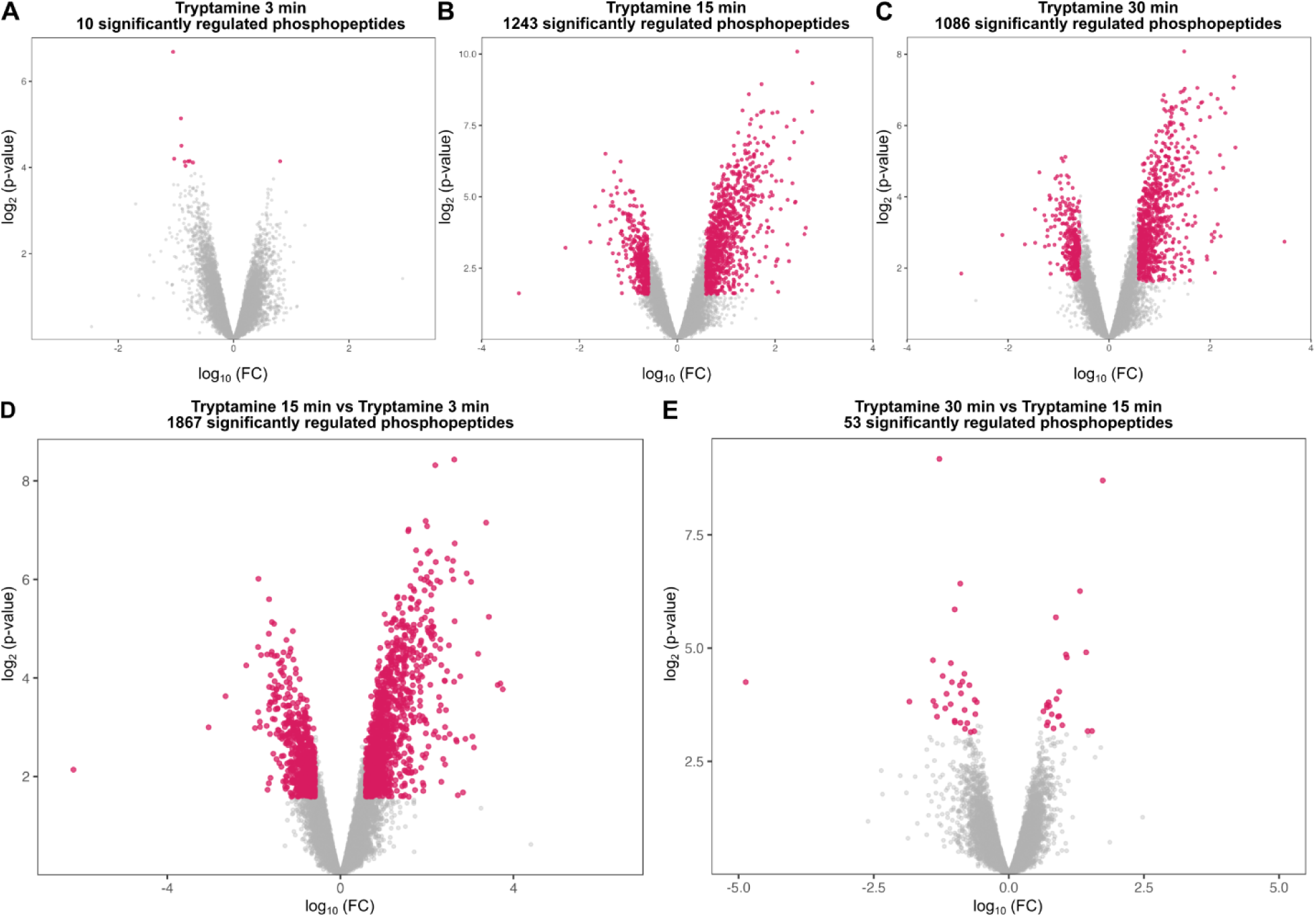
Volcano plots illustrating quantitative differences in phosphopeptides between tryptamine (37.5 µM)-treated or DMSO-treated *S. lacustris*. Phosphorylation differences are quantified for unique phosphosites relative to abundance in DMSO controls. The x-axis shows log2 FC of tryptamine/DMSO. The y-axis represents -log₁₀ p-values. Colored dots indicate significantly regulated phosphopeptides.

**Fig. S16.**
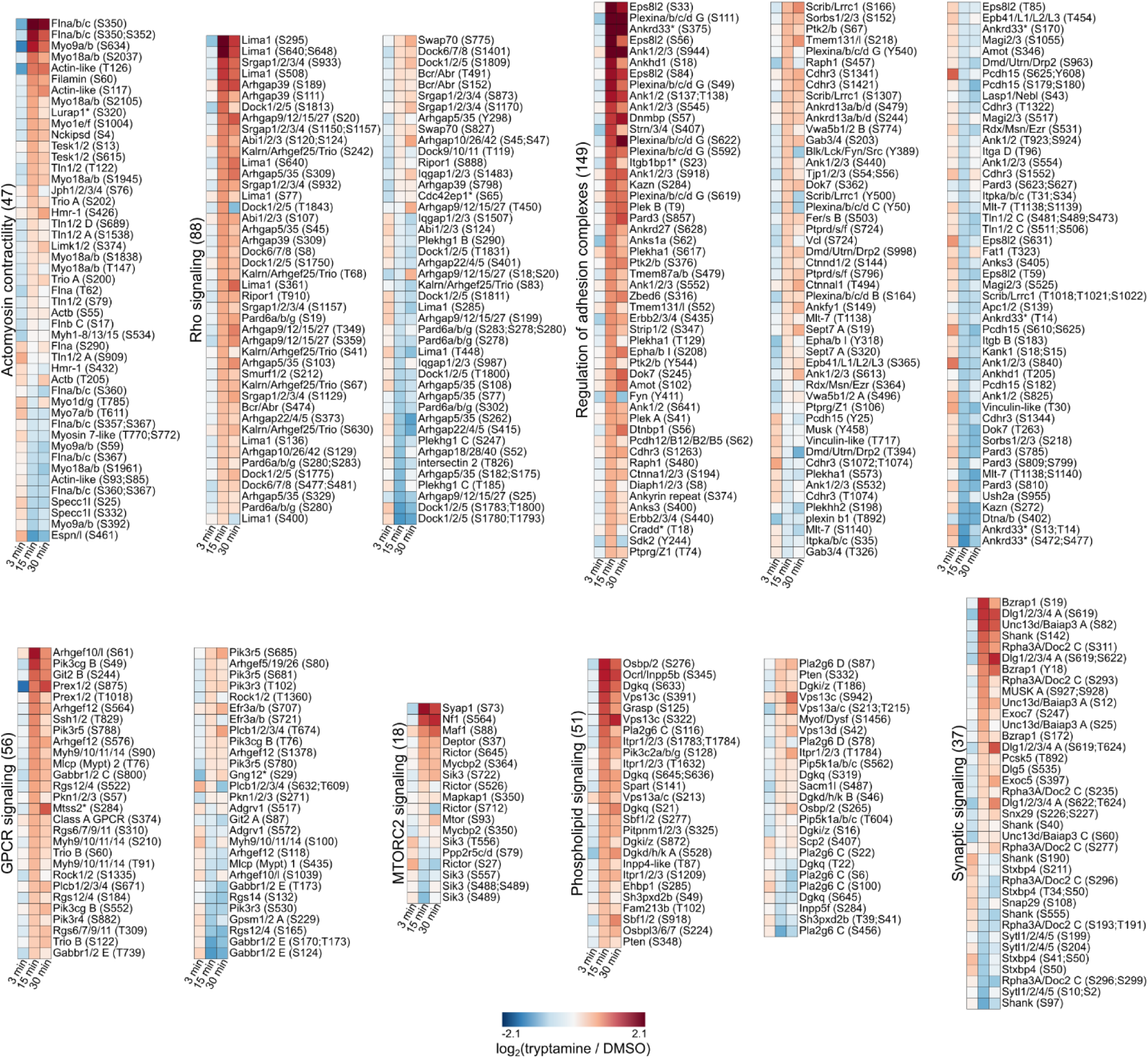
Heatmaps of significantly regulated phosphopeptides for proteins related to molecular modules of interest in tryptamine-treated *S. lacustris*. Phosphosite FC calculated relative to timepoint matched DMSO-treated controls.

**Fig. S17.**
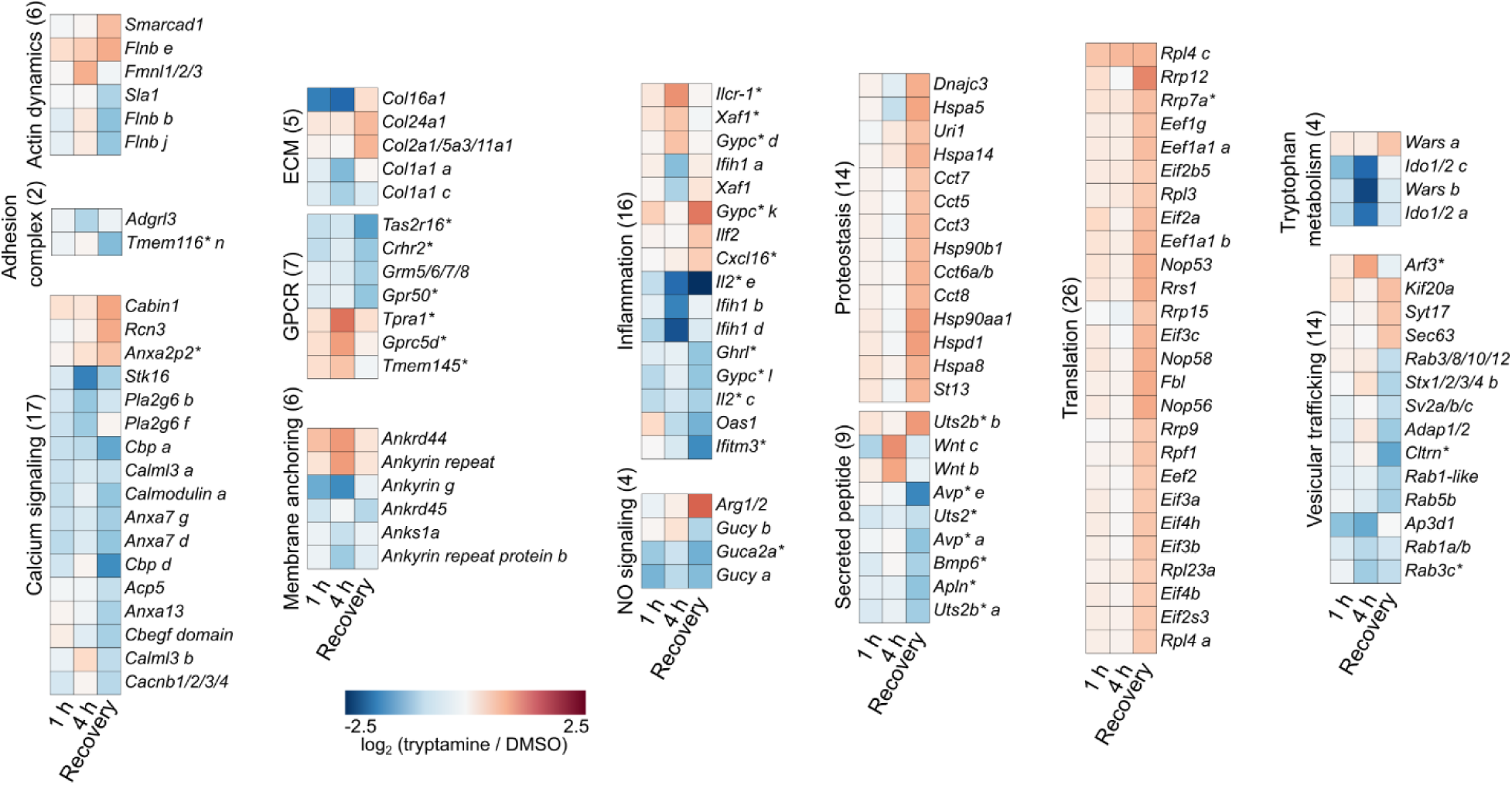
Heatmaps of differentially expressed genes categorized by molecular modules of interest after tryptamine treatment of *S. lacustris.* Color represents fold change in transcript counts between time-matched tryptamine and DMSO-treated samples.

**Fig. S18.**
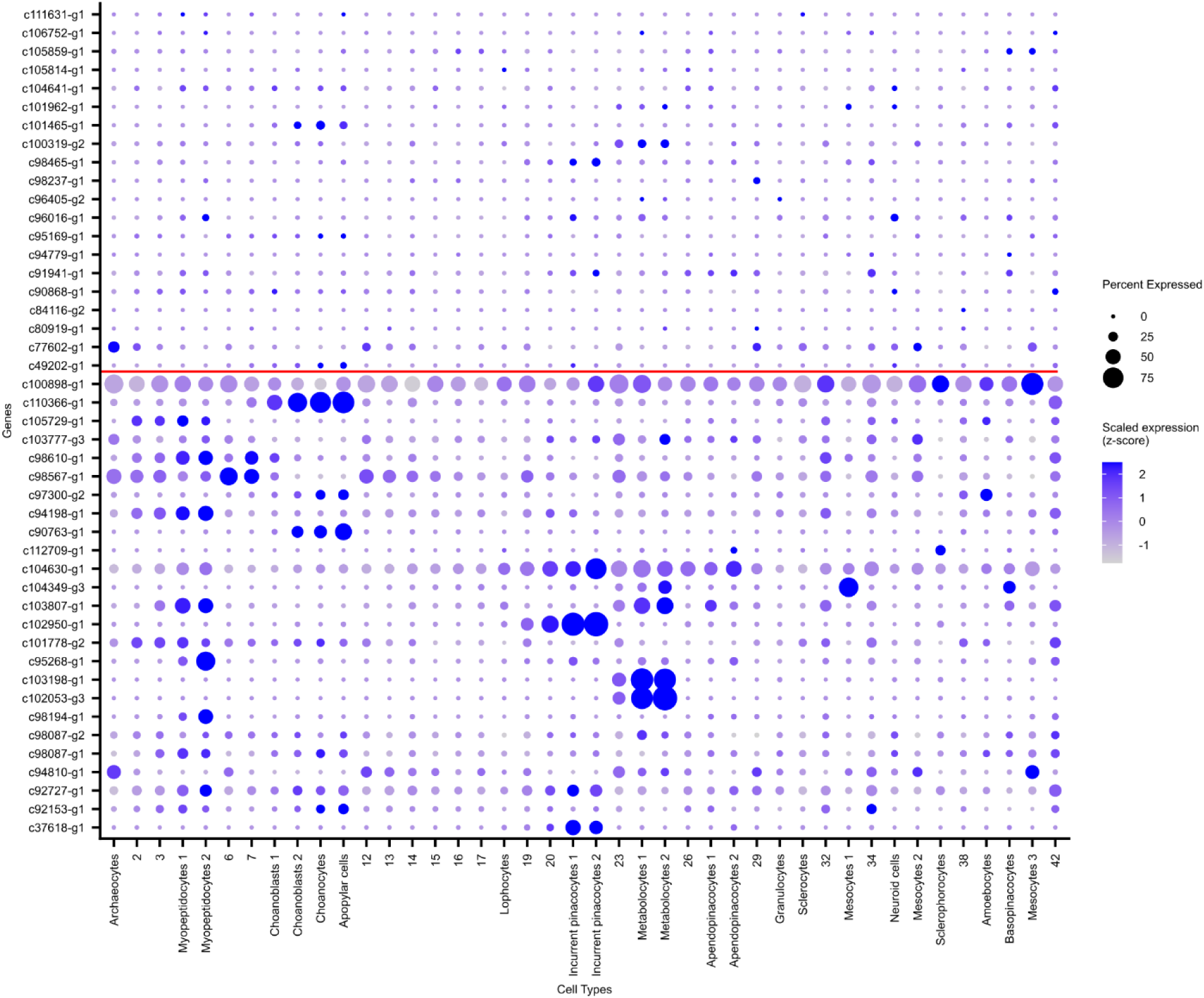
Dot plot of *S. lacustris* GPCRs. *S. lacustris* GPCRs with complete seven transmembrane domains were identified using human class A and C GPCR with *Dictyostelium* class E GPCR as query sequences. GPCRs with cell type-specific expression (beneath the red line) were subjected to further docking analysis.

**Fig. S19.**
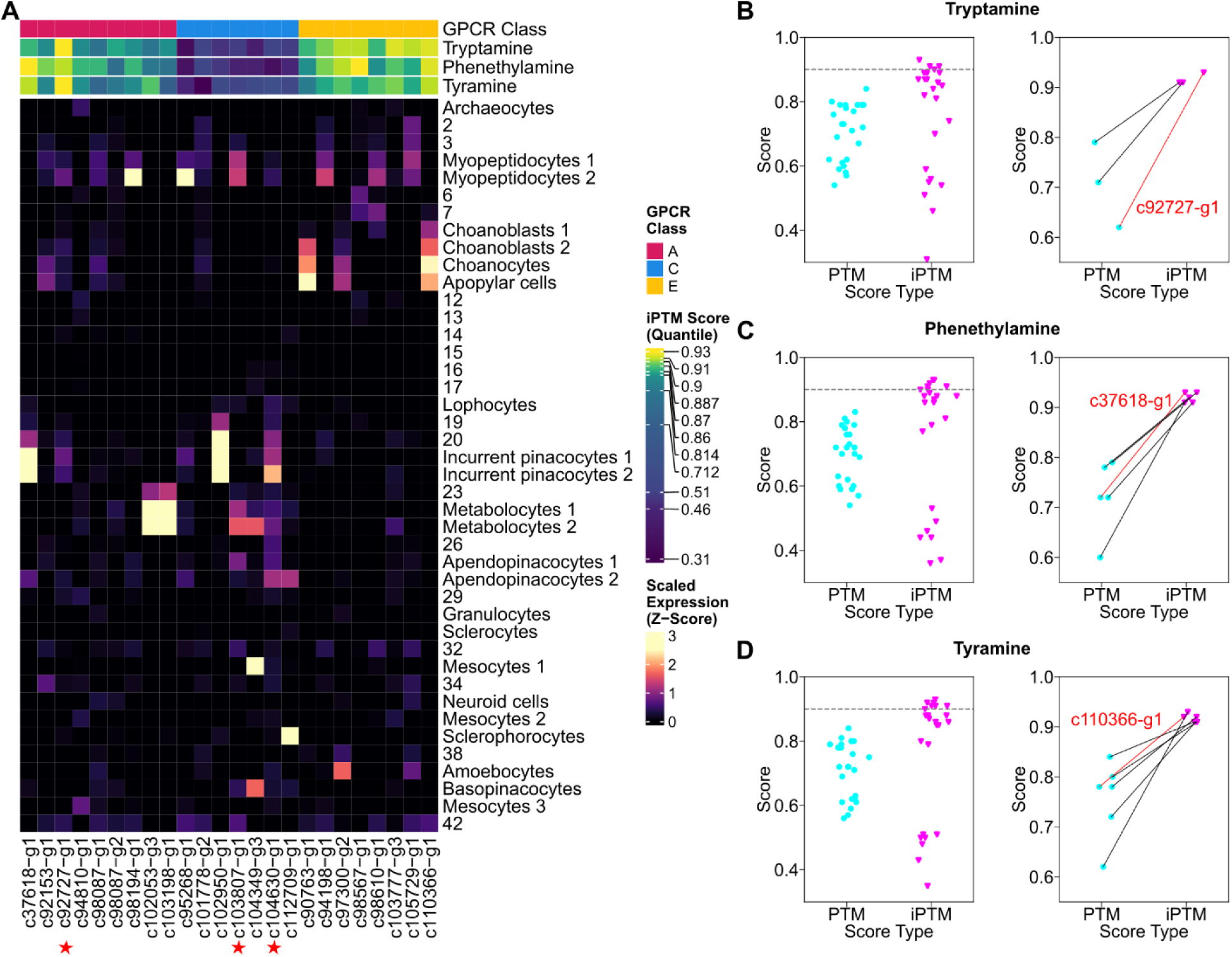
AF3 docking analysis of *S. lacustris* GPCR along with cell type expression. **(A)** Heatmap of *S. lacustris* GPCR scaled expression across all cell types. GPCRs are separated by different classes and labeled with interface predicted TM-Score when docked with tyramine, phenethylamine, and tryptamine using AF3. Color bars represent GPCR classes, iPTM scores for docking with tryptamine, phenethylamine, and tyramine. Red stars represent GPCRs phosphorylated upon tryptamine treatment. **(B)** Left: Scatter plot of PTM scores (receptor structure prediction) of GPCRs and iPTM scores (receptor-ligand interface prediction) for GPCRs docked with tryptamine using AF3. Grey dashed line represents iPTM cut-off (>0.9) used for visualization of PTM-iPTM connection plot on the right. Right: Zoomed in scatter plot of PTM and iPTM scores. Lines represent the connections of PTM and iPTM score for the same receptor. Red line highlights the receptor considered as most likely candidate for tryptamine receptor with AF3 docking analysis. The docked structure is visualized in Figure 5B. **(C)** Left: Scatter plot of PTM scores (receptor structure prediction) of GPCRs and iPTM scores (receptor-ligand interface prediction) for GPCRs docked with phenethylamine using AF3. Grey dashed line represents iPTM cut-off (>0.9) used for visualization of PTM-iPTM connection plot on the right. Right: Zoomed in scatter plot of PTM and iPTM scores. Lines represent the connections of PTM and iPTM score for the same receptor. Red line highlights the receptor considered as most likely candidate for phenethylamine receptor with AF3 docking analysis. The docked structure is visualized in Figure S21A. **(D)** Left: Scatter plot of PTM scores (receptor structure prediction) of GPCRs and iPTM scores (receptor-ligand interface prediction) for GPCRs docked with tyramine using AF3. Grey dashed line represents iPTM cut-off (>0.9) used for visualization of PTM-iPTM connection plot on the right. Right: Zoomed in scatter plot of PTM and iPTM scores. Lines represent the connections of PTM and iPTM score for the same receptor. Red line highlights the receptor considered as most likely candidate for tyramine receptor with AF3 docking analysis. The docked structure is visualized in Figure 21B.

**Fig. S20.**
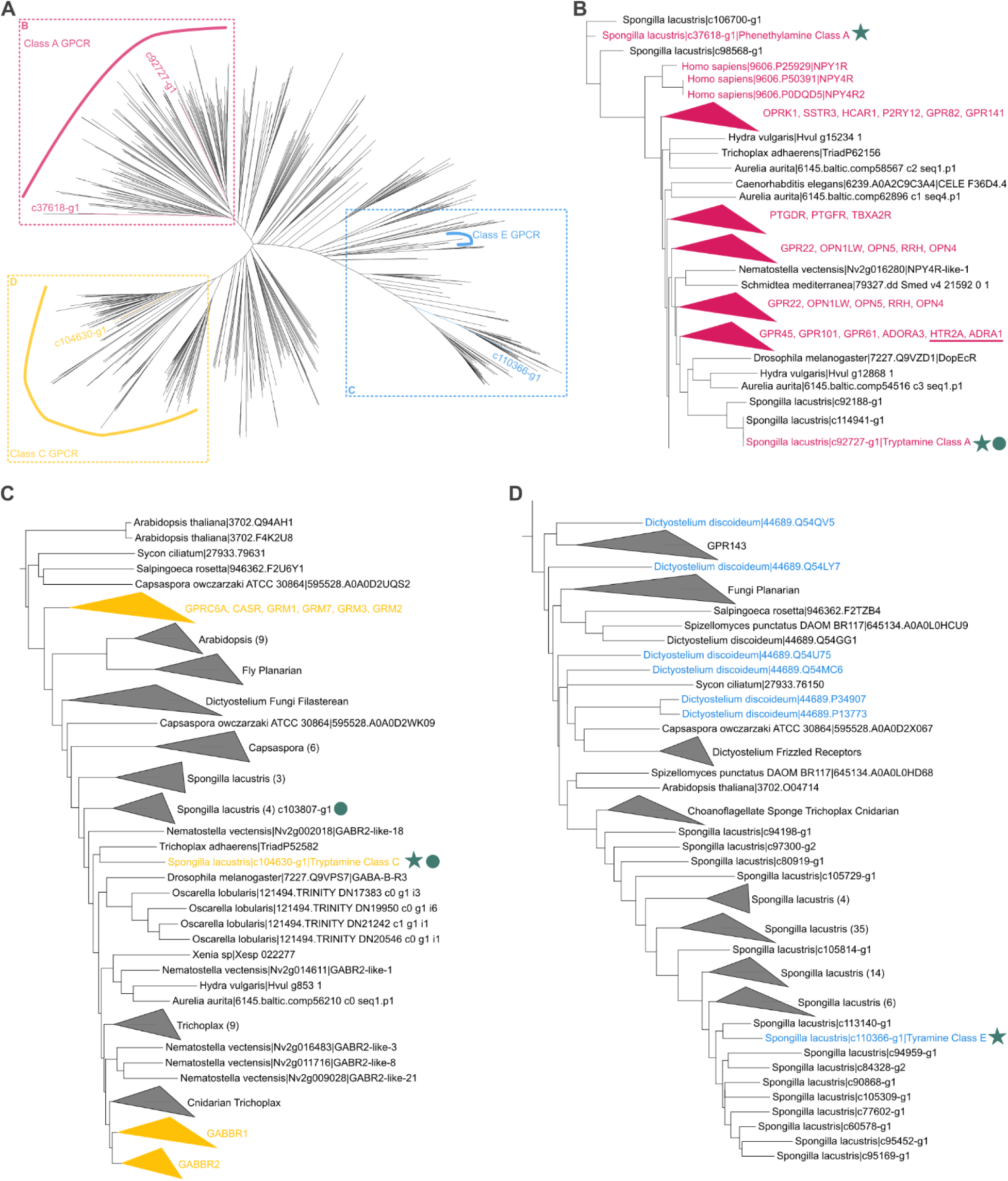
*S. lacustris* monoamine GPCR belongs to distinct GPCR classes. **(A)** Unrooted maximum likelihood tree of BLAST and PROST hits for *S. lacustris* GPCRs with high iPTM (>0.90) scores when docked with decarboxylated monoamines. GPCRs with the highest potential of binding to tryptamine, phenethylamine, and tyramine are colored according to their phylogenetic positioning with known GPCR classes. **(B)** Overview of *S. lacustris* tryptamine class A GPCR and phenethylamine class A GPCR in relationship to human class A GPCRs including (neuro)peptide receptors, opsins, and monoamine receptors (magenta). Star highlights the highest likely candidate for tryptamine and phenethylamine binding based on AF3 docking iPTM scores. Circle represents GPCRs that are differentially phosphorylated in response to tryptamine. Underlined human GPCRs represent known monoamine GPCRs. **(C)** Overview of *S. lacustris* tryptamine class C GPCR in relationship to human class C GPCRs including human metabotropic glutamate and GABA receptors (yellow). Star highlights the highest likely candidate for tryptamine binding based on AF3 docking iPTM scores. Circle represents GPCRs that are differentially phosphorylated in response to tryptamine. **(D)** Overview of *S. lacustris* tyramine class E GPCR in relationship to *Dictyostelium* class E cAMP GPCRs (blue). Star highlights the highest likely candidate for tyramine binding based on AF3 docking iPTM scores.

**Fig. S21.**
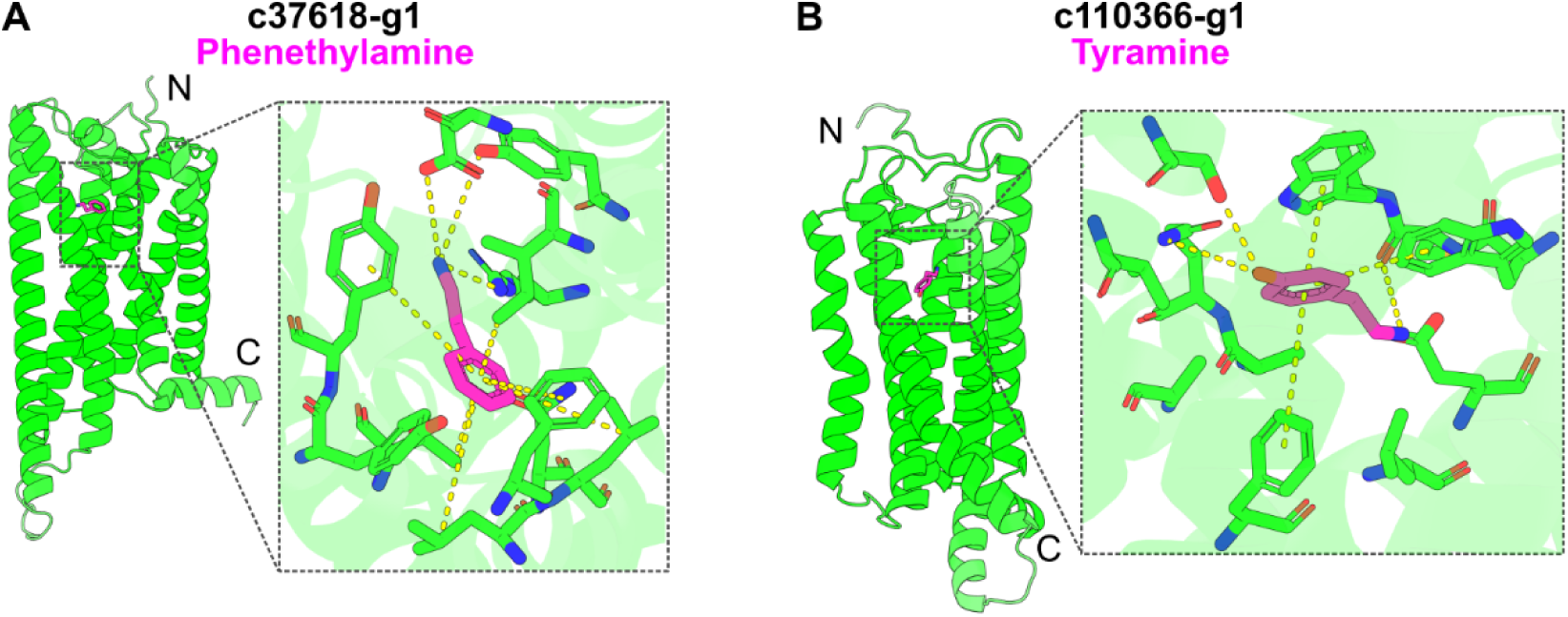
*S. lacustris* candidate phenylethylamine and tyramine GPCRs from AF3 docking analysis. **(A)** AF3 docking of phenethylamine to class A GPCR (c37618-g1, residues 20-335). Dashed line represent zoomed in highlights for the binding pockets of phenethylamine. **(B)** AF3 docking of tyramine to class E GPCR (c110366-g1, residues 16-315). Dashed line represent zoomed in highlights for the binding pockets of tyramine.

**Fig. S22.**
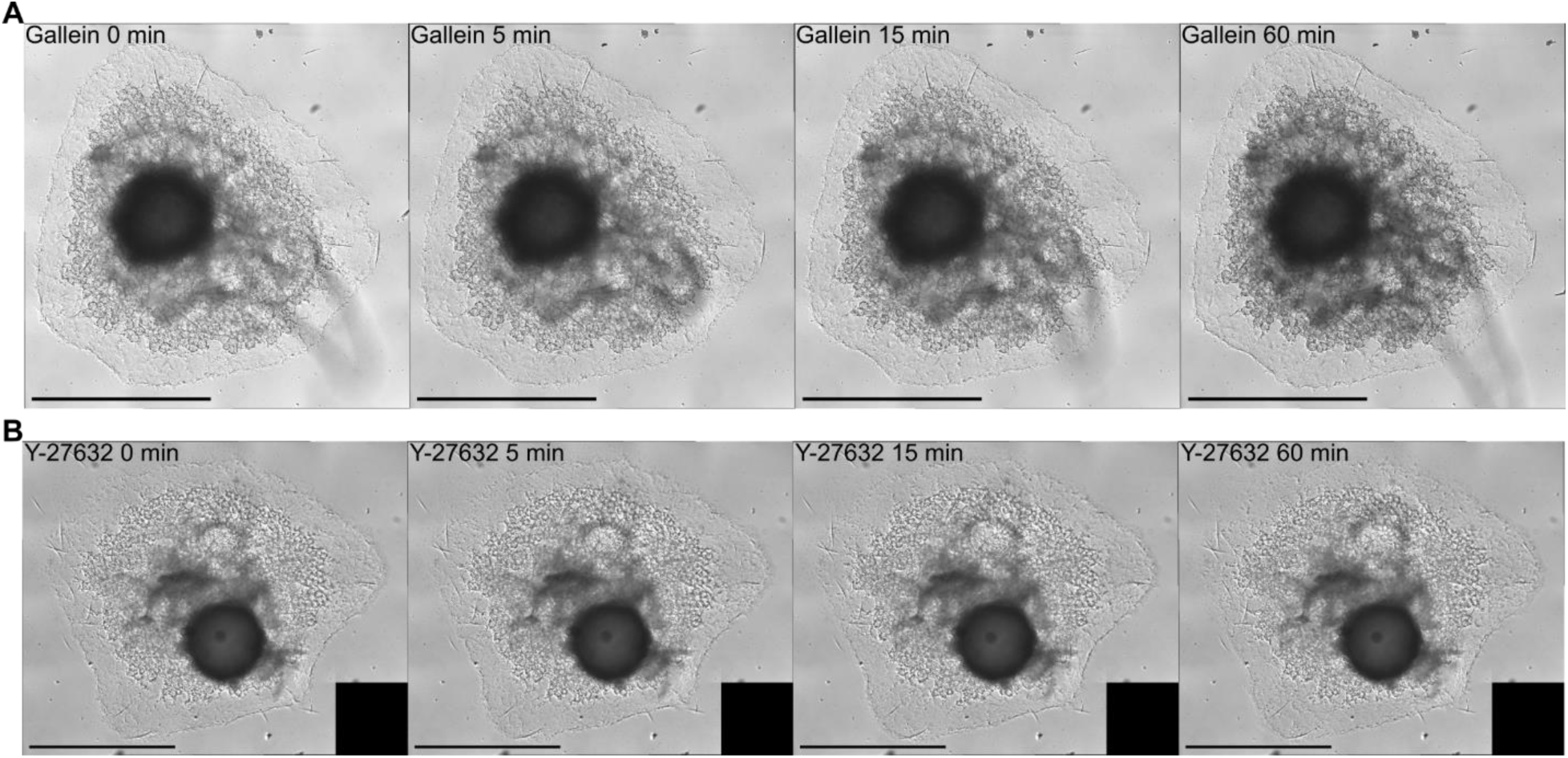
*S. lacustris* has minimal behavioral response to ROCK inhibitor (Y-27632) or Gβγ inhibitor (gallein) treatments alone. **(A)** DIC images of *S. lacustris* treated with 5 μM gallein across four timepoints. Scale bar, 1 mm. **(B)** DIC images of *S. lacustris* treated with 20 μM Y-27632 across four timepoints. Scale bar, 1 mm.

**Fig. S23.**
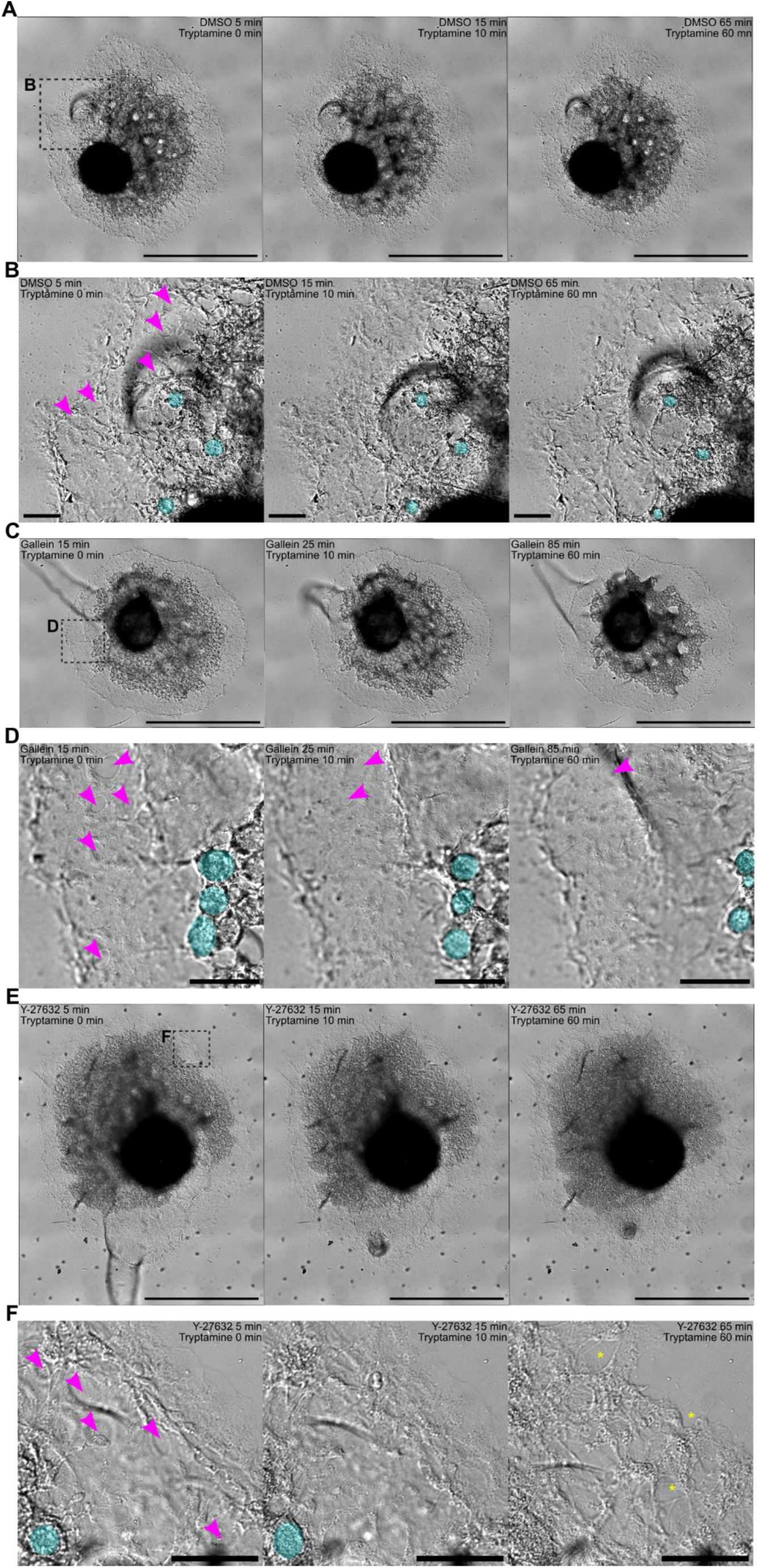
Gβγ and ROCK differentially regulate tryptamine-induced behavior in *S. lacustris*. **(A)** DIC images of control sponges treated with DMSO followed by tryptamine across three timepoints. Dashed box indicates the magnified inset in B. Scale bar, 1 mm. **(B)** Magnified inset of A. Magenta arrowhead points to the center of ostia formed by porocytes. Cyan circles indicate manual tracing of digestive choanocyte chambers. Scale bar, 100 μm. **(C)** DIC images of sponges treated with gallein followed by tryptamine across three timepoints. Dashed box indicates the magnified inset in D. Scale bar, 1 mm. **(D)** Magnified inset of C. Magenta arrowhead points to the center of ostia formed by porocytes. Cyan circles indicate manual tracing of digestive choanocyte chambers. Scale bar, 100 μm. **(E)** DIC images of sponges treated with Y-27632 followed by tryptamine across three timepoints. Dashed box indicates the magnified inset in F. Scale bar, 1 mm. **(F)** Magnified inset of E. Magenta arrowhead points to the center of ostia formed by porocytes. Cyan circles represent manual tracing of digestive choanocyte chambers. Yellow asterisks indicate loss of structural integrity at the base layer of the outer tent. Scale bar, 100 μm.

**Fig. S24.**
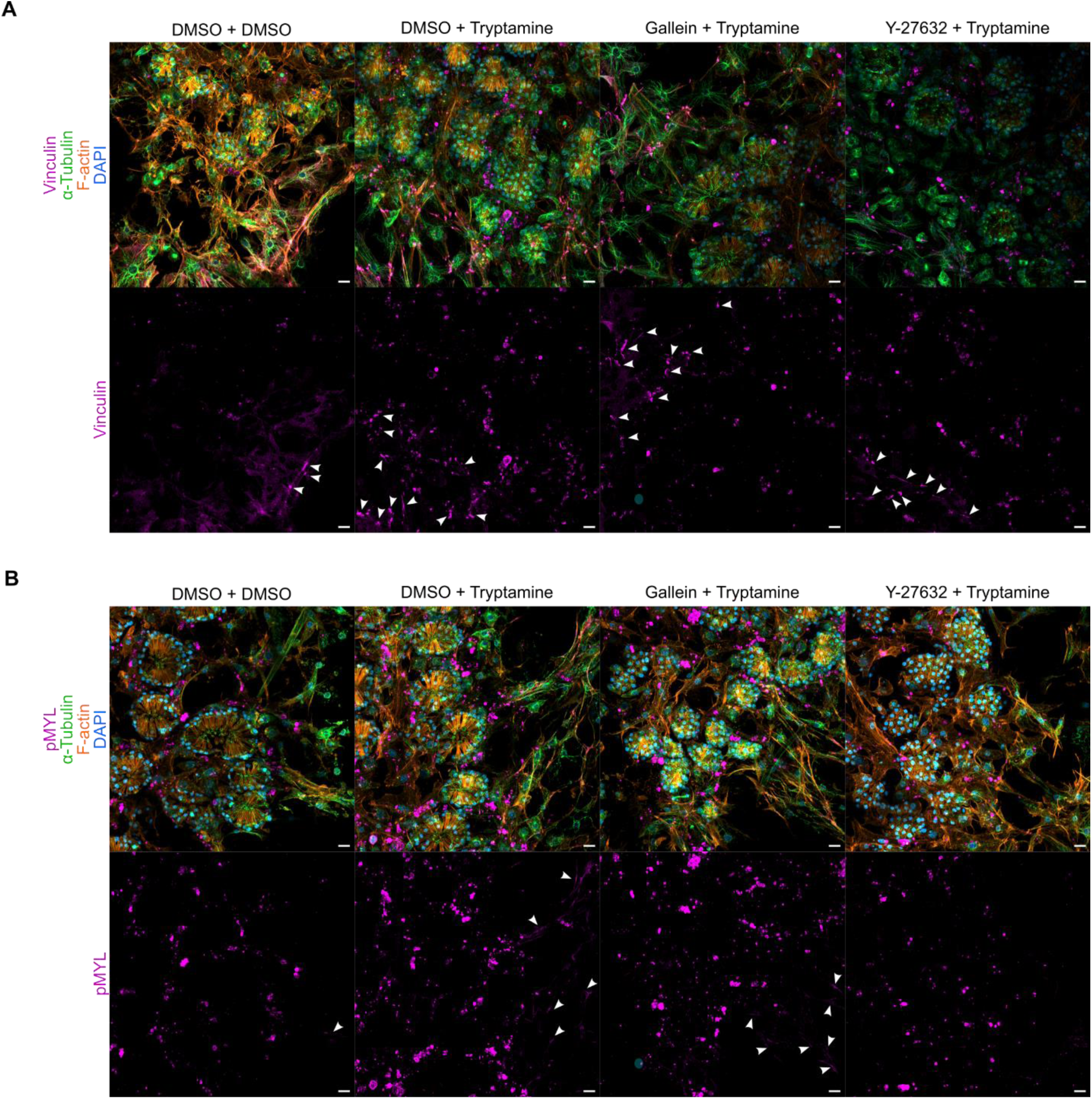
Gβγ and ROCK differentially regulate tryptamine signal transduction to modulate adhesion complexes and contractility. **(A)** Immunofluorescence maximum projection of *S. lacustris* choanoderm labeled with anti-*Em*Vinculin, anti-α-tubulin, phalloidin, and DAPI. White arrowhead indicates… Scale bar, 10 μm. **(B)** Immunofluorescence maximum projection of *S. lacustris* choanoderm labeled with anti-pMYL, anti-α-tubulin, phalloidin, and DAPI. White arrowhead indicates Scale bar, 10 μm.

**Fig. S25.**
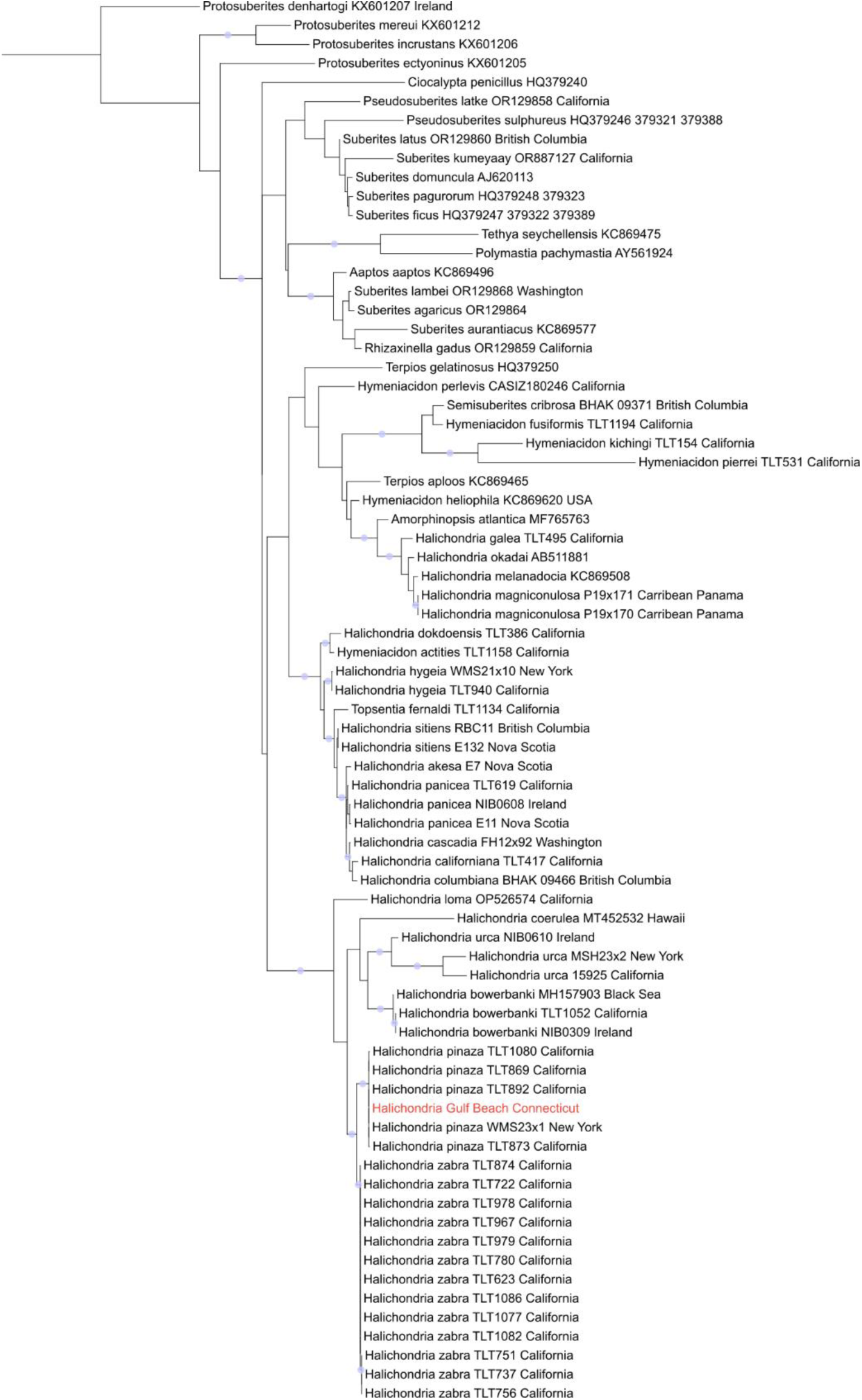
Maximum-likelihood phylogeny of *Halichondria* based on 28S rRNA. Maximum likelihood phylogeny for *Halichondria* based on 28S ribosomal RNA alignment. Blue circles at nodes indicate a bootstrap value ≥ 99%. *Protosuberites* and *Suberites* were used as outgroups. Red text indicates the positioning of *H. pinaza* collected in Milford, Connecticut.

**Fig. S26.**
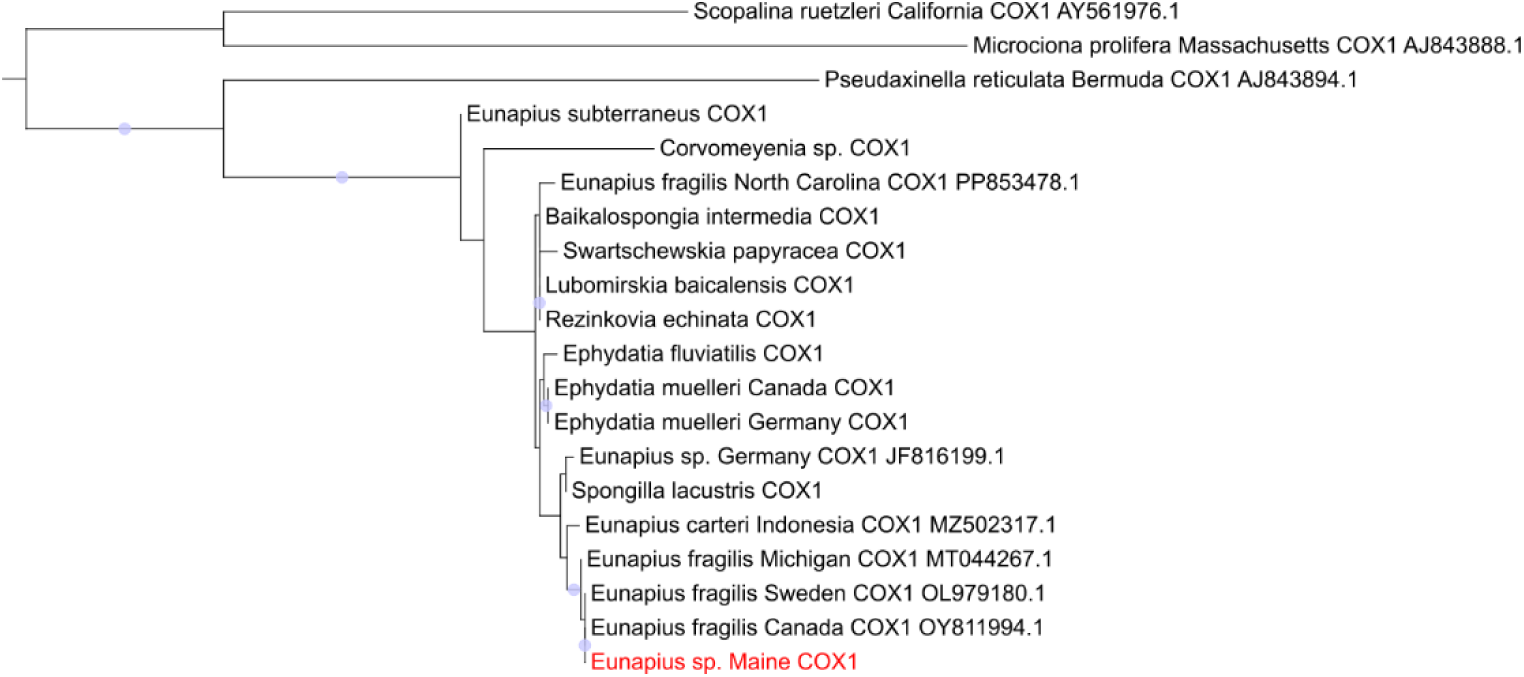
Maximum-likelihood phylogeny of freshwater sponges based on COI protein sequences. Maximum-likelihood phylogeny for freshwater sponges based on COI protein alignment. Blue circles at nodes indicate a bootstrap value ≥ 90%. *S. ruetzleri, M. prolifera,* and *P. reticulata* were marine sponges used as outgroup. Red text indicates the positioning of *E. fragilis* collected in Maine, USA.

