## Supplementary Materials and Methods for "An ancient monoaminergic signaling system coordinates contractility in a nerveless sponge"

**The PDF file includes:**

Materials and Methods
Figs. S1 to S26
Tables S1
References

**Other Supplementary Materials for this manuscript include the following:**

Movies S1 to S10
Data S1 to S5

### Materials and Methods

Collection and cultivation of sponges

During winter, freshwater sponges (*Spongilla lacustris*, *Ephydatia mulleri,* and *Eunapius fragilis* in this study) form protected packets of stem cells called gemmules, which can be collected and stored in the lab, and induced to hatch and form juvenile sponges. Adult specimens of *S. lacustris* in overwintering stage were collected on 12 January - 8 March 2022-2024 from Lake Constance, near Kressbronn, Germany (47.586N, 9.599E). Adult specimens of *E. mulleri* in overwintering stage were collected on 12 January - 8 March 2024 from Lake Constance, near Bodeman, Germany (47.805N, 9.031E). *E. fragilis* gemmules were a gift from April Horton from Maine, USA. Individual sponge patches, composed of the sponge skeleton and gemmules, were stored in lake water at 4°C until gemmule isolation. *Halichondria pinaza* adult specimens were collected on 19 October 2025 from Gulf Beach, Milford, CT, USA (41.210N, 73.047W) under the state of Connecticut Department of Energy & Environmental Protection permit #2427005. Adult specimens were sustained in marine aquaria with temperature-calibrated Gulf Beach water supplied with $O_2$ in excess until use. All sponge samples were collected compliant with local regulations.

Gemmules were extracted from the adult tissue via gentle agitation of the skeleton structure by lightly rubbing the sponge over sandpaper (3M, 220 Grit) using one finger covered in nitrile gloves. Gemmules from different sponge patches were washed several times with $ddH_2O$ and stored separately in M-medium (1 mM $CaCl_2 \cdot 6H_2O$, 0.5 mM $MgSO_4 \cdot 7H_2O$, 0.5 mM $NaHCO_3$, 0.05 mM KCl, 0.25 mM $Na_2SiO_3$).

Juvenile freshwater sponges (*S. lacustris*, *E. mulleri*, and *E. fragilis*) were grown by first screening for viable gemmules that look to be complete without damaged outer husk. Screened gemmules were placed in culture dishes with M-medium and kept at 18 °C in the dark. *H. pinaza* primmorphs were generated by reaggregation of single-cell dissociations (see below). To culture sponges for behavioral OCM, brightfield, DIC, or confocal imaging, 3-6 gemmules were placed in a 35 mm glass-bottom culture dish (MATTEK, P35G-1.5-14-C; Greiner Bio-One North America, 627861) containing 4 ml of M-medium. For the initial screen of behavioral response of *S. lacustris* to neurotransmitters, 3-6 gemmules were placed on glass-bottom 6-well, 12-well plates or 24-well plates (Cellvis, P6-1.5H-N, P12-1.5H-N, P24-1.5H-N), each well containing 4 mL, 1.5 ml, or 0.75 mL of M-medium (to ensure a water height of ~4 mm remained constant). To culture *S. lacustris* for heavy-isotope tracing metabolomics, 50 gemmules were placed in a 35 mm tissue-culture dish containing 4 mL of M-medium. To culture *S. lacustris* for phosphoproteomics, 100 gemmules were placed in a 60 mm tissue-culture dish containing 10 mL of filtered lake water or M-medium. To culture *S. lacustris* for transcriptomics, 50 gemmules were placed in a 60 mm tissue-culture dish containing 10 mL of filtered lake water or M-medium.

M-medium was replaced every other day, beginning around day 7 when juvenile sponges began to develop canals. Juvenile sponges were treated or fixed for experiments after 8-14 days of growth, when they had acquired all major features of adults: an osculum, a well-developed canal system, and numerous choanocyte chambers. Variance in the exact time at which this stage occurs is high, even for neighboring sponges grown from clonal gemmules that are present in the same culture dish. In general, we tended to err on the side of letting sponges develop further, so that all sponges in the dish exhibited these features.

Generation of *H. pinaza* primmorphs

Pieces of *H. pinaza* adult tissue were dissected from rock using a clean razor blade and transferred to $Ca^{2+}$, $Mg^{2+}$ free artificial sea water (495 mM NaCl, 9.7 mM KCl, 27.6 mM $NaHCO_3$, 50 mM Tris-HCl) with 25 mM EDTA for 1 hour rocking at 100 RPM under room temperature. Sponges were gently agitated again by repeated pipetting, causing a relatively rapid dissociation of all cells. Once completely dissociated, cells were passed through a 40 µm mesh filter (Fisher, 22363547) twice and collected in polystyrene test tubes (Falcon, 352058). Cells were washed twice in 0.2 µm filtered (Corning, 430758) Gulf Beach water (fGBW) with 5 min centrifugation at 800 G in-between. Resuspended cells (in 100 uL of fGBW) were diluted 1:450 (9 uL) into 35 mm glass bottom dishes filled with 4 mL of fGBW and left on rotator at 20 RPM under room temperature for 72 hours in the dark before moving for long-term culture at 18 °C.

DNA and RNA isolation

Deoxyribonucleic acid (DNA) and RNA were extracted using the Qiagen All-Prep Micro Kit (Qiagen, 80284) according to manufacturer's instructions, with the following minor modification. Each plate with sponges was washed once with 1X phosphate-buffered saline (PBS; Life Technologies, 10010-023) and lysed with 350 µL RLT lysis buffer with repetitive pipetting. Lysed sponges were placed into Eppendorf tubes and homogenized five times with a 16G needle (BD Biosciences, 305198) before proceeding with loading onto DNA and RNA-binding columns. Extracted DNA and RNA were stored at -80 °C and submitted to Yale Center for Genome Analysis for Illumina-based whole-genome sequencing or whole-transcriptome sequencing.

Sponge DNA and RNA sequencing

DNA samples were purified using Ampure XP spri beads (Beckman Coulter, A63882) and library preparation was performed using the Watchmakers Library Prep Kit with Fragmentation (Watchmaker Genomics, 7K0013-096). RNA polyA library preparation was performed using the Watchmaker mRNA Library Prep Kit (7VK001-096). Both DNA and RNA samples were sequenced at the Yale Center for Genome Analysis on a Novaseq X Plus at 2×150bp to obtain 20 million reads per sample.

Sponge species identification and *de novo* transcriptome assembly

*S. lacustris* and *E. mulleri* gemmules from individually cleaned patches were barcoded by Sanger sequencing of PCR products generated by primers targeting the COI gene (FWD: GGTCAACAAATCATAAAGAYATYGG; RVS: TGTTGRGGGAAAAARGTTAAATT). Species identity of *H. pinaza* primmorphs were identified based on 28S ribosomal RNA alignment generated using *in silico* PCR (*65*) on whole genome sequencing reads using published primers (*66*). Gulf beach 28S ribosomal RNA sequence matched 100% to *H. pinaza* found in California [TLT892, TLT873] and New York [WMS23x1] (*66*). As *H. pinaza* has never been described in Connecticut, we aligned its 28S sequence to the sequences of recently described *Halichondria* species around the Pacific and Atlantic using Clustal Omega with 100 cluster size and full iterations. Phylogenies were inferred using maximum likelihood in IQ-TREE2 (*67*). The best-fit substitution model was selected by ModelFinder (*68*). Branch support was estimated using 1000 ultrafast bootstrap replicates with the BNNI correction (fig. S25) (*69*).

*E. fragilis* was identified based on a COI phylogeny containing sequences from diverse freshwater sponges. First, we generated a *de novo* mitochondrial genome assembly using GetOrganelle (*70*) on whole genome DNA sequencing data, using the animal_mt mode to target animal mitochondrial signatures, supplemented with a custom seed database (freshwater-sponges_mt.fasta) containing representative Spongillidae mitochondrial sequences to enhance read recruitment. The process utilized the "bait-and-extend" algorithm with default K-mer values. The mitochondrial genome was aligned with the *Eunapius subterraneus* COI gene (NC_016431) and the corresponding region was extracted for BLAST. The top BLAST hit revealed 100% identity to mitochondrial genome of *E. fragilis* from Canada sequenced as part of the Wellcome Sanger Tree of Life Programme (*71*). For sequence alignment, we obtained additional COI nucleotide sequences available on NCBI labeled with the *Eunapius* genus (Fig S. Eunapius). As all COI sequences were partial, the first 900 bp of COX1 were extracted from mitochondrial genomes from freshwater sponges (including the *de novo* mitochondrial genome of Maine *E. fragilis sp.*) for alignment using Clustal Omega with 100 cluster size and full iterations. Phylogenies were inferred using maximum likelihood in IQ-TREE2 (*67*). The best-fit substitution model was selected by ModelFinder (*68*). Branch support was estimated using 1000 ultrafast bootstrap replicates with the BNNI correction (fig. S26) (*69*).

Optical coherence microscopy

A detailed description of optical coherence microscopy (OCM) is described previously (*9*). For live imaging, sponge grown in 35 mm glass-bottom culture dishes and positioned on an X, Y, Z manual translation sample stage. To record tryptamine (Sigma, 193747), phenethylamine (Sigma, 128945), and narciclasine (MedChemExpress LLC, HY-16563) treatment, sponge specimen began with 4 mL of M-medium and 2 mL of monoamine or DMSO (Sigma, 472301) control solution (3X concentration, e.g., 112.5 µM and 0.1% for *S. lacustris*) was carefully added to the dish without direct disruption of sponge tissue to a final concentration of 1X (e.g., 37.5 µM and 0.03% for *S. lacustris*). In case of double treatments, sponge began with 2.67 mL of M-medium and 1.33 mL of DMSO, gallein (MedChemExpress LLC, HY-D0254), or Y-27632 (MedChemExpress LLC, HY-10583) solution (3X concentration) was carefully added to the dish to a final concentration of 0.03%, 5 µM, or 20 µM before the addition of tryptamine. We empirically determined that a 5 min and 15 min gap was required for tryptamine addition after Y-27632 and gallein treatment, respectively, for the sponge tissue to reach a stabilized state before tryptamine addition (fig. S22). OCM image acquisition (1 volume / 20 sec) started before the addition of pharmaceuticals or monoamines to ensure that no endogenous deflation was occurring and stopped 60 min after monoamine addition. For *S. lacustris* washout experiments, OCM image acquisition was set to 1 volume / 2 min and imaged for 120 min after monoamine addition and 60 min after washout. All OCM raw data were processed as previously described (*9*). We empirically determined that a 5 min and 15 min gap was required after Y-27632 and gallein treatment, respectively, for the sponge tissue to stabilize before tryptamine addition (fig. s22).

Brightfield/Differential interference contrast microscopy

For live brightfield and DIC imaging, specimens grown in 35 mm glass-bottom culture dishes were imaged on a Leica THUNDER Imager using a 4X dry (NA=0.13), 10X dry (NA=0.45), or 20X dry (NA=0.8) objective.

The initial behavioral screen took place in glass-bottom plates. Sponges in 6-, 12-, or 24-well plates begin with 4 mL, 1.5 ml, or 0.75 mL M-medium. Different volumes of treatment solutions (2 mL, 0.75 mL, or 0.375 mL) at 3X concentration were carefully added to each well to a final concentration of 1X (Fig. S1). Image acquisition (~1 image/ 20 sec) started before the addition of candidate transmitter to ensure that no endogenous behavior was occurring and was stopped 60 min after treatment. The following amino acids, ammonium, monoamine, and polyamine transmitters were tested: L-glutamic acid monosodium (Thermo, J63424.09), L-γ-aminobutyric acid (GABA, Sigma, A2129), glycine (Sigma, G7126), acetylcholine chloride (Sigma, A6625), (−) epinephrine (dissolved in 1% acetic acid in M-medium, Sigma, E4250), (−) norepinephrine bitartrate (Sigma, A9512), dopamine hydrochloride (Sigma, E4250), serotonin hydrochloride (Sigma, H9523), histamine dihydrochloride (Sigma, H7250), melatonin (Sigma, M5250), octopamine hydrochloride (Sigma, O0250), tryptamine, tyramine (Sigma, 193747), phenethylamine, spermine (Sigma, S3256), spermidine (Sigma, S2626), cadaverine dihydrochloride (Sigma, C8561), and putrescine dihydrochloride (Sigma, P7505). As different transmitters come in different salt formats or required specialized solvent, the following control substances were used to account for the non-specific effect of ions or solvent: ammonium chloride (Thomas Scientific, C842G29), glacial acetic acid (A.C.S. Reagent, 9509-33), sodium tartrate dihydrate (Fisher Scientific Company, S-433), hydrochloric acid (ACS Reagent, 9535-33), and DMSO. Data were collected from at least two independent biological replicates ($n \geq 2$).

To record additional tryptamine and phenethylamine treatments, sponge began with 4 mL of M-medium and 2 mL of monoamine solution (3X concentration) was carefully added to the dish to a final concentration of 1X. In case of double treatment, the sponge specimen began with 2.67 mL of M-medium and 1.33 mL of DMSO, gallein, or Y-27632 solution (3X concentration) was carefully added to the dish to a final concentration of 0.03%, 5 µM, or 20 µM, respectively, before the addition of tryptamine. Image acquisition (~1 image/ 20 sec) started before the addition of pharmaceuticals or monoamines to ensure that no endogenous behavior was occurring and was stopped 60 min after monoamine addition. Data were collected from three independent biological replicates (n=3).

For live brightfield imaging of tyramine treatment, sponges were imaged at 1 image/min (*S.* lacustris) or 1 image/2 min (*E. mulleri* and *E. fragilis*) intervals for 12 hours and 10 minutes in total while covered by a lid, except for the addition of tyramine. For *S. lacustris*, data were collected from seven independent biological replicates (n=7). For *E. mulleri* and *E. fragilis* data were collected from three independent biological replicates (n=3).

<u>Behavioral microscopy data processing and quantification</u>

For the quantification of cross-sectional tissue area and excurrent canal area of *S. lacustris* treated with tryptamine, representative OCM slices across timelapse was processed in FIJI to adjust brightness, contrast, crop, and Gaussian blur ($\sigma = 1$) the region containing the entire sponge. Image sequences were then transferred to Ilastik v1.4.1rc2-gpu. Three labels were assigned to distinguish the sponge's canal system from the surrounding tissue and background: (1) background, (2) excurrent canals, and (3) tissue. ~40 images from each sequence were manually selected for training until accurate segmentations were achieved for the entire sequence. Subsequently, the segmentation labels were imported into Python and the cross-sectional areas of tissue and excurrent canal system were calculated and plotted. For plots, the time of tryptamine addition was set to 0 min. Areas were normalized against the first frame in each sequence (showing sponges immediately prior to treatment) to plot relative changes.

For the quantification of tent and tissue height of *S. lacustris* treated with phenethylamine, representative OCM slices across the timelapse were processed in FIJI to adjust brightness, contrast, crop, and Gaussian blur (σ = 1) the region containing the entire sponge. Each timelapse was subset into 42 evenly spaced timepoints and the tent/tissue heights were measured manually using the line tool. Efforts were made to ensure the same tissue area and position of height measurements were consistent across samples. Measured heights were imported into Python for plotting. For plots, the time of tryptamine addition was set to 0 min. Heights were normalized against the first frame in each sequence (sponges right before treatment) to plot relative changes.

For the quantification of top-down tissue area of *S. lacustris* treated with tyramine, representative brightfield slices across timelapse was processed in FIJI to adjust brightness, contrast, and crop the images around sponges. Three labels were assigned to distinguish the sponge's canal system from the surrounding tissue and background: (1) background, (2) inner tissue area (including the choanoderm and canals), and (3) outer tissue area (including the outer tent and basopinacoderm). ~40 images from each sequence were manually selected for training until accurate segmentations were achieved for the entire sequence. Subsequently, segmentation labels were imported into Python and the areas of tissue and excurrent canal system segmentations were calculated and plotted. For plots, the time of tyramine addition was set to 0 min. Areas were normalized against the first frame in each sequence (sponges right before treatment) to plot relative changes.

For the quantification of canal movements of *S. lacustris* treated with tyramine, bright field timelapses of sponges were processed as described above. A line was drawn with the segmented line tool circumferentially around the tissue connecting incurrent canals while crossing orthogonally to as many excurrent canal structures as possible (Fig. 1K). Re-slicing of the line generated a kymograph representing the movement of canals and tissue over time. Kymographs were imported into R and the correlation distance between pair-wise pixels (i.e. two vertically adjacent pixels) on the kymograph were calculated after the addition of tyramine. Plots represent the correlation distance across a time with a window of 1 frame (1 min). For visualization purposes, the limits of y-axis (correlation distance between pair-wise pixels) were set to remove extrema during a deflation event. One DMSO-treated biological replicates were not considered for this analysis as it has not reached the juvenile stage with cells observed to leave the gemmule.

For the quantification of tent height of *S. lacustris* treated with gallein + tryptamine or Y-27632 + tryptamine representative OCM slices across timelapse was processed in FIJI to adjust brightness, contrast, crop, and Gaussian blur (σ = 1) the images around sponges. Each timelapse was subset into 51 or 42 evenly spaced timepoints and the tent/tissue heights were measured manually using the line tool. Efforts were made to ensure the same tissue area and position of height measurements were consistent. Measured heights were imported into Python for plotting. For plots, the time of gallein addition was set to −15 min, the time of Y-27634 addition was set to −10 min, and the time of tryptamine addition was set to 0 min. Heights were normalized against the first frame in each sequence (sponges right before tryptamine treatment) to plot relative changes.

For the quantification of cross-sectional tissue area and excurrent canal area of *S. lacustris* treated with gallein + tryptamine or Y-27632 + tryptamine, representative OCM slices across timelapse were processed in FIJI to adjust brightness, contrast, crop, and Gaussian blur ($\sigma = 1$) the images around sponges. Each timelapse was subset into 51 or 42 evenly spaced timepoints. Image sequences were then transferred to Ilastik. Three labels were assigned to distinguish the sponge's canal system from the surrounding tissue and background: (1) background, (2) canals, and (3) tissue. ~20 images from each sequence were manually selected for training until accurate segmentations were achieved for the entire sequence. Subsequently, the simple segmentation labels were imported into Python and the cross-sectional areas of canal system segmentations were calculated and plotted. For plots, the time of gallein addition was set to −15 min, the time of Y-27634 addition was set to −10 min, and the time of tryptamine addition was set to 0 min. Areas were normalized against the first frame in each sequence (sponges right before tryptamine treatment) to plot relative changes.

<u>Examination of bacterial load in *S. lacustris* gemmules</u>

Approximately 50 *S. lacustris* gemmules were cleaned with 1% $H_2O_2$ by incubation for 5 min at 4°C with 5 times of rinsing in M-media. Surface sterilized gemmules and untreated gemmules were resuspended in 200 µL sterile M-medium and crushed with a sterile Eppendorf pestle. Gemmule extracts were vortexed briefly and 100 µL was plated on LB agar plates (5 g/L yeast extract, 10 g/L peptone, 10 g/L NaCl, 12 g/L agar) without antibiotics. Plates were stored at 18 °C for two days before imaging with a ChemiDoc Imaging Systems (Biorad, California, USA)

<u>Untargeted heavy-isotope tracing metabolomics sample preparation and LC-MS/MS measurement</u>

*Sample preparation:* 15 x 50 *S. lacustris* gemmules were cleaned with 1% $H_2O_2$ by incubation for 5 min at 4°C with 5 times of rinsing in M-media. Surface sterilized gemmules were plated in 4 mL filtered lake water (in 35 mm culture dishes) and kept at 18°C in the dark until usage. At day 9 post plating, bright field pictures were taken of the sponges to record their approximate size, then the M-medium was removed and replaced with 4 mL of 1 mM precursor amino acids dissolved in M-medium: L-tryptophan (RPI, T60080, n=2), L-phenylalanine (Sigma, P2126; n=2), L-tyrosine (Sigma, T3754, n=2), L-$^{13}C_{11}$-tryptophan (MedChemExpress LLC, HY-N0623S2; n=3), L-$^{13}C_9$-phenylalanine (MedChemExpress LLC, HY-N0215S10; n=3), and L-$^{13}C_9$-tyrosine (MedChemExpress LLC, HY-N0473S3; n=3). Heavy isotope-labeled amino acids were ~90% pure. Twenty-four hours after incubation in amino acids, the medium was removed, and the sponges were immediately flash frozen by dipping the bottom of the culture dishes into liquid nitrogen. To prevent thawing and potential metabolic activity, the dishes were put on metal blocks cooled to approximately −80 °C by dry ice. A pre-chilled 16 G needle (BD Biosciences, 305198) was used to scrape off the sponges and transfer into pre-cooled 2 mL cryotubes. The samples were stored at −80 °C until further processing.

For metabolite extraction, 300 µL of 80% acetonitrile (ACN) was added to each sample. Subsequently, samples were homogenized on dry ice via a bead beater (FastPrep-24; MP Biomedicals, CA, USA) at 6.0 m/s (5 x 30 s, 5 min pause time) using 1.0 mm and 2.3 mm zirconia/glass beads (Biospec Products, OK, USA). After incubation for 20 min at -20°C, samples were centrifuged at 15,000 × g and 4 °C for 10 minutes using a 5415R microcentrifuge (Eppendorf, Hamburg, Germany). Ultimately, the supernatants were transferred to analytical glass vials for LC-MS analysis, which was initiated within one hour after sample preparation.

LC-MS/MS analysis was performed on a Vanquish UHPLC system coupled to an Orbitrap Exploris 240 high-resolution mass spectrometer (Thermo Fisher Scientific, MA, USA) in positive ESI (electrospray ionization) mode. Chromatographic separation was carried out on an Atlantis Premier BEH Z-HILIC column (Waters, MA, USA; 2.1 mm x 100 mm, 1.7 µm) at a flow rate of 0.25 mL/min. The mobile phase consisted of water:acetonitrile (9:1, v/v; mobile phase phase A) and acetonitrile:water (9:1, v/v; mobile phase B), which were modified with a total buffer concentration of 10 mM ammonium formate. The aqueous portion of each mobile phase was pH-adjusted (negative mode: pH 9.0 via addition of ammonium hydroxide; positive mode: pH 3.0 via addition of formic acid). The following gradient (20 min total run time including re-equilibration) was applied (time [min]/%B): 0/95, 2/95, 14.5/60, 16/60, 16.5/95, 20/95. Column temperature was maintained at 40°C, the autosampler was set to 4°C and sample injection volume was 5 µL. Analytes were recorded via a full scan with a mass resolving power of 120,000 over a mass range from 60 - 900 *m/z* (scan time: 100 ms, RF lens: 70%). To obtain MS/MS fragment spectra, data-dependent acquisition was carried out (resolving power: 15,000; scan time: 22 ms; stepped collision energies [%]: 30/50/70; cycle time: 900 ms). Ion source parameters were set to the following values: spray voltage: 4100 V, sheath gas: 30 psi, auxiliary gas: 5 psi, sweep gas: 0 psi, ion transfer tube temperature: 350°C, vaporizer temperature: 300°C.

All experimental samples were measured in a randomized manner, with cell extracts and cell supernatant extracts being analyzed in individual sequences. Pooled quality control (QC) samples were prepared by mixing equal aliquots from each processed sample. Multiple QC samples were injected at the beginning of the analysis for system equilibration, throughout the sequence after every other experimental sample, and at the end to monitor instrument performance over time. For determination of background signals and subsequent background subtraction, an additional processed blank sample was recorded. Data was processed using MS-DIAL 4.9 (*72*) and raw peak intensity data was exported for relative metabolite quantification. Level 1 feature identification was based on an in-house library for metabolomics (EMBL-MCF 2.0) using accurate mass, isotope pattern, MS/MS fragmentation, and retention time information with a minimum matching score of 80%. For untargeted isotope tracing analysis, CompoundDiscoverer 3.2 (Thermo Fisher Scientific) was used and relative peak areas were exported using the SIL_Area Version 1.1 scripting node.

Identification of candidate decarboxylases, vesicular transporters, and GPCRs in *S. lacustris*.

For decarboxylases and vesicular transporters, human AADC (Uniprot ID: P20711), SLC6A17 (Q9H1V8), SLC17A5 (Q9NRA2), SLC17A7 (Q9P2U7), SLC18A1 (P54219), SLC18B1 (Q6NT16), and SLC22A1 (O15245) canonical amino acid sequences were used as query for BLAST search against the *S. lacustris de novo* 70 amino acid proteome.

For GPCRs, representatives of all six GPCR families were used as query for BLAST search against the *S. lacustrsi de novo* 70 amino acid proteome. These sequences included human CCR1 (P32246) ACKR1 (Q16570), AGTR1 (P30556), OPRD (P41143), LT4R1 (Q15722), V1AR (P37288), MTLR (O43193), FPR1 (P21462), MTR1A (P48039), LSHR (P22888), HCAR1 (Q9BXC0), PTAFR (P25105), S1PR1 (P21453), PD2R (Q13258), GP132 (Q9UNW8), OPSB (P03999), ADRB1 (P08588), ACM1 (P11229), 5HT1A (P08908), VN1R1 (Q9GZP7), GLR (P47871), CELR1 (Q9NYQ6), DIHR (P35464), GRM5 (P41594), FZD1 (Q9UP38), yeast STE2 (D6VTK4), and *Dictyostelium* CAR1 (P13773).

All protein hits were deemed as a candidate after manual examination to ensure that all the necessary domains are present. These included the pyridoxal-dependent decarboxylase domain, 12 transmembrane domains typical of transporters in the major facilitator superfamily, and 7 transmembrane domains typical of GPCRs. For hits with incomplete sequences in our *de novo* proteome, we used TBLASTN to search incomplete *S. lacustris* protein sequence against transcript reads from *S. lacustris* PACBIO Isoseq. Up to 20 top-hits from TBLASTN were aligned using MUSCLE to generate a consensus transcript for translation. Complete *S. lacustris* candidates were subsequently filtered based on their single-cell expression. Candidate proteins with no or minimal expression in any cell type were ignored for future analysis (Fig. S9 and Fig, S13).

<u>*C. elegans* plasmids</u>

Decarboxylase and vesicular transporter rescue plasmids were generated by codon optimizing *S. lacustris* candidate protein sequences for *C. elegans*. Custom DNA fragments were generated using GenScript's GenTitan™ platform or Twist Biosciences Gene Fragments platform. Each CDS was combined with the endogenous *C. elegans cat-1* promoter and 5'UTR (*cat-1p*, -1624 bp to -1 bp relative to ATG for N2 strain) (*73*) and a SL2::mCherry::unc-54(3'UTR) bicistronic cassette backbone (*74*) using overlap PCR and Gibson assembly. *C. elegans* AADC (*bas-1*), *tdc-1*, and VMAT (*cat-1*) CDS were also cloned in a similar manner to serve as positive controls.

<u>*C. elegans* strains and maintenance.</u>

All worms were maintained on Nematode Growth Media (NGM) seeded with *Escherichia coli* OP50 at 20 °C, and hermaphrodites were used for all experiments. Transgenic worms carrying extrachromosomal arrays were generated by injections as previously described (*75*). All injection mixes included the following elements: 5 ng/µL of rescue plasmid carrying a decarboxylase or vesicular transporter driven by *cat-1p*, 2 ng/µL of *myo2p::GFP* as a co-injection marker in the pharynx, and 15 ng/µL *rps-0p::HygR* as a selection marker for hygromycin resistance. A list of *C. elegans* strains used in this work is provided in Table S1. When two independent transgenic lines were able to be generated, both lines were used for all experiments. For all behavioral assays, animals were maintained on these plates for a single generation by transferring 6-10 adult worms or 40-50 eggs or 30-40 L4 worms at 20°C.

<u>*C elegans* behavioral assays.</u>

*Tyramine-dependent touch reversal (TR):* TR was performed as previously described (*24*). ~15 age-matched L4 animals were transferred to OP50-seeded NGM plates and grown at

20 °C one day before assay until maturation into young adults. Young adult worms were touched with a fine eyelash behind the posterior bulb of the pharynx and number of reversal body bends were recorded across a minimum of three independent experiments and the total biological replicates are below: N2, n=22; RB681, n=25; MOY0155, n=27; MOY0156, n=27; MOY0157, n=28; MOY0158, n=28; MOY0159, n=25; MOY0161, n=24; MOY0162, n=23; MOY0163, n=20; MOY0164, n=17; MOY0165, n=28; MOY0244, n=25; MOY0245, n=23; MOY0246, n=23; MOY0247, n=22; MOY0248, n=25; MOY0249, n=23; RB993, n=25; MOY0171, n=26; MOY0172, n=25; MOY0173, n=24; MOY0174, n=25). In all cases, plates were coded so that the experimenter was blind to the genotype of the assayed strains.

*Dopamine-dependent basal slow rate (BSR)*: BSR was performed as previously described (*22*) with the following modifications. ~30 age-matched L4 animals were transferred to OP50-seeded NGM plates and grown at 20 °C a day before assay for maturation into young adults. ~20 well-fed young adult animals were first transferred onto NGM plate with a thin lawn of OP50. Five minutes after transfer, the number of body bends in 20 s intervals was sequentially recorded for all the animals on the assay plate to calculate locomotory rate on food. Then worms were carefully picked with minimal number of bacteria onto empty NGM plates. Five minutes after transfer, the number of body bends in 20 s intervals was sequentially recorded for all the animals on the assay plate to calculate locomotory rate off food. The basal slow rate is calculated as $\left(1-\frac{\text{on−food locomotory rate}}{\text{off−food locomotory rate.)}}\right)\times 100\%$. A minimum of three independent experiments were performed and the total biological replicates are below: N2, n=23; RB681, n=22; MOY0155, n=25; MOY0156, n=27; MOY0165, n=23; MOY0248, n=22; MOY0249, n=30; LC33, n=22; MOY0166, n=23; MOY0167, n=24; MOY0168, n=16; MOY0169, n=23; MOY0170, n=18. In all cases, plates were coded so that the experimenter was blind to the genotype of the assayed strains.

*C. elegans* serotonin immunofluorescence

Serotonin immunofluorescence was done as previously described (*76*). Dense well-fed worms were fixed, denatured, digested, and stained with 1:100 anti-serotonin antibody (Sigma, S5545) and 1:100 anti-rabbit Alexa Fluo 647 antibody (Jackson ImmunoResearch, 111-605-003). Stained worms were mounted on 1% agarose (americanBIO, AB00972) pad, covered with #1.5 glass coverslip (VWR, 48366-249), and imaged with a 63X water immersion objective (NA=1.2) on a Leica THUNDER Imager Cell with DIC, 476 nm excitation LED with 519 nm emission filter, and 605 nm excitation LED with 642 nm emission filter. For wildtype (N2) and *bas-1 (tm351)* worms, a Z-stack image was taken for every worm. For *cat-1 (ok411)* worms, a Z-stack image was taken for worms that showed positive NSMs neuron staining for serotonin. For extrachromosomal array strains, a Z-stack image was only taken if a worm possesses the co-injection pharynx marker (*myo2p::GFP*). Three independent technical replicates were performed and the total biological replicates consisting of adult/late L4 worms are below: N2, n=90; RB681, n=90; MOY0155, n=90; MOY0156, n=90; MOY0165, n=88; MOY0248, n=90; MOY0249, n =90; LC33, n=90; MOY0166, n=90; MOY0167, n=90; MOY0168, n=90; MOY0169, n=90.

*S. lacustris* hybridization chain reaction (HCR)

HCR probes for RNA transcripts of interest were designed by Molecular Instruments after providing *S. lacustris* transcripts for genes or cell types of interest. Probes were manually screened against *S. lacustris de novo* transcriptome by BLAST to ensure specificity.

Juvenile *S. lacustris* grown on 35 mm dishes were fixed day 9-12 post plating. Sponges were fixed overnight with 4% PFA (Thermo, 28908) in ¼ HF buffer (590 mM NaCl; 6.7 mM KCl; 7.6 mM $CaCl_2$; 2.4 mM NaHC) at 4 °C. After washing out fixative with ¼ HF buffer, dehydration was performed by a series of washes in 30% methanol (JT Baker, JTB-9093-03; in ¼ HF buffer), 70% methanol, 100% methanol, 100% ethanol (Sigma, E7023), 70% ethanol, and 30% ethanol in 1X PBS with 0.1% tween (PBST). Sponges were treated with 5 µg/mL Proteinase K (Sigma, 03115828001) in PBST for 1.5 minutes at room temperature and quenched twice with 2 mg/mL glycine for 3 minutes. Post-fixation was performed by incubation of sponges in 4% PFA in PBST for 1 hour at room temperature. After two PBST washes and two 1X saline sodium citrates (SSC; Sigma, S6639) washes for 5 minutes each, sponges were incubated in HCR™ HiFi Probe Hybridization Buffer (HF buffer; Molecular Instruments) for 30 minutes at 37 °C. While keeping the sponge at 37 °C, carefully remove HF buffer and add hybridization probe (Table S HCR; dissolved 1:100 in HF buffer) for overnight 37 °C incubation. The next day, prepare amplifier hairpins according to the manufacturer's guidelines. Sponges from overnight probe incubation were washed four times in preheated HCR™ Probe Wash Buffer (PWB; Molecular Instruments) at 37 °C, two washes of 1X SSC at room temperature, and finally two washes in HCR™ Amplifier Buffer (AB; Molecular Instrument) for 3 minutes each. Amplifier hairpins were mixed in AB and added to the specimens and incubated overnight at RT in the dark. After amplification, sponges were washed twice with 1X SSC at room temperature for 3 minutes each. Sponges were stained with 1 µg/mL DAPI (Invitrogen, D3571) in 1X PBS for 20 minutes at room temperature and washed once with 1X PBS before mounting in Fluoromount-G (SouthernBiotech, 0100-01) for confocal microscopy on a Zeiss LSM 980.

Samples were imaged with a 20X air objective (NA=0.8) or 40 X water immersion objective (NA=1.2) as Z-stacks in multiplex CO-8Y Aryscan 2 mode and later underwent Aryscan processing with default parameters.

<u>Phylogenetic analysis of *S. lacustris* decarboxylases, vesicular transporters, and GPCRs</u>

*S. lacustris* amino acid sequences were queried via BLAST (*77*) and PROST (*78*) against a custom curated proteome consisting of unicellular eukaryotes, fungi, sponges, trichoplax, cnidarians, and model bilaterians (*Aurelia aurita*, *Caenorhabditis elegans*, *Drosophila melanogaster*, *Homo sapiens*, *Oscarella lobularis*, *Schmidtea mediterranea*, *Sycon ciliatum*, *Hydra vulgaris*, *Nematostella vectensis*, *Spongilla lacustris*, *Xenia sp.*, *Trichoplax adhaerens*, *Arabidopsis thaliana*, *Capsaspora owczarzaki*, *Chlamydomonas reinhardtii*, *Dictyostelium discoideum*, *Monosiga brevicollis*, *Saccharomyces cerevisiae*, *Salpingoeca rosetta*, *Ustilago maydis*, *Spizellomyces punctatus*). Protein sequences were queried using the BLOSUM62 (*79*) substitution matrix with an E-value cutoff of 1e-6 for BLAST. PROST searches were conducted using the same E-value threshold. For decarboxylases, we further included proteomes prokaryotic species that possess decarboxylases as part of human gut microbiota. After performing BLAST and PROST, a sequence alignment of all the hits with query sequences was generated using MAFFT (*80*) with the L-INS-i strategy (local pairwise alignment with iterative refinement) and 1,000 refinement iterations. For decarboxylases, additional annotated sequences that were not discovered by PROST and BLAST were added from decarboxylase enzyme categories (4.1.1.15, glutamate decarboxylase; 4.1.1.22, histidine decarboxylase; 4.1.1.25, tyrosine decarboxylase; 4.1.1.28, aromatic-L-amino-acid decarboxylase; 4.1.1.105, L-tryptophan decarboxylase) in the Expasy Swiss Bioinformatics Resource Portal. For vesicular transporter, we downsampled by only selecting the top 100 hits from BLAST and PROST results separately for each organism due to many solute carrier paralogs amongst our proteomes for alignment. We also downsampled by selecting the overall top 30 hits from BLAST and the top 5 hits from PROST for each organism per query due to many GPCR paralogs. Sequence alignments were fed into iQTree2 (*67*) for generation of a phylogenetic tree with branch support assessed using 1,000 ultrafast bootstrap replicates and 1,000 SH-like approximate likelihood ratio test (aLRT) replicates. Phylogenetic trees were visualized in iTOL (*81*).

<u>Quantitative phosphoproteomics and sample preparation for MS</u>

6 X 100 *S. lacustris* gemmules were plated in 10 mL M-medium (in 60 mm culture dishes) and kept at 18°C in the dark. Plates were treated either for 3, 15, or 30 minutes with either 0.03% DMSO (control) or 37.5 µM tryptamine. The induction of behavioral response was monitored via brightfield microscopy. For all plates when reaching the desired post-treatment timepoint, the medium was discarded, and sponges were immediately flash-frozen by dipping the bottom of the culture dishes into liquid nitrogen. This ensured immediate stop of cellular function and ensured that measured phosphorylation events are solely attributed to the treatments. To prevent thawing and potential metabolic activity, the dishes were put on metal blocks cooled to about −80 °C by dry ice. A pre-chilled 16 G needle (BD Biosciences, 305198) was used to scrape off the sponges and transfer into pre-cooled 2 mL cryotubes. The samples were stored at −80 °C until further processing. The above sample preparation was repeated in three independent experiments to generate biological replicates.

Cell pellets were lysed in 500 µL of 4 M guanidinium isothiocyanate, 5 mM Tris(2-carboxyethyl)phosphine hydrochloride (TCEP), 20 mM 2-chloroacetamide (CAA), 1% N-lauroylsarcosine, 5% isoamyl alcohol, and 40% acetonitrile in 125 mM HEPES pH 8.5 and vortexed. To enhance lysis, bead beating was applied by adding 1.6mm beads and 3 cycles of 20 seconds 4m/s bead beating, followed by centrifugation for 10 minutes at 17000 x g. The resulting supernatant was transferred to a new tube, small aliquot was taken for protein determination by tryptophan assay (*82*) and the remaining supernatant was incubated for 1 hour at room temperature to ensure complete reduction and alkylation.  Proteins were precipitated by adding four volumes of ice-cold 100% acetonitrile. Protein pellets were collected by centrifugation at 17,000 × g for 5 minutes, washed twice with ice-cold 100% acetonitrile, followed by three additional washes with ice-cold 70% ethanol. The protein pellets were digested with TPCK-treated trypsin (Thermo Fisher Scientific) at a ratio of 1:50 (w/w) at 37°C in 50 mM TEAB pH 8.5 overnight. Post-digestion, peptide concentrations were determined using a tryptophan assay (*82*).

Phosphopeptide enrichment was performed essentially as previously described (*83*). Briefly, peptides were resuspended in IMAC loading solvent (80% acetonitrile, 0.4% TFA). Of each sample a small aliquot was used for full proteome analysis, while the remaining peptides were subjected to phosphopeptide enrichment using the KingFisher Apex™ platform (Thermo Fisher) and magnetic Fe-NTA beads (Cube Biotech). Enriched phosphopeptides were eluted with 0.2% dimethylamine in 80% acetonitrile to facilitate subsequent TMT labeling."

Up to 20 µg of peptides (after phosphopeptide enrichment and for lysate) were labeled using TMTpro™ 18plex reagent as previously described (*84*). Briefly, 0.5 mg of TMT reagent was dissolved in 45 µL of 100% acetonitrile. Subsequently, 8 µL of this solution was added to each peptide sample, followed by incubation at room temperature for 1 hour. The labeling reaction was quenched by adding 8 µL of a 5% aqueous hydroxylamine solution and incubating for an additional 15 minutes at room temperature. Labeled samples were then combined for multiplexing, desalted using an Oasis® HLB µElution Plate (Waters) according to the manufacturer's instructions, and dried by vacuum centrifugation.

Offline high-pH reversed-phase fractionation (*85*) was carried out using an Agilent 1200 Infinity high-performance liquid chromatography (HPLC) system, equipped with a Gemini C18 analytical column (3 µm particle size, 110 Å pore size, dimensions 100 x 1.0 mm, Phenomenex) and a Gemini C18 SecurityGuard pre-column cartridge (4 x 2.0 mm, Phenomenex). The mobile phases consisted of 20 mM ammonium formate adjusted to pH 10.0 (Buffer A) and 100% acetonitrile (Buffer B). The peptides were separated at a flow rate of 0.1 mL/min using the following linear gradient: 100% Buffer A for 2 minutes, ramping to 35% Buffer B over 59 minutes, increasing rapidly to 85% Buffer B within 1 minute, and holding at 85% Buffer B for an additional 15 minutes. Subsequently, the column was returned to 100% Buffer A and re-equilibrated for 13 minutes. During the LC separation, 48 fractions were collected. These were pooled into twelve fractions by combining every twelfth fraction. The pooled fractions were then dried using vacuum centrifugation.

An UltiMate 3000 RSLCnano LC system (Thermo Fisher Scientific) equipped with a trapping cartridge (µ-Precolumn C18 PepMap™ 100, 300 µm i.d. × 5 mm, 5 µm particle size, 100 Å pore size; Thermo Fisher Scientific) and an analytical column (nanoEase™ M/Z HSS T3, 75 µm i.d. × 250 mm, 1.8 µm particle size, 100 Å pore size; Waters) was used. Samples were trapped at a constant flow rate of 30 µL/min using 0.05% trifluoroacetic acid (TFA) in water for 6 minutes. After switching in-line with the analytical column, which was pre-equilibrated with solvent A (3% dimethyl sulfoxide [DMSO], 0.1% formic acid in water), the peptides were eluted at a constant flow rate of 0.3 µL/min using a gradient of increasing solvent B concentration (3% DMSO, 0.1% formic acid in acetonitrile).

Peptides were introduced into an Orbitrap Fusion™ Lumos™ Tribrid™ mass spectrometer (Thermo Fisher Scientific) via a Pico-Tip emitter (360 µm OD × 20 µm ID; 10 µm tip, CoAnn Technologies) using an applied spray voltage of 2.2 kV. The capillary temperature was maintained at 275 °C. Full MS scans were acquired in profile mode over an m/z range of 375–1,650, with a resolution of 120,000 at m/z 200 in the Orbitrap. The maximum injection time was set to 50 ms, and the AGC target limit was set to 'standard'. The instrument was operated in data-dependent acquisition (DDA) mode, with MS/MS scans acquired in the Orbitrap at a resolution of 30,000. The maximum injection time was set to 110 ms, with an AGC target of 200%. Fragmentation was performed using higher-energy collisional dissociation (HCD) with a normalized collision energy of 34%, and MS2 spectra were acquired in profile mode. The quadrupole isolation window was set to 0.7 m/z, and dynamic exclusion was enabled with a duration of 20 seconds. Only precursor ions with charge states 2–7 were selected for fragmentation.

<u>Quantitative transcriptomic and sample preparation</u>

6 X 50 *S. lacustris* gemmules were plated in 10 mL M-medium (in 60 mm culture dishes) and kept at 18°C in the dark. Plates were treated either for 1 or 4 hours with either 0.03% DMSO (control) or 37.5 µM tryptamine (4 plates). A recovery condition was also tested (2 plates) where plates were treated for 4 hours (DMSO or tryptamine) followed by a media change and recovery of 24 hours. The induction of behavioral response was monitored via brightfield microscopy. For all plates when reaching the desired post-treatment timepoint, the medium was discarded, and RNA extraction took place using the Qiagen AllPrep Micro kit as described above. Extracted RNA were stored at - 80 °C until further processing as described above in RNA sequencing. The above sample preparation was repeated in three independent experiments to generate biological replicates.

GPCR AlphaFold3 docking with monoamines.

The docking potential between monoamines and GPCR candidates was predicted and assessed using AlphaFold3 model (*47*) with custom ligands indicated by CCD codes (TSS for tryptamine, PEA for phenethylamine, and AEF for tyramine). Every GPCR candidate was docked with three monoamines. The criteria for favorable docking included an interface predicted template modeling (ipTM) score above 0.9 (Fig. 5B, Fig. S20 B to D, Fig. S21 A to B)

*S. lacustris* mRNA injection, electroporation, and imaging.

*S. lacustris* gemmules were plated in 4 mL M-medium (in 35 mm glass-bottom culture dishes) and kept at 18°C in the dark. Twenty-four hours before desired tryptamine treatment experiment, sponge tissue were injected with ~ 10 pL of 1 mg/mL Dasher-GFP or mCherry with Picospritzer II (Parker NA) followed by immediate electroporation with Nepagene Super Electroporator NEPA21 Type II (Bulldog Bio) with the following parameters: Poring Pulse voltage at 120 volts, pulse length of 5.0 milliseconds, pulse interval of 50.0 milliseconds, 2 pulses, 10% decay rate, +/- polarity, and current limit turned on. Transfer Pulse voltage at 20 volts, pulse length of 50.0 milliseconds, pulse interval of 50.0 milliseconds, 5 pulses, 40% decay rate, and +/- polarity. Injected sponges were kept at 18°C in the dark to recover for at least 24 hours. Fluorescently label sponges were imaged using a spinning disk confocal microscope (Nikon Ti2-E connected to Yokogawa W1) using a 60× oil objective (NA = 1.40) or a 40x multi-emersion objective (NA = 1.25). A lack of endogenous behavior was verified before the addition of 37.5 µM of tryptamine and sponges were imaged at 1 frame/20-30 seconds for ~30 minutes after treatment. Samples were visualized with DIC and a 488 nm laser excitation or a 561 nm laser excitation.

*S. lacustris* Immunofluorescence

*S. lacustris* gemmules were plated in 4 mL M-medium (in 35 mm glass-bottom culture dishes) and kept at 18°C in the dark. One hour before experimentation, M-medium of each plate was changed to 2.67 mL. After letting plates rest at room temperature in the dark for one hour, they were first treated with 0.03% DMSO, 5 µM gallein, or 20 µM Y-27632 (by addition of 1.33 mL of 3X concentrated stock solution). Five minutes after the first treatment, or 15 minutes in the case of gallein, sponges were treated with 2 mL of 0.1% DMSO or 112.5 µM tryptamine to reach a final concentration of 0.03% or 37.5 µM. The entire treatment process was monitored via brightfield microscopy to keep track of behavioral responses. For all plates reaching ~ 8 minutes post addition of the second treatment (DMSO or tryptamine), the medium was discarded, and sponges were immediately fixed in 2 mL immunofluorescence fixative solution (45% ethanol, 4% methanol-free formaldehyde, and 51% M-medium) for 45 minutes in the dark at room temperature. Fixation was quenched with 2 mL 0.2 M glycine (Sigma, G7126) in PBST for 10 minutes. After three washes with PBST, sponges were blocked in 3% bovine serum albumin (Sigma; A9647) in PBST for 45 minutes. Once blocking solution is removed, sponges were incubated with 100 µL of primary antibody solution (Anti-α-tubulin, 1:100, DSHB 12G10; Anti-EmVinculin, 1:500, a gift from Scott Nichols (*33*); Anti-phospho-myosin light chain S19, 1:50, Cell Signaling Technology, 3671S in blocking buffer) for overnight at 4 °C in the dark. Primary antibodies were washed three times in PBST before the addition of 100 µL of secondary antibody solution (anti-mouse IgG 488, 1:500, Jackson ImmunoResearch, 115-545-044; anti-rabbit IgG 647, 1:500, Jackson ImmunoResearch, 111-605-003) supplemented with 2X phalloidin Alexa Fluor 546 (Thermo, A12380) and 1 ng/µL DAPI. After 45 minutes of secondary antibody incubation in the dark, sponges were washed three times in PBST and imaged immediately on a Zeiss LSM980 confocal microscope in PBS. Samples were imaged with a 40 X water immersion objective (NA=1.2) as Z-stacks in multiplex CO-8Y Aryscan 2 mode and later underwent Aryscan processing with default parameters.

*C. elegans* behavior statistical analysis

For tyramine-dependent touch reversal, the number of reversals for wildtype (N2), mutant, and extrachromosomal array strains were plotted using Prism 9 (GraphPad). For dopamine-dependent slowing on food, the basal slow rate for wildtype (N2), mutant, and extrachromosomal array strains were plotted using Prism 9 (GraphPad). Statistical significance was determined using a nonmatching analysis of variance (ANOVA) test followed by a Šídák multiple-comparison test with a 95% confidence interval ($\alpha = 0.05$).

Immunofluorescence quantification and statistical analysis

As part of *C. elegans* anti-serotonin immunofluorescence, the number of serotonin positive neurons near the pharynx were counted for each worm. Wildtype (N2) worms are expected to have 7 serotonin positive neurons (2 NSMs, 2 ADFs, 1 RIH, and 2 AIMs, Fig. 3C). For rescuing decarboxylases (serotonin synthesis) in the *bas-1 (tm351)* mutant background, we counted the total number of serotonin positive neurons. For rescuing vesicular transporters (serotonin uptake) in the *cat-1 (ok411)* background, we counted the total number of serotonin positive neurons subtracted by 2 NSMs. Even though RIH and AIMs are serotonin positive by up-taking secreted serotonin, it has been observed that ADFs have significant weaker staining in the *cat-1 (ok411)* background (*23*). Number of serotonin positive cells were plotted using Prism 9 (GraphPad). Statistical significance was determined using a nonmatching analysis of variance (ANOVA) test followed by a Šídák multiple-comparison test with a 95% confidence interval ($\alpha = 0.05$).

For quantification of number of vinculin foci in *S. lacustris* outer tent, image analysis was performed using a custom macro in ImageJ/Fiji. Z-stack maximum intensity projections were split into separate channels for processing. For nuclei quantification (DAPI), images were converted to 8-bit grayscale, smoothed using a Gaussian blur ($\sigma = 5.0$), and background-corrected using a rolling ball algorithm (radius = 100 pixels). Nuclei were segmented using the "Default" auto-thresholding method, followed by a watershed algorithm to separate touching cells. Objects were counted using the Analyze Particles command, filtering for sizes > 10 units$^2$

and circularity between 0.10 and 1.00. For vinculin quantification, a similar pipeline was employed with modified parameters to account for punctate morphology. Images were smoothed ($\sigma = 2.0$) and background-corrected (rolling ball radius = 10 pixels). Following "Default" auto-thresholding and watershed separation, particles larger than 0.5 $\mu m^2$ were counted. The number of vinculin foci was divided by the number of nuclei to normalize different amount of cells across regions of interests and plotted using Prism 9 (GraphPad). Statistical significance was determined using a nonmatching analysis of variance (ANOVA) test followed by a Šídák multiple-comparison test with a 95% confidence interval ($\alpha = 0.05$).

Vinculin focal adhesions were quantified using a custom macro in ImageJ/Fiji. Following channel splitting, the Vinculin channel was converted to 8-bit grayscale. To enhance feature detection, images were smoothed using a Gaussian blur ($\sigma = 2.0$) and processed with a rolling ball background subtraction (radius = 10 pixels). Focal adhesions were segmented using the "Default" auto-thresholding algorithm and separated using a watershed transform. Individual adhesions with an area greater than 0.5 $\mu m^2$ were analyzed to extract surface area and mean fluorescence intensity. Vinculin foci areas were plotted using Prism 9 (GraphPad). Statistical significance was determined using nonmatching, non-parametric Kruskal-Wallis test followed by a Dunn's multiple-comparison test with a 95% confidence interval ($\alpha = 0.05$).

Phospho-myosin light chain (pMYL) mean fluorescent intensity (MFI) were quantified using a custom ImageJ/Fiji macro. The F-actin channel served as a structural marker to define locations of contractility. F-actin images were smoothed (Gaussian blur, $\sigma = 2.0$) and segmented using the "Percentile" auto-thresholding algorithm to generate binary masks. These masks defined region of interests for objects > 1.0 $\mu m^2$. The defined ROIs were then applied to the corresponding pMYL channel to quantify MFI. MFI were plotted using Prism 9 (GraphPad). Statistical significance was determined using a nonmatching analysis of variance (ANOVA) test followed by a Šídák multiple-comparison test with a 95% confidence interval ($\alpha = 0.05$).

Metabolomic data processing, quantification, and statistical analysis

For each 13C enriched metabolites found by CompoundDiscoverer 3.2 (Thermo Fisher Scientific), the relative exchange rate of each 13C isotopologue was calculated by dividing the relative peak area of that isotopologue with the sum of all relative peak areas of all isotopologues of the same metabolite. The relative exchange rate of different isotopologues under the six different feeding conditions were plotted using Prism 9 (GraphPad).

Phosphoproteomics data processing, quantification, and statistical analysis

Raw files were converted to mzML format using MSConvert from ProteoWizard, using peak picking, 64-bit encoding and zlib compression, and filtering for the 1000 most intense peaks. Files were then searched using MSFragger in FragPipe (22.1-build02) against FASTA database spongilla_lacustris_180403 containing common contaminants and reversed sequences. The following modifications were included into the search parameters: Carbamidomethylation (C, 57.0215), TMTpro (K, 304.2072) as fixed modifications; Oxidation (M, 15.9949), Acetylation (protein N-terminus, 42.0106), TMTpro (peptide N-terminus, 304.2072), Phosphorylation (STY, 79.9663, only for phospho-enriched samples) as variable modifications. For the full scan (MS1) a mass error tolerance of 20 PPM and for MS/MS (MS2) spectra of 20 PPM was set. For protein digestion, 'trypsin' was used as protease with an allowance of maximum 3 missed cleavages requiring a minimum peptide length of 7 amino acids. The false discovery rate on peptide and protein level was set to 0.01. The standard settings of the FragPipe workflow 'Default' were used. The following modifications were made:

```
msfragger.allowed_missed_cleavage_1: 3, msfragger.allowed_missed_cleavage_2: 3, msfragger.isotope_error: -1/0/1/2/3, msfragger.labile_search_mode: labile, msfragger.localize_delta_mass: true, msfragger.mass_diff_to_variable_mod: 1, msfragger.mass_offsets: 0/79.966331, msfragger.max_variable_mods_per_peptide: 5, msfragger.minimum_ratio: 0.00, msfragger.misc.fragger.clear-mz-hi: 134.5, msfragger.misc.fragger.clear-mz-lo: 125.5, msfragger.misc.fragger.digest-mass-lo: 200, msfragger.misc.fragger.enzyme-dropdown-1: trypsin, msfragger.misc.fragger.precursor-charge-hi: 6, msfragger.remainder_fragment_masses: -18.01056 79.966331 79.966331, msfragger.restrict_deltamass_to: STY, msfragger.search_enzyme_name_1: trypsin, msfragger.search_enzyme_nocut_1: P, msfragger.table.fix-mods.K.lysine.: 304.2072, msfragger.table.var-mods.peptide N-terminus.304.2072: peptide N-terminus, msfragger.table.var-mods.STY.79.9663: STY, msfragger.use_topN_peaks: 300, phi-report.filter: --sequential --picked --prot 0.01 --mapmods, phi-report.prot-level-summary: false, protein-prophet.cmd-opts: --maxppmdiff 2000000 --minprob 0.5, ptmprophet.cmdline: --keepold --static --em 1 --nions b --mods M\:15.9949,n-terminus\:42.0106,-terminus\:304.2072,STY\:79.966331 --minprob 0.5, ptmprophet.run-ptmprophet: true, tmtintegrator.add_Ref: 1, tmtintegrator.channel_num: TMT-18, tmtintegrator.extraction_tool: Philosopher, tmtintegrator.max_pep_prob_thres: 0.9, tmtintegrator.min_pep_prob: 0.5, tmtintegrator.min_percent: 0.025, tmtintegrator.min_site_prob: 0.75, tmtintegrator.mod_tag: S(79.9663),T(79.9663),Y(79.9663), tmtintegrator.run-tmtintegrator: true.
```

The raw output files of FragPipe (psm.tsv for phospho data and protein.tsv files for input data, (*86*)) were processed using the R programming language (ISBN 3-900051-07-0). Only peptide spectral matches (PSMs) with a phosphorylation probability greater 0.5 and proteins with at least 2 razor peptides were considered for the analysis. Phosphorylated amino acids were marked with a * in the amino acid sequences behind the phosphorylated amino acid, labeled with a 1, 2 or 3 for the number of phosphorylation sites in the peptide and concatenated with the protein ID in order to create a unique ID for each phosphopeptide. Raw TMT reporter ion intensities were summed for all PSMs with the same phosphopeptide ID. For the input data, the reporter ion intensities were used as given in the protein.tsv output files. Phospho signals were not influenced significantly by input abundance; thus, raw phosphor intensities were used to maximize coverage. In order to correct for technical variability, batch effects were removed using the 'removeBatchEffect' function of the limma package (*87*) on the log2 transformed summed TMT reporter ion intensities. Subsequently, normalization was performed using the 'normalizeVSN' function of the limma package (VSN - variance stabilization normalization - (*88*)). Missing values were imputed with the 'knn' method using the 'impute' function from the Msnbase package (*89*).This method estimates missing data points based on similarity to neighboring data points, ensuring that incomplete data did not distort the analysis. Differential expression analysis was performed using the moderated t-test provided by the limma package (*87*). The model accounted for replicate information by including it as a factor in the design matrix passed to the 'lmFit' function. Imputed values were assigned a weight of 0.01 in the model, while quantified values were given a weight of 1, ensuring that the statistical analysis reflected the uncertainty in imputed data. To obtain p-values and false discovery rates (FDRs), the 'fdrtool' function from the fdrtool package (*90*) was used to analyze the t-values produced by limma for certain comparisons. Proteins were annotated as hits if they had a false discovery rate (FDR) below 0.05 and an absolute fold change greater than 2. Proteins were considered candidates if they had an FDR below 0.2 and an absolute fold change greater than 1.5. Gene Ontology (GO) enrichment analysis was conducted using goseq (*91*) with accounting for protein length bias. Functional annotations were derived from PROST (*78*) annotated *S. lacustris* proteome. A fixed universe consisting of unique proteins detectable from phosphoproteomics with those possessing valid GO annotations was used as background while all unique hits across all conditions were aggregated for enrichment. Wallenius' non-central hypergeometric distribution was used to calculate over-representation p-values, followed by Benjamini-Hochberg (BH) correction for multiple testing. Terms with an adjusted p-value ($< 0.05$) were considered significantly enriched. To refine the biological interpretation, terms associated with broad or potentially circular keywords—including "phosphorylation," "kinase activity," "phosphotransferase," "kinase," and "signal transduction"—were excluded from the visualization. The analysis focused specifically on Molecular Function (MF) and Cellular Component (CC) ontologies. For each ontology, the top six most significant terms, ranked by their p-values, were selected for final plotting.

Transcriptomic data processing, quantification, and statistical analysis

FASTQ files from Illumina pair-end reads of six treatment conditions in triplicates were verified to be sufficient quality with FastQC. Raw paired-end sequencing reads were quality controlled and preprocessed using fastp (Chen et al., 2018). The software was configured to automatically detect and remove adapter sequences (--detect_adapter_for_pe) and trim poly-G tails (--trim_poly_g) generated by two-color sequencing chemistry. The reference index was built using STAR (Dobin et al., 2013) with the *S. lacustris de novo* transcriptome FASTA and annotation files, optimized for 150 bp reads (--sjdbOverhang 149) and a scaled suffix array index (--genomeSAindexNbases 11). Paired-end reads were aligned to the reference using STAR, outputting coordinate-sorted BAM files. Gene expression was subsequently quantified using featureCounts (Liao et al., 2014). Counting was performed in paired-end mode (-p), requiring both ends to be successfully mapped (-B) and excluding chimeric fragments (-C).

Raw gene counts generated by featureCounts were processed in R, Median gene lengths were calculated from the raw count matrix by computing the median of exon lengths provided in the End column. Transcripts Per Million (TPM) were calculated to estimate relative abundance using these median lengths, normalizing for sequencing depth and gene length. Differential expression analysis was performed using DESeq2 (*92*). A grouped design matrix (~ 0 + Group) was constructed by combining treatments (DMSO, tryptamine) and timepoint (1h, 4h, 24h recovery) into a single factor. This approach allowed for specific contrasts without an intercept. Results were filtered using a Benjamini-Hochberg adjusted p-value cutoff of $< 0.05$. Genes were annotated as hits if they had an absolute fold change greater than 4. Genes were considered candidates if they had an absolute fold change greater than 1.5.

Gene Ontology (GO) enrichment analysis was conducted using the goseq (*91*) package to account for gene length bias. Functional annotations were derived from EggNOG-mapper. Enrichment was performed separately for genes identified as significant at the 4-hour treatment and 24-hour recovery time points against a background of all transcripts detected with GO annotations. Statistical significance for over-representation was determined using Wallenius' non-central hypergeometric distribution followed by Benjamini-Hochberg correction, adjusted p-value (< 0.05). To improve biological interpretability and reduce semantic redundancy, a refined deduplication logic was applied to the Biological Process (BP) ontology. Directional variants were collapsed into unified biological modules by stripping prefixes such as "positive regulation of," "negative regulation of," or "regulation of". Only top five non-redundant Biological Process terms for the 4-hour and recovery time points were visualized.

**Table S1 List of *C. elegans* strains.**

| *C. elegans* Strain | Genotype | Source and/or parent strains[a] | Relevant Figures |
|---|---|---|---|
| N2 | N2 (Bristol) WT | CGC | 3a-c |
| RB993 | *tdc-1(ok914)* II | CGC | 3a |
| RB681 | *cat-1(ok411)* X | CGC | 3a-c, S10 |
| LC33 | *bas-1(tm351)* III | CGC | 3b-c, S10 |
| MOY0155 | *cat-1 (ok411)* X; moaEx85 [*cat-1p::sl_c78726g1_ SL2-mCherry::unc-54 UTR + rps-0p::HygR + myo-2p::GFP::unc-54 UTR*] line 1 | Injected *sl_c78726-g1* into RB681 | 3a-c, S10 |
| MOY0156 | *cat-1 (ok411)* X; moaEx86 [*cat-1p::sl_c78726g1_SL2-mCherry::unc-54 UTR + rps-0p::HygR + myo-2p::GFP::unc-54 UTR*] line 2 | Injected *sl_c78726-g1* into RB681 | 3a-c, S10 |
| MOY0157 | *cat-1 (ok411)* X; moaEx87 [*cat-1p::sl_c93846g1_SL2-mCherry::unc-54 UTR + rps-0p::HygR + myo-2p::GFP::unc-54 UTR*] line1 | Injected *sl_ c93846-g1* into RB681 | 3a-c, S10 |
| MOY0158 | *cat-1 (ok411)* X; moaEx88 [*cat-1p::sl_c93846g1_SL2-mCherry::unc-54 UTR + rps-0p::HygR + myo-2p::GFP::unc-54 UTR*] line 2 | Injected *sl_ c93846-g1* into RB681 | 3a-c, S10 |
| MOY0159 | *cat-1 (ok411)* X; moaEx89 [*cat-1p::sl_c99657g1_SL2-mCherry::unc-54 UTR + rps-0p::HygR + myo-2p::GFP::unc-54 UTR*] line 1 | Injected *sl_ c99657-g1* into RB681 | 3a-c, S10 |
| MOY0161 | *cat-1 (ok411)* X; moaEx91 [*cat-1p::sl_c100403g1_SL2-mCherry::unc-54 UTR + rps-0p::HygR + myo-2p::GFP::unc-54 UTR*] line 1 | Injected *sl_ c100403-g1* into RB681 | 3a-c, S10 |
| MOY0162 | *cat-1 (ok411)* X; moaEx92 [*cat-1p::sl_c100403g1_SL2-mCherry::unc-54 UTR + rps-0p::HygR + myo-2p::GFP::unc-54*] line 2 | Injected *sl_ c100403-g1* into RB681 | 3a-c, S10 |
| MOY0163 | *cat-1 (ok411)* X; moaEx93 [*cat-1p::sl_c102470g2_SL2-mCherry::unc-54 UTR + rps-0p::HygR + myo-2p::GFP::unc-54 UTR*] line 1 | Injected *c102470-g2* into RB681 | 3a-c, S10 |
| MOY0164 | *cat-1 (ok411)* X; moaEx94 [*cat-1p::sl_c102470g2_SL2-mCherry::unc-54 UTR + rps-0p::HygR + myo-2p::GFP::unc-54 UTR* ] line 2 | Injected *c102470-g2* into RB681 | 3a-c, S10 |
| MOY0165 | *cat-1 (ok411)* X; moaEx95 [*cat-1p::Ce_cat1-SL2-mCherry::unc-54 UTR + rps-0p::HygR + myo-2p::GFP::unc-54 UTR*] line 1 | Injected *C. elegans cat-1* into RB681 | 3a-c, S10 |
| MOY0166 | *bas-1(tm351)* III; moaEx96 [*cat-1p::sl_c101057g1_SL2-mCherry::unc-54 UTR + rps-0p::HygR + myo-2p::GFP::unc-54 UTR*] line 1 | Injected *c101057-g1* into LC33 | 3b-c, S10 |
| MOY0167 | *bas-1(tm351)* III; moaEx97 [*cat-1p::sl_c102964g1_ SL2-mCherry::unc-54 UTR + rps-0p::HygR + myo-2p::GFP::unc-54 UTR*] line 1 | Injected *c102964-g1* into LC33 | 3b-c, S10 |

| | | | |
|---|---|---|---|
| MOY0168 | *bas-1(tm351)* III; moaEx98 [*cat-1p::Ce_bas1-SL2-mCherry::unc-54 UTR + rps-0p::HygR + myo-2p::GFP::unc-54 UTR*] line 1 | Injected *C. elegans bas-1* into LC33 | 3b-c, S10 |
| MOY0169 | *bas-1(tm351)* III; moaEx99 [*cat-1p::Ce_bas1-SL2-mCherry::unc-54 UTR + rps-0p::HygR + myo-2p::GFP::unc-54 UTR*] line 2 | Injected *C. elegans bas-1* into LC33 | 3b-c, S10 |
| MOY0170 | *bas-1(tm351)* III; moaEx100 [*cat-1p::Ce_tdc1-SL2-mCherry::unc-54 UTR + rps-0p::HygR + myo-2p::GFP::unc-54 UTR*] line 1 | Injected *C. elegans tdc-1* into LC33 | 3b-c, S10 |
| MOY0171 | *tdc-1(ok914*) II; moaEx101 [*cat-1p::sl_c101057g1_SL2-mCherry::unc-54 UTR + rps-0p::HygR + myo-2p::GFP::unc-54 UTR*] line 1 | Injected *c101057-g1* into RB993 | 3a |
| MOY0172 | *tdc-1(ok914)* II; moaEx102 [*cat-1p::sl_c102964g1_SL2-mCherry::unc-54 UTR + rps-0p::HygR + myo-2p::GFP::unc-54 UTR*] line 1 | Injected *c102964-g1* into RB993 | 3a |
| MOY0173 | *tdc-1(ok914*) II; moaEx103 [*cat-1p::Ce_bas1-SL2-mCherry::unc-54 UTR + rps-0p::HygR + myo-2p::GFP::unc-54 UTR*] line 1 | Injected *C. elegans bas-1* into RB993 | 3a |
| MOY0174 | *tdc-1(ok914)* II; moaEx104 [*cat-1p::Ce_tdc1-SL2-mCherry::unc-54 UTR + rps-0p::HygR + myo-2p::GFP::unc-54 UTR*] line 1 | Injected *C. elegans tdc-1* into RB993 | 3a |
| MOY0244 | *cat-1 (ok411)* X; moaEx153[*cat-1p::c100626_g1_SL2-mCherry::unc-54 UTR; rps-0p::HygR; myo-2p::GFP::unc-54 UTR*] line 1 | Injected *c100626-g1* into RB681 | 3a-c, S10 |
| MOY0245 | *cat-1 (ok411)* X; moaEx154[*cat-1p::c100626_g1_SL2-mCherry::unc-54 UTR; rps-0p::HygR; myo-2p::GFP::unc-54 UTR*] line 2 | Injected *c100626-g1* into RB681 | 3a-c, S10 |
| MOY0246 | *cat-1 (ok411)* X; moaEx155[*cat-1p::c102790_g4_SL2-mCherry::unc-54 UTR; rps-0p::HygR; myo-2p::GFP::unc-54 UTR*] line 1 | Injected *c102790-g4* into RB681 | 3a-c, S10 |
| MOY0247 | *cat-1 (ok411)* X; moaEx156[*cat-1p::c102790_g4_SL2-mCherry::unc-54 UTR; rps-0p::HygR; myo-2p::GFP::unc-54 UTR*] line 2 | Injected *c102790-g4* into RB681 | 3a-c, S10 |
| MOY0248 | *cat-1 (ok411)* X; moaEx157[*cat-1p::c103357_g1_SL2-mCherry::unc-54 UTR; rps-0p::HygR; myo-2p::GFP::unc-54 UTR*] line 1 | Injected *c103357-g1* into RB681 | 3a-c, S10 |
| MOY0249 | *cat-1 (ok411)* X; moaEx158[*cat-1p::c103357_g1_SL2-mCherry::unc-54 UTR; rps-0p::HygR; myo-2p::GFP::unc-54 UTR*] line 2 | Injected *c103357-g1* into RB681 | 3a-c, S10 |

| Bacterial Strain | Species/Genotype | Source and/or parent strains | Relevant Figures |
|---|---|---|---|

| | | | |
|---|---|---|---|
| OP50 | *E. coli* / WT | Laboratory strain, CGC | 3a-c |

[a]CGC – *Caenorhabditis* Genetics Center; [b]NBRP – National BioResource Project

**Movie S1. OCM orthogonal view timelapse of *S. lacustris* treated with 37.5 µM tryptamine.**

**Movie S2. DIC EDF projection timelapse of *S. lacustris* during two consecutive endogenous deflation events.**

**Movie S3. OCM orthogonal view timelapse of *S. lacustris* treated with 37.5 µM phenethylamine.**

**Movie S4. Brightfield timelapse of *S. lacustris* treated with 0.03% DMSO or 50 µM tyramine. Graph on the right represents deflations over a 12-hour period.**

**Movie S5. Brightfield timelapse of *E. mulleri, E. fragilis,* and *H. pinaza* treated with indicated concentrations of tryptamine.**

**Movie S6. Brightfield timelapse of *E. mulleri, E. fragilis,* and *H. pinaza* treated with indicated concentrations of phenethylamine.**

**Movie S7. Brightfield timelapse of *E. mulleri* and *E. fragilis* treated with 0.03% DMSO or 50 µM tyramine.**

**Movie S8. DIC EDF projection timelapse of *S. lacustris* treated with 37.5 µM tryptamine. Left: a tall *S. lacustris.* Right: Tent area of a *S. lacustris*.**

**Movie S9. OCM orthogonal view timelapse of *S. lacustris* treated with 5 µM gallein and 37.5 µM tryptamine.**

**Movie S10. OCM orthogonal view timelapse of *S. lacustris* treated with 20 µM Y-27632 and 37.5 µM tryptamine.**

**Data S1. Untargeted heavy-isotope tracing metabolomics results for *S. lacustris* fed with heavy carbon-labeled amino acids. Tables include a list of metabolites in the standard compound library and detected metabolites with different relative abundance of heavy carbon incorporation.**

**Data S2. *S. lacustris* decarboxylases and transporters BLAST and PROST hits used for phylogenetic analysis.**

**Data S3. Quantitative phosphoproteomics results of tryptamine-treated *S. lacustris.***

**Data S4. Transcriptomics results of tryptamine-treated *S. lacustris***

**Data S5. *S. lacustris* GPCR, BLAST and PROST hits used for phylogenetic analysis.**
